# Multi-scale modeling of human tissues from spatial transcriptomics with TERRA

**DOI:** 10.64898/2026.07.29.741565

**Authors:** Sebastian Birk, Mohammad Vali Sanian, Amirhossein Vahidi, Samuel Ogden, Daniyal J. Jafree, Adib Miraki Feriz, Carlo Leonardi, Arpit Merchant, Zichen He, Lloyd Steele, Adam Boxall, Marta Rosa Sallese, Jonas Maaskola, Ciro Ramírez-Suástegui, Sevda Öğüt, Koen Rademaker, MS Vijayabaskar, Batuhan Cakir, Sergio Marco Salas, Valentina Lorenzi, Josephine M. Bryant, Cecilia Kyany’a, James O. Jones, Raheleh Rahbari, Arti Mala Raghubar, Grant D. Stewart, Benjamin Rumney, Catherine Tudor, Minal Patel, Jasmine Halliwell, Hon Man Chan, Tong Li, Heather Stanley, April Rose Foster, Fani Memi, Kenny Roberts, Andrew L. Trinh, Elizabeth Tuck, Tannia Gracia, Shreya Rai, David Adams, Simone Webb, Martin Prete, Roser Vento-Tormo, Mats Nilsson, Heiko Lickert, Fabian J. Theis, Amin Ardestani, James J. Sun, Foad J. Rouhani, Lassi Paavolainen, Menna R. Clatworthy, Omer Ali Bayraktar, Muzlifah Haniffa, Thomas Mitchell, Mostafa Bakhti, Mo Lotfollahi

## Abstract

Spatial transcriptomics maps gene expression at cellular resolution, revealing how cells organize into multicellular niches. Yet computational analyses remain dataset-specific, without a transferable representation of tissue organization that generalizes across datasets, tasks and tissues or predicts how tissues behave under perturbation. We present TERRA, a foundation model pretrained on 112 million human cells profiled by spatial transcriptomics. From a single pretrained backbone, TERRA yields embeddings at the scale of cells, the genes they express and the neighborhoods in which they reside, and supports spatial *in silico* perturbation, all applied zero-shot to unseen tissues. At the cell level, in newly generated spatial data for developing pancreas, TERRA identified an islet-associated capillary state which we posit represents a developmental precursor of the mature islet microvasculature. At the gene level, in untreated kidney sections, in silico knockout of immune-checkpoint targets predicted a gene program of immune-checkpoint-blockade-associated nephrotoxicity, which we validated in treatment-exposed tissue and recovered in blood. At the neighborhood level, TERRA mapped macrophages across tissues to identify recurring cross-organ niches, which we term archetypes, including a tumor-boundary niche associated with poor prognosis in kidney cancer. Together, TERRA captures the spatial and multicellular logic of human tissue and predicts, *in silico*, its response to perturbation, providing a multi-scale framework for tissue biology, therapeutic development and clinical application.

## Introduction

Human physiology and disease emerge from processes that operate across molecular, cellular and multicellular scales. Within a cell, genes act not in isolation but in coordinated gene programs that define cellular identity and state, shaped by and feeding back into the local tissue environment as cells communicate^1^ and organize into microenvironments that recur across tissues. Although single-cell RNA sequencing studies have contextualised how cell type-specific malfunctions contribute to many diseases, spatial studies of healthy and diseased human tissues have reinforced the importance of spatial organization of cells into multicellular niches governing homeostasis, disease progression and therapeutic response^2–4^, opening opportunities for molecular stratification, intervention and non-invasive disease monitoring^3,5^.

Spatial transcriptomics^5–7^ has made tissue organization directly measurable, mapping how cell types, transcriptional programs and cell–cell interactions contribute to disease in their native context. However, learning from these data at scale poses distinct challenges. Measurements are sparse and noisy, are generated across heterogeneous targeted gene panels that differ between studies and platforms, and remain comparatively scarce and costly to produce. As a result, most analyses are still dataset-specific: computational methods are fit to the experiment at hand for clustering, integration, spatial-domain detection or imputation, limiting the transfer of knowledge across tissues, studies and platforms, and introducing inconsistencies between datasets. Moreover, these methods remain largely descriptive, characterizing tissue organization as observed but unable to predict how it responds to perturbations such as genetic knockouts. Overcoming these limitations calls for artificial intelligence (AI) models that learn transferable, spatially aware representations across many datasets and can both represent tissue organization and predict its response to perturbation.

Foundation models have shown that large-scale self-supervised pretraining on single-cell RNA sequencing data can capture transferable biological structure, supporting tasks from cell-type annotation to perturbation prediction^8–13^. Yet independent benchmarks show that their zero-shot embeddings can be matched by simple baselines and that their perturbation predictions do not yet outperform linear models^14–17;^ and because they are trained on dissociated cells, they cannot observe the spatial context in which multicellular responses are organized. Spatial foundation models have begun to close this gap, using large-scale pretraining to transfer spatial context across datasets^18–21^, to integrate paired histology^22,23^, or to support generative simulation and perturbation^24^. Each, however, addresses only part of the problem. The models that transfer spatial context^18–21^ represent tissue at a single scale, as cell embeddings or as spatial domains, rather than jointly at gene, cell and neighborhood resolution, and their zero-shot representations often fail to generalize to unseen tissues or assays. Because these are aggregated embeddings that discard gene-level information, they remain largely descriptive, delineating niches or domains but unable to predict how perturbing a gene would reshape a cell and its neighborhood; image-omics^22,23^ and generative^24^ models add histology or simulation but likewise provide no transferable, multi-scale representation for zero-shot analysis across tissues. As a result, no existing model is at once broadly pretrained and zero-shot, multi-scale, and capable of in silico perturbation at gene resolution.

Here we present TERRA (Tissue Environment Relational Representation Architecture), a spatial foundation model built on two ideas that set it apart from prior spatial models: it tokenizes each cell together with its spatial neighborhood as a single sequence of gene tokens, so that one model yields representations at gene, cell and neighborhood scales while preserving gene identity rather than collapsing each cell to a single embedding; and it is pretrained with a joint-embedding predictive architecture (JEPA) that predicts masked molecular and spatial context in latent space rather than reconstructing counts. Pretrained on 112 million human cells from spatial transcriptomics, TERRA’s gene-resolved, context-aware design unifies three capabilities in a single framework: zero-shot generalization, so it applies directly to new datasets without retraining, labeled examples or task-specific fine-tuning; multi-scale representation, with embeddings that link genes to cells and to the tissue niches they form; and in silico perturbation, which edits individual gene tokens to predict how a cell and its surrounding niche respond when a gene is knocked out or its expression is altered, analogous to a CRISPR screen. Across benchmarks, TERRA resolved niches and cell types on previously unseen tissues and transferred across samples, donors, datasets, and assays. Applied to newly generated spatial data of human tissues, TERRA identified a developmental precursor of the mature islet microvasculature in pancreas, modeled tissue-level effects of long-term immune checkpoint blockade in the kidney, and mapped recurrent macrophage niche archetypes across nine adult organs. Emphasizing generalist over task-specific capabilities, TERRA moves spatial biology beyond dataset-specific description by providing a single model that resolves tissue organization across scales and predicts perturbation-induced responses.

## Results

### A multi-scale spatial foundation model for zero-shot tissue analysis

TERRA is a spatial foundation model for zero-shot analysis of imaging-based spatial transcriptomics: it can be applied directly to a new dataset, without retraining or labeled examples. As input it takes one or more tissue sections, using only single-cell gene expression and spatial coordinates. From a single pretrained model it then returns embeddings at three nested scales (genes, cells and cellular neighborhoods) that capture complementary biology (Fig. 1a). TERRA’s spatially aware tokenization represents each cell together with its local neighborhood and supports analysis both within and across cells around an index cell (Fig. 1b).

**Fig. 1.**
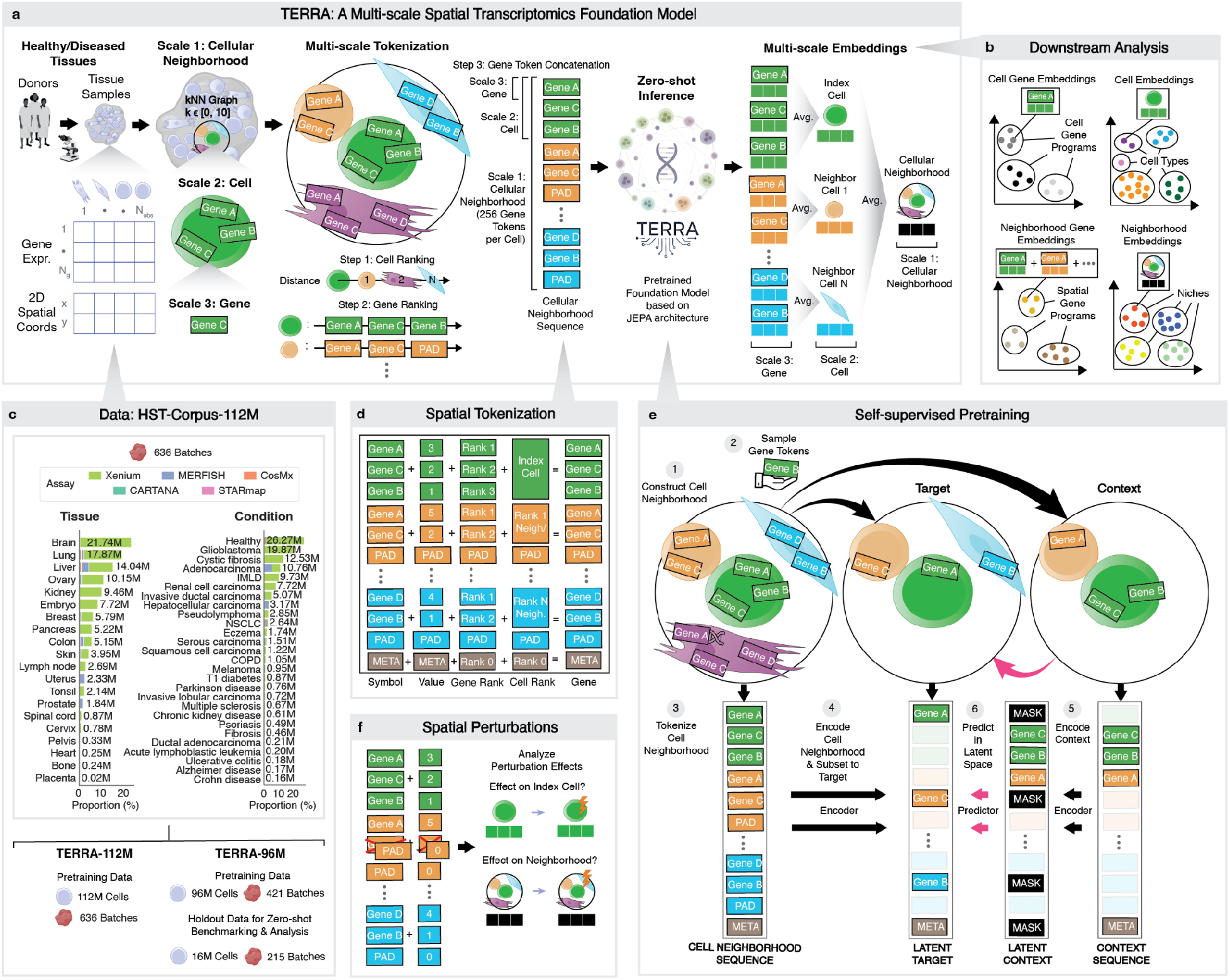
TERRA: a multi-scale spatial transcriptomics foundation model. **a**, TERRA is a self-supervised transformer foundation model for zero-shot analysis of imaging-based spatial transcriptomics data. Given a newly profiled tissue section, TERRA represents biology at three scales: cellular neighborhoods, cells, and genes. For each index cell, a cellular neighborhood is defined by the cell and its *k* nearest neighbors. The neighborhood is then tokenized into a sequence of gene tokens in three steps. First, cells are ranked by distance to the index cell. Second, genes within each cell are ranked by expression. Third, gene tokens from all cells in the neighborhood are concatenated. The resulting sequence is provided to TERRA without dataset-specific fine-tuning. TERRA outputs embeddings for individual gene tokens, which are aggregated to obtain multi-scale embeddings: cell gene embeddings (gene tokens from the index cell), neighborhood gene embeddings (tokens for a given gene averaged across the neighborhood), cell embeddings (average of all gene tokens from the index cell) and neighborhood embeddings (average of all gene tokens in the neighborhood). **b**, These embeddings support downstream analyses at multiple scales, including identification of cell(-intrinsic) gene programs, spatial gene programs, cell types and states, and niches from a single pretrained model. **c**, TERRA was pretrained on HST-Corpus-112M comprising 112 million single-cell-resolved spatial transcriptomic profiles from 636 tissue sections/batches generated with five imaging-based assays, spanning 20 tissues, and healthy tissue plus 26 disease conditions. For held-out evaluation, we additionally trained TERRA-96M on a 96-million-cell subset after withholding 215 sections/batches. IMLD, immune-mediated liver disease; NSCLC, non-small cell lung cancer; COPD, chronic obstructive pulmonary disease. **d**, TERRA uses a spatial tokenization scheme that combines count- and rank-based representations. Each gene token is represented by a gene symbol token, a value token encoding expression abundance, a gene-rank token denoting the expression rank of the gene within a cell, and a cell-rank token denoting the proximity rank of the cell within the neighborhood relative to the index cell. Optional metatokens encode neighborhood-level covariates, such as batch. **e**, TERRA is pretrained using a joint-embedding predictive architecture (JEPA). For each cellular neighborhood sequence, observed gene tokens are provided as context and masked gene tokens as targets. Context and target streams are encoded separately, and a predictor module learns to predict target embeddings from the context, enabling the model to capture spatial tissue organization and cellular interactions. **f**, Because TERRA operates at gene resolution within spatially defined cellular neighborhoods, the same framework can be used for spatial perturbation modeling. After perturbing selected gene tokens in specified cells, changes can be evaluated across gene, cell and neighborhood embeddings.

At its core, TERRA produces a contextual embedding for each gene in each cell that captures how that gene behaves both within its cell’s intrinsic state and within the surrounding cellular neighborhood. These per-gene embeddings are then aggregated into the higher scales. Isolating the gene embeddings of the index cell resolves cell-intrinsic gene programs (GPs) active within that cell, whereas aggregating a gene’s embeddings across neighboring cells yields intercellular gene embeddings that capture spatial gene programs (SGPs) operating across the local microenvironment. Averaging gene embeddings within the index cell produces a cell embedding that summarizes the state of that cell, supporting identification of cell types and states regardless of where a cell sits in the tissue. Lastly, aggregating all embeddings across the neighborhood produces a neighborhood embedding that captures variation in tissue microenvironments, which clustering resolves into tissue niches, the multicellular communities that define tissue organization. Because these multi-scale representations are produced by a single pretrained model, they can be applied directly to a newly generated dataset, without additional training, to resolve cell types and states, delineate tissue niches, identify cell-intrinsic and spatial gene programs, and perform in silico perturbation of genes in their native tissue context.

To span a broad range of human tissue biology in health and disease, we assembled Human Spatial Transcriptomics (HST)-Corpus-112M, a curated collection of 112 million single-cell-resolved spatial transcriptomic profiles from 636 tissue sections generated with five imaging-based assays (Xenium, MERFISH, CosMx, STARmap and ISS CARTANA) (Fig. 1c and Supplementary Fig. 1; Methods). The HST-Corpus-112M spans 20 human tissues, including healthy tissue, developmental tissue and 26 disease conditions encompassing malignant, autoimmune, inflammatory and degenerative pathologies. To separate model development from zero-shot evaluation, we trained two versions of the model (Fig. 1c): the TERRA-112M model was pretrained on the full HST-Corpus-112M and used for analyses across the complete collection, whereas the TERRA-96M model was pretrained after holding out 215 sections for benchmarking and analysis of unseen datasets (Methods).

TERRA uses a multi-part tokenization scheme tailored to targeted spatial transcriptomics data (Fig. 1d and Supplementary Fig. 2). Each gene token is represented by four components: a gene symbol token, a value token encoding expression abundance, a gene-rank token denoting the position of the gene within a cell, and a cell-rank token denoting the position of the cell within the neighborhood relative to the index cell. Optional metatokens encode neighborhood-level metadata. During pretraining, we included a batch metatoken to help account for assay- and sample-specific technical variation, including differences in gene panels, detection sensitivity, and noise characteristics.

Pretraining uses a joint-embedding predictive architecture (JEPA)^25^ in which a subset of gene tokens from the cellular neighborhood is retained as context and the remaining tokens are masked as targets (Fig. 1e). An encoder processes the context and target sequences separately, and a predictor maps the context representation to the latent representation of the target tokens (Methods). This objective encourages the model to recover stable latent structure rather than reconstruct every observed count directly, a bias that is well suited to spatial transcriptomics data, which are sparse and noisy. Batch metatokens are provided to both context and target streams during training to improve generalization to unseen datasets and padded during inference to facilitate integration (Methods).

We established TERRA’s tokenization and architecture through systematic ablations on a paired mouse-brain benchmark, in which MERFISH^26^ and STARmap^27^ profiled and matched coronal sections were annotated against the Allen Reference Atlas^28^. We varied the tokenizer, per-cell sequence length, expression and spatial encoding, count normalization and masking ratio (Supplementary Figs. 3–5 and Supplementary Notes 1–2). On the same benchmark, the batch metatoken enabled integration across assays without explicit batch correction (Supplementary Fig. 6), and TERRA outperformed both task-specific spatial methods and zero-shot single-cell and spatial foundation models at cross-assay niche identification while achieving strong native integration (Supplementary Figs. 7, 8 and Supplementary Note 3).

Finally, TERRA can move beyond descriptive analysis to predict how tissues respond to perturbation, because it operates at gene resolution within spatially defined cellular neighborhoods (Fig. 1f; Methods). A perturbation is specified by editing the gene token components of selected cells and re-encoding the neighborhood, so that predicted consequences can be generated for the index cell or its surrounding neighborhood. In this way, TERRA can simulate the knockout of specific genes, akin to a virtual CRISPR perturbation, and predict how the affected cells and their local tissue environment respond.

### TERRA generalizes across tissues, assays, patients, and tasks

We next evaluated how far TERRA’s pretrained neighborhood and cell embeddings can be used across downstream analyses spanning the neighborhood, cell and sample scales (Fig. 2 and Supplementary Fig. 9).

**Fig. 2.**
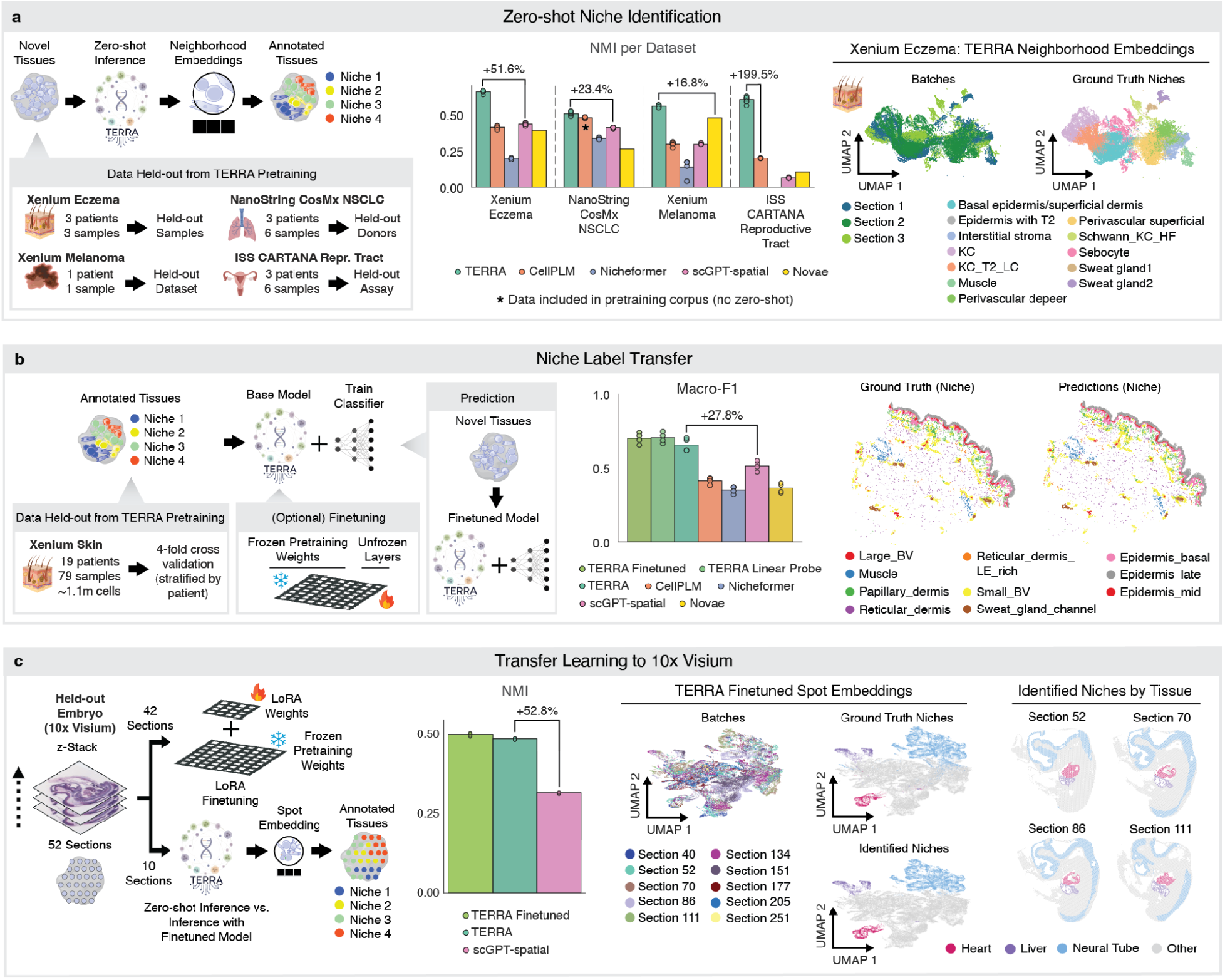
Zero-shot benchmarking of TERRA across tissues, assays and platforms. **a**, Zero-shot niche identification on four datasets held out from TERRA pretraining, spanning increasing levels of generalization: Xenium eczema skin (held-out samples), NanoString CosMx NSCLC (held-out donors), Xenium melanoma (held-out dataset) and ISS CARTANA healthy reproductive tract (held-out assay). Sections were tokenized and passed to the pretrained model without fine-tuning to obtain neighborhood embeddings, which were clustered and compared to ground-truth niches. Middle, NMI per dataset for TERRA and four spatial foundation models (CellPLM, Nicheformer, scGPT-spatial and Novae); bars show the mean and dots individual Leiden clustering resolutions. TERRA achieved the highest NMI on every dataset; brackets give TERRA’s relative improvement in mean NMI over the best-performing zero-shot baseline (scGPT-spatial for eczema and NSCLC, Novae for melanoma, CellPLM for the reproductive tract). The asterisk marks CellPLM on NanoString CosMx NSCLC, whose pretraining corpus included this dataset (not zero-shot); TERRA’s improvement there is therefore computed against scGPT-spatial. Nicheformer embeddings collapsed on the ISS CARTANA dataset. Right, UMAP of TERRA neighborhood embeddings for the three Xenium eczema sections, colored by section and by ground-truth niche, showing cross-section integration with clear niche separation. **b**, Niche label transfer on Xenium skin (19 patients, 79 samples, ∼1.1M cells), by 4-fold cross-validation stratified by patient. A classifier was trained on TERRA neighborhood embeddings with frozen weights (linear probe) or with selective unfreezing (fine-tuning) and applied to held-out patients. Middle, macro-F1 (averaged over niches) for TERRA fine-tuned, TERRA linear probe, TERRA (zero-shot), CellPLM, Nicheformer, scGPT-spatial and Novae (bars, mean; dots, folds); the bracket gives the improvement of zero-shot TERRA over scGPT-spatial. Right, a representative held-out section colored by ground-truth and predicted niche. **c**, Transfer learning to 10x Visium using a held-out embryo z-stack (52 serial sections; Methods). TERRA was adapted by low-rank adaptation (LoRA) on 42 sections with pretraining weights frozen, and evaluated on 10 held-out sections, comparing zero-shot to fine-tuned inference. Middle, NMI for TERRA fine-tuned, TERRA (zero-shot) and scGPT-spatial (bars, mean; dots, sections); the bracket gives the improvement of zero-shot TERRA over scGPT-spatial. Right, UMAP of TERRA fine-tuned spot embeddings colored by section, ground-truth niche and identified niche, with identified niches mapped onto four representative sections.

At the neighborhood scale, clustering TERRA neighborhood embeddings groups cellular neighborhoods into recurrent niches, so that niche maps can be built directly from the pretrained representation. To test this, we held out datasets from pretraining spanning increasing levels of generalization, from unseen samples to unseen donors, to an unseen dataset and finally an unseen assay (Fig. 2a, left). Benchmarking zero-shot niche identification against a panel of spatial foundation models, TERRA achieved the highest agreement with ground-truth niches across all held-out datasets, consistently outperforming the next-best method (Fig. 2a, middle, and Supplementary Fig. 9a). On Xenium eczema data, TERRA neighborhood embeddings integrated the tissue sections while cleanly separating the ground-truth niches (Fig. 2a, right).

Because TERRA places cellular neighborhoods from different samples and assays in a shared embedding space, niche annotations from one tissue can be transferred to new sections. On a large multi-patient skin dataset, TERRA transferred niche labels to held-out patients more accurately than competing spatial foundation models, whether applied zero-shot, as a linear probe on its fixed embeddings, or with selective fine-tuning (Fig. 2b and Supplementary Fig. 9b). Fine-tuning added little over the linear probe, indicating that the pretrained embeddings already capture niche structure without further weight updates, and the predicted niche maps closely matched the ground truth (Fig. 2b and Supplementary Fig. 9b).

TERRA can also be adapted to spatial transcriptomics technologies outside its imaging-based pretraining corpus, including sequencing-based platforms that measure expression at a coarser spatial scale. Using a held-out 10x Visium z-stack of a developing embryo (52 serial sections), where each spot captures multiple cells, we applied low-rank adaptation (LoRA), which keeps the pretrained weights fixed and trains only a small set of added low-rank parameters, on 42 sections and evaluated niche identification on 10 held-out sections (Fig. 2c and Supplementary Fig. 9c). LoRA fine-tuning improved over zero-shot inference, and both outperformed scGPT-spatial. The resulting spot embeddings recovered anatomically coherent niches that mapped onto expected embryonic structures, including heart, liver and neural tube (Fig. 2c).

The same principle applies at the cellular scale: clustering TERRA cell embeddings identifies cell types directly, without dataset-specific training. Across held-out datasets spanning unseen samples, donors, an unseen dataset and an unseen assay, TERRA identified cell types on par with the strongest spatial foundation models and well above weaker baselines (Supplementary Fig. 9d). Notably, TERRA matched these models even though its cell embeddings are a derived, aggregated output of a neighborhood-centered architecture, rather than the primary, cell-level representation the other models are explicitly optimized to produce. Finally, pooling cell embeddings into sample-level summaries enabled disease classification directly from spatial sections: TERRA distinguished eczema from psoriasis at both the patient and sample level, performing comparably to the best competing model and above the next-best method, drawing on signatures of cellular composition and microenvironmental organization (Supplementary Fig. 9e).

Consistent with foundation-model scaling, increasing model capacity improved zero-shot niche identification mainly at the harder hold-out levels, whereas performance on unseen samples was already near-saturated at the smallest model size; increasing the amount of pretraining data followed the same trend (Supplementary Fig. 10; Methods). Cell-type identification, by contrast, is saturated at a smaller model size, with no further gain from additional capacity (Supplementary Fig. 11).

Together, these results show that niche identification, niche label transfer, zero-shot cell-type identification and disease classification can all be performed directly from TERRA’s pretrained multi-scale embeddings on unseen data, and that the same embeddings support cross-platform transfer with only lightweight adaptation.

### TERRA resolves developmental niches and gene programs in the human pancreas

Organogenesis provides a context in which to test a spatial foundation model’s ability to resolve multicellular niches over time, as cell state, spatial organization and gene expression change rapidly over a developmental timeline. We thus evaluated TERRA’s ability to perform spatially informed biological discovery in a developing tissue not included in model training by generating a new spatial transcriptomic atlas of the developing human pancreas, a tissue that was held out from the TERRA-96M pretraining corpus. During pancreas development, multipotent pancreatic progenitors progressively segregate into distinct epithelial domains, including acinar-biased tips and ductal-endocrine-biased trunks, that ultimately give rise to the exocrine (acinar and ductal cells) and endocrine (islets of Langerhans) compartments (Supplementary Fig. 12a)^29,30^. Although previous studies have established that mesenchymal, endothelial, and neural compartments influence pancreatic epithelial growth and differentiation, the spatial organization, temporal dynamics, epithelial-domain-specific niches and how other cells contribute to functional maturation in the developing human pancreas remain poorly resolved^30,31^. To address this gap, we generated a new spatial single-cell transcriptomic atlas of human pancreatic development across seven developmental stages (8, 10, 11, 12, 13, 14, and 17 post-conceptional weeks, PCW) capturing the transition from early epithelial remodeling and patterning to the emergence of exocrine and endocrine lineages (Fig. 3a and Supplementary Fig. 12b). We then applied TERRA to systematically dissect cell types, multicellular niche organization, and cellular interactions across human pancreatic organogenesis. TERRA resolved 1,586,341 cells into the major pancreatic lineages, confirming that TERRA accurately resolves cell-type identity in a tissue context the model has never encountered (Fig. 3b,c and Supplementary Fig. 12c,d).

**Fig. 3.**
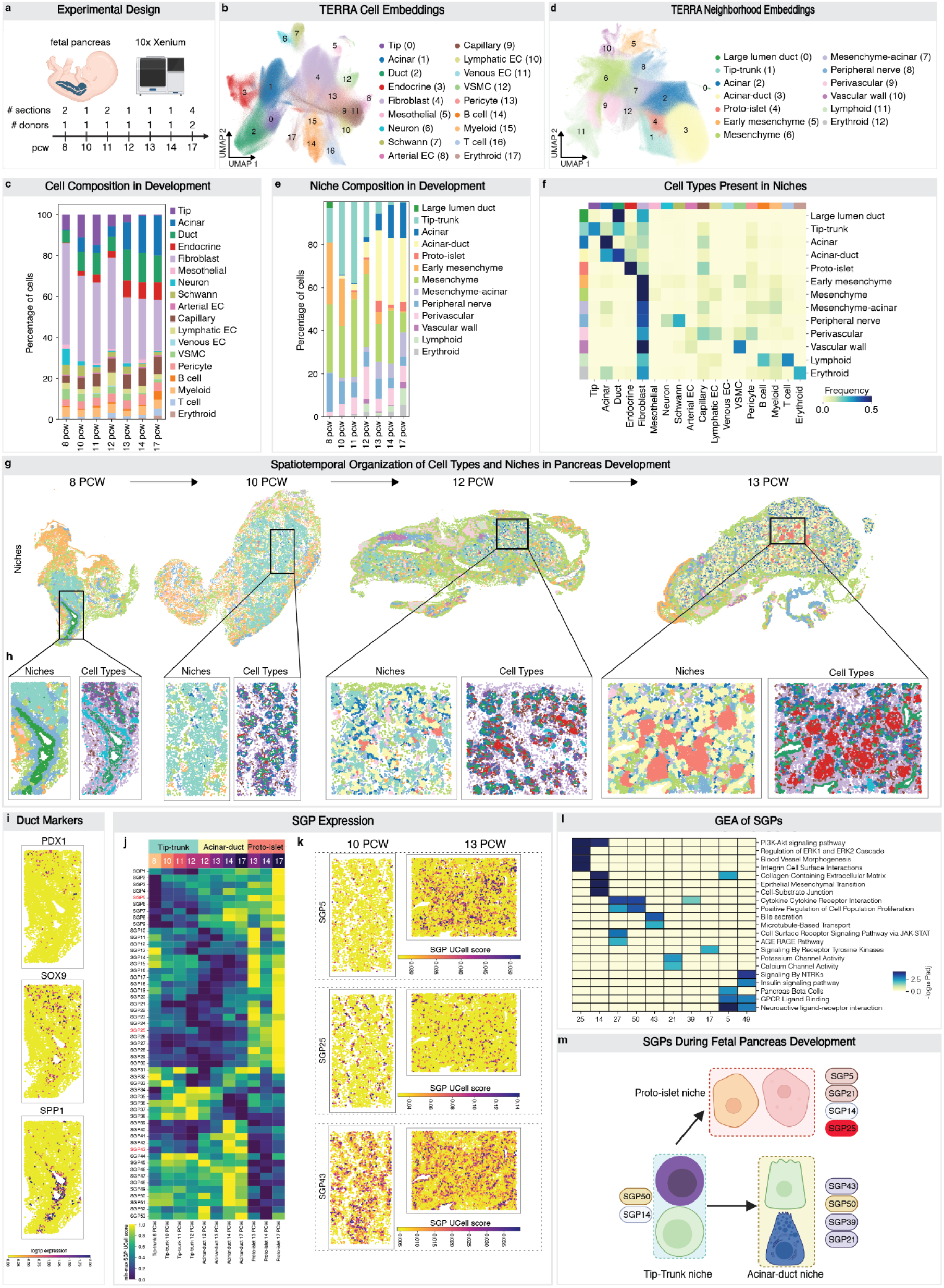
TERRA maps niches and gene programs across human pancreas development. **a**, Experimental outline and sampling strategy across fetal developmental stages. 12 sections from 7 developmental stages were profiled with Xenium 5K. **b**, UMAP of TERRA cell embeddings colored by cell-type annotations. Cell types were annotated using scVI. **c**, Cell-type composition across developmental stages. **d**, UMAP of TERRA neighborhood embeddings colored by niche annotations (13 niches). **e**, Niche composition across developmental stages. **f**, Heatmap of cell-type frequencies within niches. **g**, Spatial organization of niches in the indicated developmental stages. **h**, Magnified views of the regions shown in g, showing niche and cell-type annotations. **i**, Spatial expression of trunk-associated *PDX1* and duct-associated *SOX9* and *SPP1* at 8 PCW. **j**, Heatmap of min-max normalized SGP UCell scores for the 53 fetal pancreas SGPs across the indicated epithelial niches and developmental stages. Higher UCell scores indicate higher expression of genes comprising each SGP. SGPs shown in **k** are highlighted in red. **k**, Spatial projection of SGP expression in the indicated developmental stages. SGP5 - endocrine associated, SGP25 - endothelial associated, SGP43 - exocrine associated. **l**, Gene enrichment analysis of the indicated SGPs. **m**, Schematic summary of niche-associated SGP activation during fetal pancreas development. SGP, spatial gene program.

We next assessed whether TERRA could recover known multicellular niche architecture. Unsupervised clustering of TERRA neighborhood embeddings identified 13 multicellular niches (Fig. 3d and Supplementary Fig. 12e), which were manually annotated based on cell-type composition and spatial localization. We then mapped the distribution of these niches across pancreas development (Fig. 3e and Supplementary Fig. 12f). Focusing on epithelial-associated niches, TERRA captured the expected progression of pancreatic epithelial differentiation. The tip-trunk niche progressively declined, while acinar-ductal and acinar niches expanded, consistent with tip-to-acinar differentiation^32,33^. The proto-islet niche, primarily composed of endocrine cells, capillaries, and fibroblasts, emerged at 10 PCW and expanded at later stages, coinciding with endocrine differentiation from trunk progenitors and the progressive aggregation of endocrine cells during fetal islet morphogenesis^34^ (Fig. 3d-f and Supplementary Fig. 12e-g). Together, these results demonstrate that TERRA robustly resolves expected developmental niche architecture in an unseen tissue. In addition to this known developmental trajectory, TERRA identified a distinct epithelial niche characterized by a large lumen at 8 PCW, located outside the pancreatic epithelial compartment (tip-trunk niche) and embedded within the peripheral nerve niche (Fig. 3g,h). Given the close developmental and molecular relationship between biliary and pancreatic ductal systems, the presence of a large lumen led us to hypothesize that this niche represents extrahepatic biliary epithelium that arises from adjacent foregut endoderm^35,36^. To characterize the large-lumen duct (LLD) niche, we performed unbiased reclustering across developmental stages of epithelium from the LLD, tip-trunk and acinar-duct niches. This resolved six epithelial subclusters, with the LLD population forming a transcriptionally distinct cluster (cluster 5) that persisted beyond 8 PCW (Supplementary Fig. 13a-d). While cells within both LLD and trunk niches share ductal epithelial markers, including *SOX9*, at 8 PCW they displayed divergent transcriptional profiles (Supplementary Fig. 13e). Supporting their distinct transcriptional identity, LLD niche cells were enriched for *SPP1* and *TSPAN8* and showed very low or absent expression of pancreatic epithelial markers such as *PDX1*, *NKX6-1*, and *PTF1A*^37^ (Fig. 3i and Supplementary Fig. 13e-g). At later developmental stages, cluster 5 cells progressively upregulated canonical pancreatic ductal markers, and were found intermingled with pancreatic ductal epithelium (Supplementary Fig. 13g). Together, these findings suggest that the LLD niche represents a transcriptionally distinct ductal population at 8 PCW that progressively acquires pancreatic ductal identity, potentially reflecting the convergence of extrahepatic duct-derived epithelium into the forming pancreatic ductal network during early organogenesis^35,36^.

To complement our spatial analysis with gene-level information, we next applied TERRA to identify spatial gene programs (SGPs) across distinct pancreatic developing niches. Given the central role of the pancreatic epithelium in giving rise to the major pancreatic compartments, we computed neighborhood-gene embeddings only from cells residing within epithelial niches (LLD, tip-trunk, acinar-duct, acinar, mesenchyme-acinar, and proto-islets) across all development stages (see Methods). This yielded embeddings for 5,009 genes, which were clustered into 53 SGPs (Fig. 3j and Supplementary Table 1). We uncovered SGPs consistent with the established functions of the exocrine (e.g. SGP43) and endocrine pancreas (e.g. SGP5) (Fig. 3k). Accordingly, pathway enrichment analysis identified terms corresponding to the known functions of the cell types in which these SGPs were enriched (e.g. SGP5 - *Pancreas Beta Cells* and SGP25 - *Blood Vessel Morphogenesis*) (Fig. 3l and Supplementary Table 1). Notably, several SGPs enriched in the early tip-trunk niche persisted at later stages in either the acinar-duct (SGP50) or proto-islet (SGP14) niches, highlighting continued deployment of components of the early tip-trunk transcriptional program during niche maturation (Fig. 3j,m). Finally, the proto-islet niche showed progressive enrichment of SGP25 (including genes *PLVAP*, *NOTCH4*, *ESM1*, *ANGPT2*, *KDR*) over developmental time, corresponding to endothelial development and remodeling, indicative of active angiogenesis during islet cell differentiation and morphogenesis (Fig. 3j,k), consistent with the established role of endothelial cells (ECs) in supporting islet development^38^. These findings demonstrate that TERRA identifies SGPs that define niche identity and reveal their dynamic remodeling throughout pancreatic development.

### TERRA traces an islet-associated capillary state from development to adult pancreas

Several recent spatial transcriptomic studies have examined the mesenchymal niche during pancreas development^39–41^. However, we focused on interrogating EC heterogeneity and contribution to multicellular niches in the developing human pancreas given the central role of ECs in islet vascularization and development^42–45^. Although scRNA-seq studies have identified a distinct population of adult pancreatic islet ECs^46,47^, a systematic and spatially aware analysis of vascularization during human pancreas development is lacking. We therefore used TERRA to generate cell and neighborhood embeddings for 172,904 cells annotated as ECs within our human developing pancreas atlas. Clustering of TERRA cell embeddings identified 10 endothelial subpopulations, comprising two arterial (*GJA5, EFNB2*, *SOX17*), five capillary (*PLVAP*), two lymphatic (*NTS*, *PROX1*, *FLT4*), and one venous (*SELP*) cluster (Fig. 4a and Supplementary Fig. 14a). EC cells within epithelial niches (tip-trunk, acinar-duct, acinar, mesenchyme-acinar and proto-islets) grouped closely together in the neighborhood embedding-based UMAP (Fig. 4b). To further explore the relationship of the EC subpopulations and their local niches we examined the frequency of EC subpopulations within niches. Of the five capillary subpopulations identified using TERRA cell embeddings, three are present across multiple niches (termed general (G) capillaries, G1, G2, G3), while one subpopulation is associated with acinar niches (acinar-associated, AA) and one with proto-islets (islet-associated, IA) (Fig. 4c). The majority (53.4%) of IA capillaries reside within the proto-islet niche (Fig. 4d). Within this niche, IA capillaries progressively increase in relative abundance during development, becoming the predominant capillary subpopulation by 14 PCW (Supplementary Fig. 14b,c). Consistent with this, of all capillary subpopulations, IA capillaries displayed the closest spatial association with endocrine cells (Supplementary Fig. 14d-i). This suggests that the IA capillaries identified by TERRA correspond to the islet-associated ECs previously described in scRNA-seq studies of the human adult pancreas^46,47^.

**Fig. 4.**
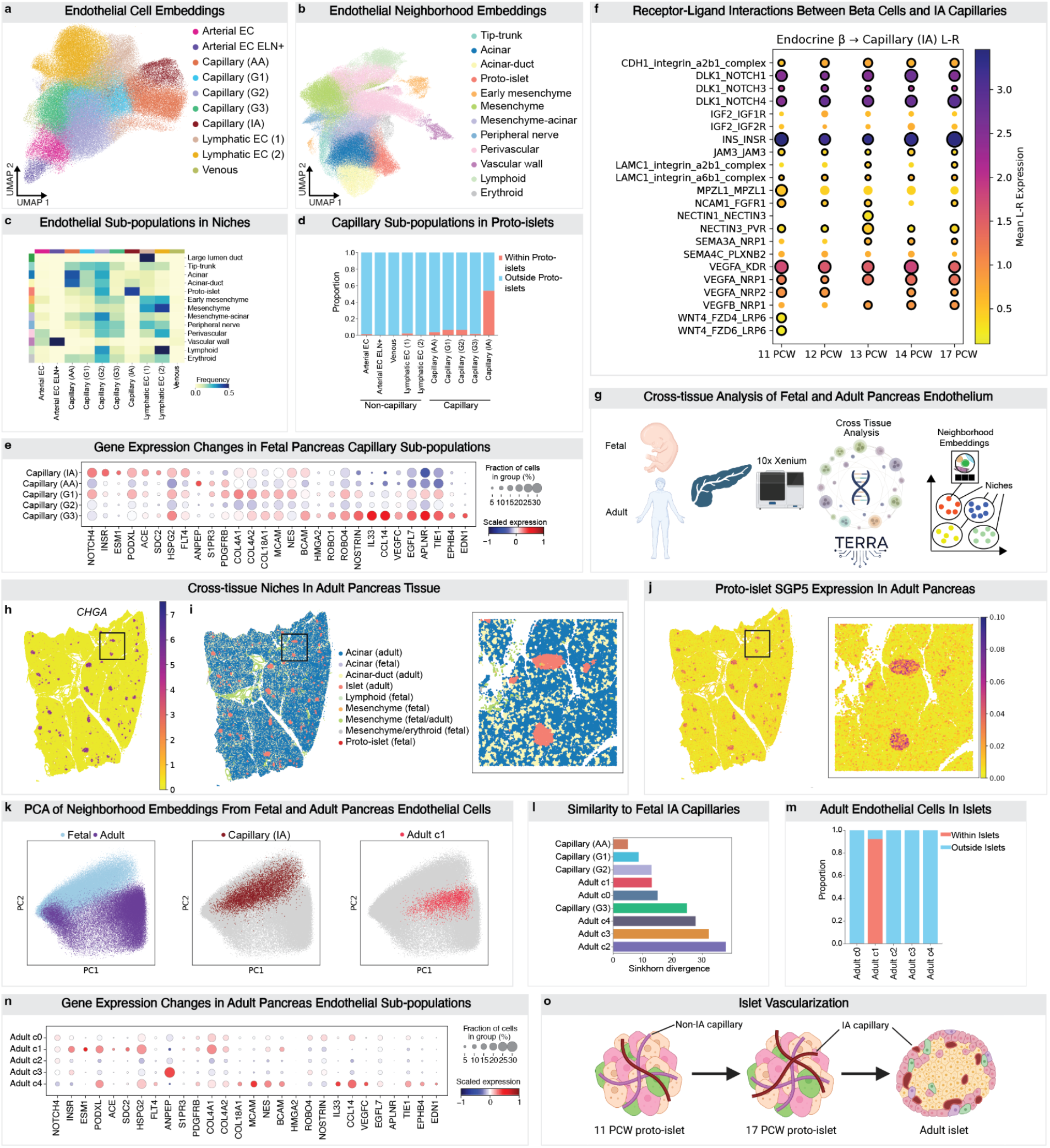
An islet-associated capillary state links developing and adult pancreas endothelium. **a**, UMAP of fetal pancreas endothelial TERRA cell embeddings colored by endothelial subpopulation annotation. **b**, UMAP of fetal pancreas endothelial TERRA neighborhood embeddings colored by niche annotation. **c**, Heatmap of endothelial subpopulation frequency across niches. **d**, Proportion of each endothelial subpopulation that is located within or outside the proto-islet niche. **e**, Expression of representative marker genes across fetal capillary subpopulations. **f**, Predicted ligand-receptor interactions between endocrine β cells (sender) and islet-associated capillaries (receiver) across developmental stages. Dot size indicates interaction score, color indicates mean ligand-receptor expression and black outlines denote statistically significant interactions (*P* < 0.05). **g**, Experimental design for the cross-tissue analysis of fetal and healthy adult pancreas tissue. Adult pancreas Xenium data were generated from 2 healthy donors, with 2 sections profiled per donor. **h**, *CHGA* expression (log1p normalized counts) in healthy adult pancreas. *CHGA* marks endocrine cells. **i**, Spatial organization of niches in healthy adult pancreas tissues. Niches were identified using cross-tissue TERRA neighborhood embeddings. **j**, Expression (UCell score) of fetal pancreas SGP5 (endocrine) in adult pancreas tissue. **k**, PCA of cross-tissue TERRA neighborhood embeddings of fetal and adult endothelial cells. **l**, Sinkhorn divergence in cross-tissue TERRA neighborhood embeddings space between fetal islet-associated capillaries and fetal subpopulations or adult endothelial clusters (Adult c0-4). Lower distances indicate greater similarities between the indicated subpopulation and fetal islet-associated capillaries. **m**, Barplot showing the proportion of each adult endothelial cluster located within or outside islets of Langerhans. **n**, Expression of the indicated marker genes in adult endothelial clusters. **o**, Proposed model summarizing endothelial niche remodeling during pancreas development.

To define the molecular signature of IA capillaries in the human developing pancreas, we performed differential gene-expression analysis across capillary clusters. IA capillaries displayed a distinct transcriptional profile, marked by elevated expression of genes including *NOTCH4*, *INSR*, *ESM1*, *ACE*, and *SDC2* (Fig. 4e and Supplementary Table 2). This transcriptional signature may be driven by the local proto-islet microenvironment rather than representing a stable subpopulation, as AA, G1, G2, and G3 capillary subpopulations adopted a similar gene-expression profile to IA capillaries when located within the proto-islet niche (Supplementary Fig. 14j,k). Consistent with this, expression of IA markers within G3 capillaries increases with closer proximity to endocrine cells (Supplementary Fig. 14l). Given the close spatial association between IA capillaries and endocrine cells within the proto-islet niche, we next performed ligand-receptor analysis to identify candidate signaling interactions between beta cells and IA capillaries across developmental stages (Supplementary Table 3; Methods). Focusing on beta cell-to-endothelial signaling, IA capillaries engaged the well-established VEGF endocrine-endothelial axis^48,49^ (*VEGFA-KDR*, *VEGFA*-*NRP1*, *VEGFA*-*NRP2*, *VEGFB*-*NRP1*) and additionally showed candidate interactions including *INS*-*INSR, SEMA4C-PLXNB2, NECTIN1-NECTIN3*, and *WNT4-FZD4/6-LRP6* (Fig. 4f). Together, these findings identify a proto-islet associated endothelial transcriptional program during human pancreas development.

To examine if TERRA could map the identified pancreatic EC subpopulations to their mature equivalents in the adult organ we performed single-cell spatial transcriptomics on four sections from two healthy adult human pancreas donors (Fig. 4g). TERRA neighborhood embeddings revealed four major niches in the adult pancreas, including acinar, acinar-duct, mesenchyme and islet (Fig. 4h,i and Supplementary Fig. 15a,b). TERRA successfully mapped several SGPs identified in the fetal pancreas to their adult counterparts, detecting high activity of proto-islet SGP5 in the adult islet niche and acinar SGP43 in the adult exocrine niche. The EC-related SGP25 likewise showed elevated activity within the adult islet niche (Fig. 4j and Supplementary Fig. 15c-e). In the adult organ, TERRA resolved five distinct endothelial clusters, of which cluster 1 accounted for approximately 90% of endothelial cells within the islet niche (Supplementary Fig. 15f,g). We next compared ECs across fetal and adult pancreas within a shared neighborhood embedding space. Among adult endothelial populations, cluster 1 bore the closest resemblance (shortest Sinkhorn distance) to fetal IA capillaries, while fetal G3 capillaries were similar to cluster 4 (Fig. 4k,l and Supplementary Fig. 15h,i). Notably, cells from cluster 1 are nearly exclusively confined to the adult islet niche (Fig. 4m) and are in closer proximity to endocrine cells (Supplementary Fig. 15j). In contrast, cluster 4 was largely confined to mesenchymal niches (Supplementary Fig. 15g), similar to fetal G3 capillaries (Fig. 4c). Cluster 1 cells upregulate several genes characteristic of fetal IA capillaries, including *NOTCH4, INSR, ESM1*, *ACE*, and *SDC2* (Fig. 4n). Taken together, these findings demonstrate that the proto-islet niche is populated by diverse capillary subtypes during early development, with IA capillaries becoming progressively dominant over time. Cross-stage comparison with the adult pancreas suggests that IA capillaries represent a developmental precursor of cluster 1, the predominant islet-associated endothelial population in the mature organ (Fig. 4o).

Cross-tissue scRNA-seq studies have demonstrated remarkable transcriptional heterogeneity in organ-specific capillary ECs during development^50,51^, thought to underpin the specialized functional maturation of diverse microvascular beds in adult life. As pancreatic capillaries are absent or scarce in these datasets, we integrated developing pancreatic cells with a recent study performing Xenium spatial transcriptomics of an intact human embryo at 6 PCW^51^, and applied TERRA to facilitate cross-tissue analysis. TERRA successfully integrated and discriminated shared from tissue-restricted niches across datasets (Supplementary Fig. 16a-d). We next compared developing ECs across tissues, focusing on fetal pancreas capillary cells relative to other nascent microvascular beds (Supplementary Fig. 16e-g). Compared with other embryonic ECs, fetal pancreatic capillaries showed elevated activity of several SGPs, including the fetal pancreas SGP25 (Supplementary Fig. 16h-j). Among the genes comprising SGP25, the basement membrane components *COL4A1* and *HSPG2*, which are key constituents of the islet basement membrane^52^, together with capillary specialization markers *PLVAP*, *CLEC14A*, and *ACE*^46^ were upregulated in fetal pancreatic capillaries. SGP25 is expressed in fetal pancreatic capillaries, across all stages of pancreas development profiled (Supplementary Fig. 16k). These results suggest that fetal pancreatic capillaries undergo organ-specific transcriptional specialization early in pancreas development. Together, our analyses demonstrate that TERRA enables multi-scale spatial analysis of human pancreas development, despite never having been trained on pancreas tissue. In a zero-shot manner TERRA links cell identities, multicellular niches, and SGPs, uncovering both developmental trajectories and the emergence of islet-associated endothelial precursors.

### *In silico* perturbation with TERRA predicts a checkpoint-blockade-associated injury program in the human kidney

We then applied TERRA to test whether a broadly pretrained spatial foundation model could recover the highly ordered architecture of the human kidney in spatial transcriptomics data unseen by the model. The kidney is organized into anatomically and functionally specialized compartments, including the glomerulus, which facilitates plasma ultrafiltration, and the tubular nephron, which, together with its surrounding vasculature and interstitium, supports a diversity of physiological functions including acid–base balance, fluid homeostasis and control over endocrine signaling axes^53^. Recent spatial atlases of mouse and human kidneys have shown that these compartments are further structured into recurrent cellular niches associated with development, homeostasis and injury^54–62^. In particular, prior human kidney studies have resolved major kidney cell classes together with injury-associated epithelial and stromal states and localized these within spatial neighborhoods^54,60^, while other work has identified glomerular, immune, tubular and fibrotic microenvironments across healthy and diseased kidneys^57,62^.

We applied TERRA to build on this characterization of kidney niches, generating a unified, zero-shot analysis of human kidney cell types, niches and SGPs (Fig. 5a). We performed zero-shot analysis on nine unseen Xenium sections of non-tumor kidney tissue, profiled with the Xenium 5K panel, from five patients who underwent nephrectomy for localized renal cell carcinoma (RCC) (Supplementary Fig. 17a). After tokenizing and embedding these sections, we transferred cell-type labels from a public human kidney atlas^57^ using scANVI^63^, resolving 31 cell types; the nine untreated sections analyzed here comprised 1,080,937 cells, grouped into seven major compartments consistent with the kidney’s established microanatomical layout: proximal tubule, loop of Henle, collecting duct, glomerulus, immune, stromal and vascular across different estimated glomerular filtration rates (eGFR) (Supplementary Fig. 17b, c). Within these compartments, TERRA’s cell embeddings also resolved rare and specialized glomerular cell populations, identifiable by their canonical marker expression, including *VEGFR2*^+^ glomerular capillaries^64,65^, *NPNT*^+^ podocytes^66^, *CLDN1*^+^ parietal epithelium^67,68^ and *PDGFRB*^+^ mesangial cells^69,70^, each correctly localized to its expected anatomical position (Fig. 5b). Large-scale kidney atlases have previously characterized these same rare populations, but through iterative subclustering and manual annotation of each new dataset. Here, TERRA’s zero-shot embeddings already resolve them directly, in spatial context, requiring only reference-based label transfer rather than manual subclustering and re-annotation.

**Fig. 5.**
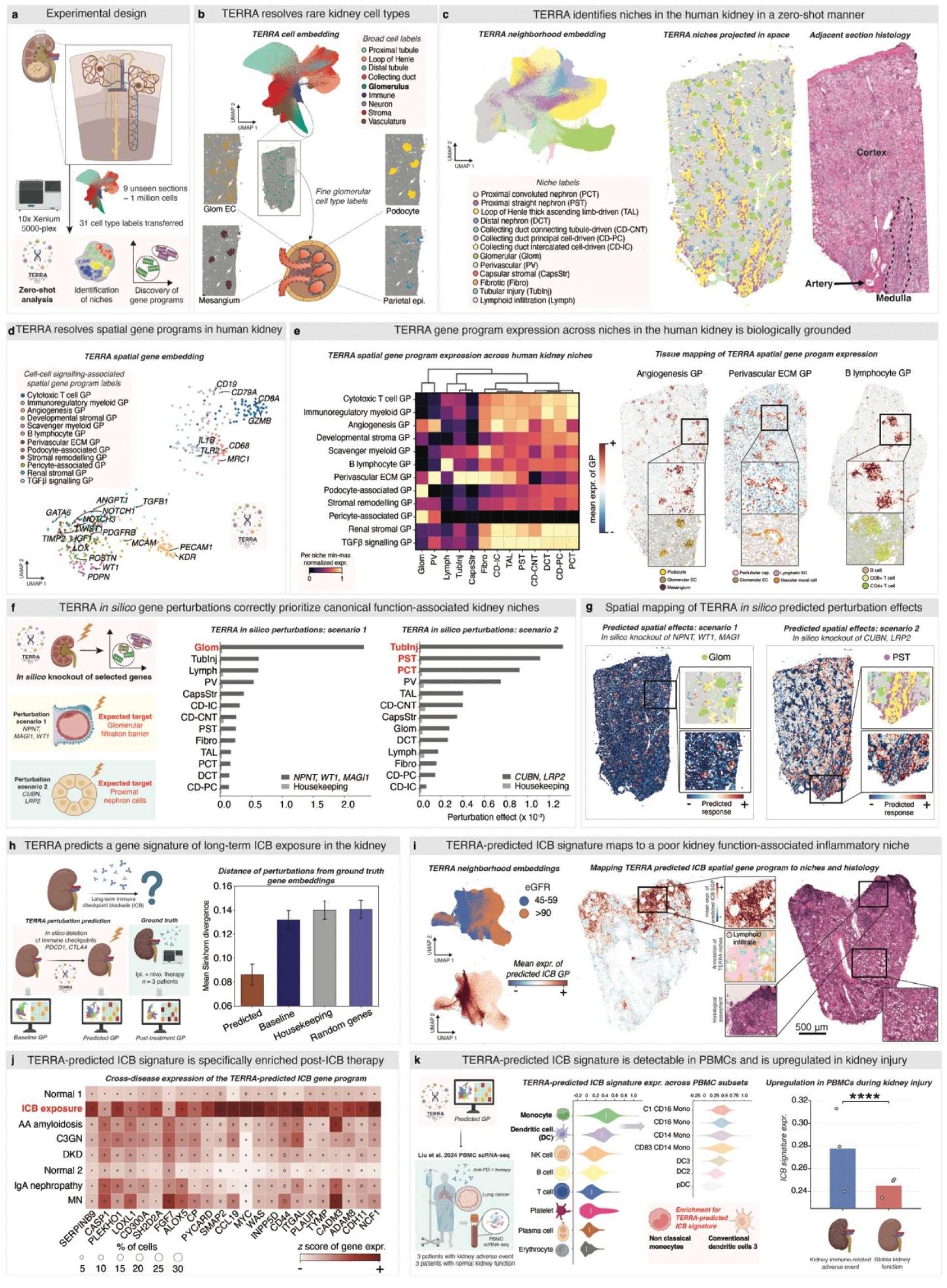
TERRA profiles the human kidney and predicts a checkpoint-blockade injury program detectable in blood. **a**, Experimental design: nine sections of non-tumor human kidney profiled with Xenium 5K and passed to pretrained TERRA; cell types (n = 31) transferred from a public reference atlas^57^. **b**, Top, UMAP of TERRA cell embeddings colored by broad compartment. Bottom, glomerular cluster magnified to resolve fine subtypes, mapped onto a representative section. **c,** Left, UMAP of TERRA neighborhood embeddings colored by niche (13 niches). Middle, niches projected in spatial coordinates. Right, H&E of the adjacent section (cortex, medulla, interlobular artery). **d**, UMAP of TERRA spatial gene embeddings colored by inter-cellular spatial gene program (SGP; 12 programs), with representative marker genes annotated. **e**, Left, heatmap of SGP expression (min–max normalized) across niches, hierarchically clustered. Right, spatial z-score of three representative SGPs, with contributing cell types indicated. **f**, Left, schematic of the *in silico* perturbation strategy. Middle/right, perturbation effect (W2 distance, ×10⁻³) across niches for knockout of podocyte-enriched genes at cell and neighborhood embeddings (scenario 1) versus megalin–cubilin complex genes (scenario 2), each versus housekeeping-genes knockout; strongest-responding niche highlighted in red. **g**, Spatial projection of per-cell W2 perturbation score for scenario 1 (left; NPNT, WT1, MAGI1) and scenario 2 (right; CUBN, LRP2), with strongest-responding niche overlaid (glomerular; proximal straight tubule). **h**, Left, schematic: *CTLA4*/*PDCD1* knocked out *in silico* at cell and neighborhood embeddings in treatment-naive sections and compared with real post-treatment sections (ipilimumab + nivolumab, n = 3 patients). Right, mean Sinkhorn divergence (± s.d.) from the real post-treatment gene program for TERRA’s predicted perturbation versus random-gene, baseline and housekeeping-gene knockouts. **i**, Left, UMAP of neighborhood embeddings colored by eGFR (top) and mean expression of the predicted ICB gene program (bottom). Middle, predicted ICB program (z-score) on a post-treatment section, with niche annotation highlighting a lymphoid infiltrate niche. Right, matching H&E. **j**, Cross-disease expression of the TERRA-predicted 23-gene ICB program: ICB-exposed versus untreated kidneys and four other kidney diseases. Dot size, % cells expressing; color, z-scored expression. **k**, Left, schematic: predicted ICB gene program scored in public PBMC scRNA-seq from six pembrolizumab-treated lung cancer patients (three with kidney toxicity, three without). Middle, expression across PBMC subsets, Myeloids subdivided into C1 CD16+ monocytes, CD16+ monocytes, CD14+ monocytes, CD83+ CD14+ monocytes, DC3, cDC2, pDC. Right, predicted ICB program expression with versus without kidney toxicity (Wilcoxon rank-sum test on all cells; **** indicates p < 0.0001).

Unsupervised clustering of TERRA neighborhood embeddings identified 13 kidney niches, annotated on the basis of cell-type composition and anatomical location (Fig. 5c and Supplementary Fig. 17d). These comprised capsular stromal, collecting tubule-driven, intercalated cell-driven, principal cell-driven, distal tubule-driven, fibrotic, glomerular, thick ascending limb (TAL)-driven, lymphocytic, perivascular, proximal convoluted tubule (PCT) nephron, proximal straight tubule (PST) nephron and tubular injury niches (Supplementary Fig. 17e). This niche organization aligns well with previous human kidney spatial studies^54^. In particular, the glomerular, lymphocytic, perivascular and fibrotic niches are concordant with previously described microenvironments^57,62^, whereas the compartmentalization of niches along the tubular nephron reflects the segmental specialization that underpins renal physiology. The tubular injury and fibrotic niches, enriched for *VCAM1*^+^ injured proximal tubules and myofibroblasts^60,71,72^, are also consistent with previously described maladaptive epithelial and fibro-inflammatory states associated with failed repair^54,56,59^.

We then extended the analysis to the gene level by examining contextualized SGPs learned by TERRA. Of the 5,001 genes in the Xenium panel, 4,947 were embedded in latent space, and clustering followed by manual annotation identified both intra-cellular SGPs, such as *Proliferation and Transcription* (Supplementary Fig. 18a-c), and inter-cellular SGPs associated with multicellular or signaling-related tissue context (Fig. 5d). Grouping inter-cellular SGPs by the niche in which they were most enriched revealed patterns that recapitulated known kidney physiology or pathology, each with expected spatial localization (Fig. 5e). For example, *angiogenesis* SGPs were prioritized in the glomerular niche, consistent with the secretion of vascular growth factors such as *VEGFA* and *ANGPT1* by podocytes, essential for the integrity of the glomerular capillary tuft^73–76^. Conversely, a *perivascular extracellular matrix (ECM)* SGP was localized to arterioles lined by mural cells and endothelial cells^62^, whereas the B lymphocyte SGP mapped to a lymphoid niche enriched for B cells and T cells, corresponding to tertiary lymphoid structure identity^62,77^. Together, these analyses show that TERRA can go beyond recovering established kidney niches in unseen tissue sections to resolve the niche-associated SGPs that underlie them.

We next leveraged TERRA’s ability to model *in silico* gene perturbations within cells and niches. As a proof of principle, we simulated knockout of genes with well-established, cell type-specific functions in kidney physiology (Fig. 5f). We first examined genes required for podocyte integrity and glomerular function, including the podocyte-enriched genes *WT1*^78^, *NPNT*^79^ and *MAGI1*^80^, by simulating their knockout across all cells and niches in healthy kidney spatial data. As expected, these perturbations produced their strongest effect in the glomerular niche (W2 distance: 2 × 10⁻³), exceeding both the effect of housekeeping-genes knockout (*PCNA* and *HPRT1*; W2 distance: 7 × 10⁻⁶) and the effect observed in every other niche. At the cell level, *in silico* knockout of *WT1*, *NPNT* and *MAGI1* was predicted to have the strongest effect in podocytes, mesangial cells, glomerular endothelial cells and parietal epithelium (Supplementary Fig. 19a), confirming that TERRA’s *in silico* perturbations capture known multicellular consequences of podocyte gene perturbation^78^. We then tested a second canonical pathway in kidney physiology by simulating knockout at both cell and niche level for *LRP2* and *CUBN*, which encode receptors of the proximal tubule-enriched megalin-cubilin complex required for reabsorption of proteins, vitamins and hormones from the urinary filtrate^81^. Consistent with known biology, perturbation of these genes had the greatest effect in proximal tubule-associated niches, with the strongest signal in the tubular injury niche (W2 distance: 1 × 10⁻³), the proximal convoluted tubule (W2 distance: 9 × 10⁻⁴) and proximal straight tubule (W2 distance: 1 × 10⁻³) niches (Fig. 5f and Supplementary Fig. 19b). Mapping these perturbation effects back onto tissue sections showed the expected spatial localization of glomerular filtration-barrier perturbations to glomerular regions and megalin–cubilin perturbations to proximal tubular regions (Fig. 5g). Together, these analyses indicate that TERRA’s *in silico* perturbation framework captures biologically meaningful, spatially localized gene dependencies in native tissue context.

We next asked whether TERRA’s *in silico* perturbations could predict how niches and gene programs shift in a clinically relevant pathological setting. Immune checkpoint blockade (ICB) and VEGF-targeted tyrosine kinase inhibitors (TKIs) are a cornerstone for the management of kidney cancer^82,83^ and other malignancies, but are associated with adverse renal events, manifesting in acute kidney injury and necessitating cessation of therapy^84,85^. To test whether TERRA could predict treatment-associated molecular shifts, we performed *in silico* knockout at both cell and niche level in untreated kidney sections of genes encoding the targets of the ICB agents ipilimumab and nivolumab, namely *CTLA4* and *PDCD1*. As a ground-truth reference against which to compare our perturbations, we generated Xenium 5K data from non-tumor kidney tissue from nephrectomy specimens of patients with RCC, collected either before systemic therapy or after treatment with ICB: either nivolumab alone, or in combination with ipilimumab (Fig. 5h). Simulated knockout of *CTLA4* and *PDCD1* shifted the nine untreated sections toward a post-treatment state: the mean Sinkhorn divergence of neighborhood gene embeddings from the real post-treatment reference was lower for these knockouts (∼0.08) than for unperturbed controls (∼0.13), random-gene perturbations (∼0.14) or housekeeping-gene perturbations (∼0.14). Random-gene and housekeeping-gene perturbations were each associated with a greater mean Sinkhorn divergence than the corresponding knockout. Thus, TERRA’s perturbations moved tissue representations in a direction more consistent with the observed therapy-exposed kidney state than either unperturbed or biologically unrelated controls did.

ICB-related kidney injury most commonly manifests as acute interstitial nephritis or other forms of acute tubulointerstitial injury^84,86^. Although these clinicopathological associations are well recognized, it remains unclear to what extent prolonged exposure to these therapies reshapes kidney spatial organization and molecular architecture. We therefore sought to leverage TERRA to further characterize how niches and gene programs are altered by long-term ICB exposure. Ranking the TERRA-predicted shift in neighborhood gene embeddings after *CTLA4* and *PDCD1* knockout, we identified 23 genes among the top predicted shifts that were also significantly differentially expressed between real untreated and ICB-treated kidney sections (Fig. 5i). We calculated an ICB-associated SGP from these 23 genes and projected this onto a joint neighborhood embedding containing treatment-naive and treatment-exposed kidney sections. Within this embedding space, the ICB-associated GP mapped to regions from patients who have lower eGFR, a surrogate marker of kidney function. Projecting the ICB-associated GP predicted by TERRA onto the post-treatment sections further showed that this gene program was enriched in regions containing lymphoid infiltrate, a finding supported both by niche annotations and by hematoxylin and eosin staining of aligned sections.

To determine whether TERRA was detecting a GP specific to long-term exposure of the kidney to ICB, or whether it simply reflected a more generic signature of inflammation in the kidney, we performed a cross-disease analysis using publicly available spatial data from multiple kidney diseases spanning inflammatory and metabolic etiologies^62^ (Fig. 5j). Individual genes within the signature were not, on their own, specific to ICB. The pro-inflammatory chemokine *CCL19*, for example, was also elevated in other inflammatory kidney diseases such as C3 glomerulonephritis and membranous nephropathy, consistent with its general role in organizing lymphoid infiltrates across chronic kidney disease^87^. However, the composite 23-gene GP predicted by TERRA, when considered as a coordinated program rather than as individual markers, was largely specific to kidneys with long-term ICB exposure relative to these other kidney diseases, indicating that this spatially coordinated combination of genes distinguishes ICB-exposed tissue from different etiologies of kidney inflammation. Finally, we asked whether the ICB-associated GP predicted by TERRA was generalizable beyond kidney tissue itself, and whether it could be detected in a more readily accessible clinical sample. We extracted publicly available scRNA-seq data of 35,088 PBMCs from six patients with lung cancer treated with pembrolizumab, an alternative ICB that targets PD-1^88^, which is known to increase disease free survival in kidney cancer post-surgery^89^. Of these six patients, three were documented to have ICB-associated kidney injury. We used CellTypist^90^ to annotate these PBMCs. We then computed the expression of the ICB-associated GP predicted by TERRA. This GP was enriched in two myeloid populations, non-classical monocytes and DC3 (Fig. 5k). DC3 is a recently defined dendritic cell subset. It arises from a distinct, monocyte-related developmental lineage^91^. DC3 has also been linked to priming of tissue-resident memory T cells^92^. Across the aggregated dataset, the ICB-associated GP predicted by TERRA was expressed at significantly greater levels in PBMCs from patients with documented ICB-associated kidney injury than in patients with stable kidney function (Fig. 5k) (∼11% higher in ICB-associated kidney injury).

Together, these results establish TERRA as a unified framework for profiling the kidney at cell, niche and gene resolution. The *in silico* perturbations proved grounded enough in real biology to predict, from treatment-naive kidney, a GP unique to long-term ICB exposure, a program that maps to a histologically evident lymphocytic niche in post-treatment tissue, is detectable in blood and is associated with kidney dysfunction.

### TERRA defines cross-tissue spatial archetypes of macrophages in human health and disease

That certain niches recur across organs is well established, from tertiary lymphoid structures to fibrotic and perivascular niches. We therefore extended TERRA’s ability to resolve niches within a single tissue to systematically map which cellular ecosystems, and the gene programs that define them, are conserved across organs and which are organ-specific, addressing a central challenge in biology: the principles that govern conserved versus organ-specific tissue organization^5,93^. We define an archetype as a group of niches that recur across tissues and share both a conserved local cellular composition and the cell gene programs (GPs) their cells express. In adopting the term archetype, we build on its use to describe recurring, prototypical patterns of immune organization across cancers^94^ and, more recently, analogous spatial-functional patterns among stromal cells^95^. We applied this framework to macrophages, which are pervasive across human tissues, either tissue-resident or derived from infiltrating monocytes, and, beyond conserved functions such as clearing debris, defending against infection, and linking innate and adaptive immunity, are highly heterogeneous in both ontogeny and local microenvironment^96–100^, giving rise to distinct subsets in the liver^101^, skin^102^ and kidney^103,104^, among several organs. Disease adds further heterogeneity, from immunosuppressive macrophages in cancer^105^, to lipid-laden macrophages in atherosclerosis^106^ and other pathologies^107–109^. To understand archetypes of macrophages, we leveraged the subset of adult human tissues from HST-5K-Corpus (∼28M cells, 276 Xenium sections, nine organs: brain, lung, liver, kidney, breast, skin, ovary, cervix and prostate; Fig. 6a and Supplementary Fig. 1e,f). After integrating TERRA’s neighborhood embeddings across HST-5K-Corpus (Methods), we identified macrophages using pan-macrophage (*CD68, CD163*) and population-specific markers, resolving 1,522,556 macrophages within TERRA’s embedding space (Fig. 6b,c), including specialized subtypes such as *MRC1+ MARCO+* alveolar macrophages^110^ (Supplementary Fig. 20a) and *TMEM119+ P2RY12+* microglia^111^. Unsupervised clustering revealed 298 macrophage-enriched niches, classified as tissue-private or cross-tissue (Supplementary Fig. 20b), providing a data-driven framework to systematically characterize macrophage archetypes across human tissues.

**Fig. 6.**
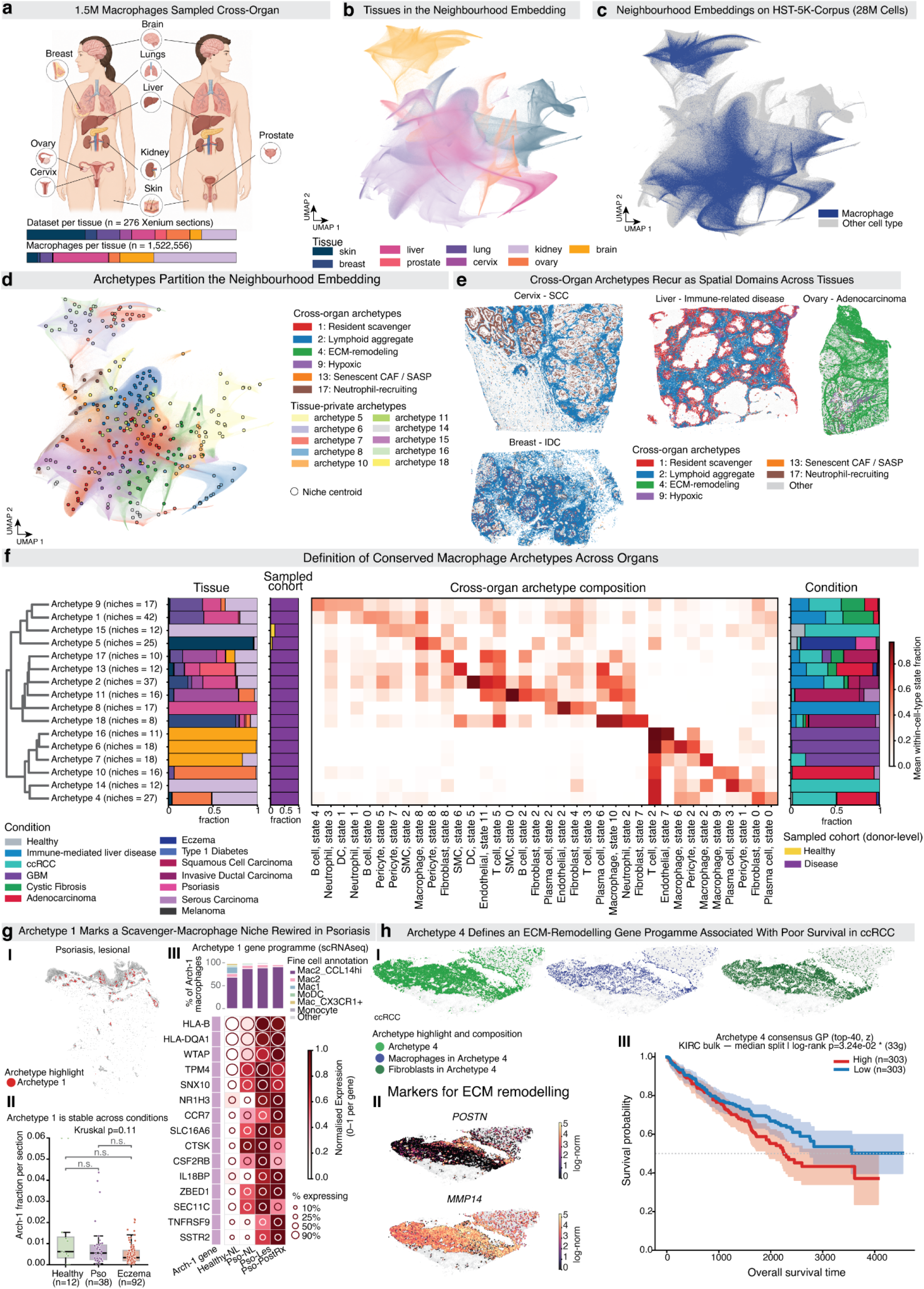
Cross-tissue macrophage niche archetypes derived from the TERRA neighborhood embedding. **a**, Overview of the cross-organ macrophage atlas. Xenium data were assembled across nine adult organs (skin, breast, lung, cervix, liver, prostate, ovary, brain, kidney), comprising 276 Xenium sections and 1,522,556 macrophages; pancreas was excluded from all analyses because the available sections were mixed with developing tissue. Stacked bars show the per-tissue composition of the dataset by section count (top) and by macrophage count (bottom); colors denote tissue of origin. **b**, UMAP of the TERRA neighborhood embedding of HST-5K-Corpus (28M cells; non-embryonic, pancreas excluded), colored by tissue of origin (skin, breast, lung, cervix, liver, prostate, ovary, brain, kidney). **c**, UMAP of TERRA neighborhood embeddings of HST-5K-Corpus, with macrophages colored dark blue and all other cell types gray. **d**, Archetypes partition the neighborhood embedding. Niche centroids (open circles) are overlaid on the neighborhood embedding UMAP, colored by the archetype assignment of the underlying cells. Six recurrent cross-organ archetypes are named: Resident scavenger (1), Lymphoid aggregate (2), ECM-remodeling (4), Hypoxic (9), Senescent CAF / SASP (13) and Neutrophil-recruiting (17); the remaining ten archetypes (5, 6, 7, 8, 10, 11, 14, 15, 16, 18) are tissue-private (legend). Archetype identifiers are retained from the original solution and are therefore non-consecutive. **e**, Cross-organ archetypes recur as spatial domains across tissues and disease contexts. Representative Xenium sections (cervix, squamous cell carcinoma; liver, immune-mediated liver diseases; ovary, adenocarcinoma; breast, invasive ductal carcinoma) with cells colored by their assigned cross-organ archetype: Resident scavenger (1, red), Lymphoid aggregate (2, blue), ECM-remodeling (4, green), Hypoxic (9, purple), Senescent CAF / SASP (13, orange) and Neutrophil-recruiting (17, brown). Cells in tissue-private niches or without an archetype assignment are shown in gray. **f**, Definition of conserved macrophage archetypes across organs. Macrophage-enriched niches (n = 298) were grouped by Ward linkage on their cell-type-state composition; two archetypes composed almost entirely of developmental-pancreas niches were removed post hoc, yielding 16 cross-tissue archetypes (dendrogram, left; niche count per archetype in parentheses). Stacked bars show, per archetype, the tissue composition (Tissue; colors as in **a**) and the disease-condition composition (Condition; Healthy, Immune-mediated Liver Disease, ccRCC, GBM, Cystic Fibrosis, Adenocarcinoma, Eczema, Type 1 Diabetes, Squamous Cell Carcinoma, Invasive Ductal Carcinoma, Psoriasis, Serous Carcinoma, Melanoma). The Sampled cohort bar shows the donor-level healthy versus disease split (Healthy, yellow; Disease, purple) and reflects the sampling cohort rather than per-section lesional status. The heatmap shows the mean within-cell-type state fraction across the top distinguishing cell-type-state features per archetype (white to dark red, color bar); this value is normalized within each cell type and is therefore independent of overall cell-type abundance. **g**, Archetype 1 marks a scavenger-macrophage niche transcriptionally rewired in psoriasis. **(I)** Representative psoriatic lesional section with Archetype 1 cells colored red on all other cells in gray, showing dermal localization of the niche. **(II)** Archetype 1 abundance is stable across conditions: per-section Archetype 1 fraction in healthy (n = 12), psoriatic (Pso, n = 38) and eczematous (Eczema, n = 92) skin (box, interquartile range and median; points, individual sections; Kruskal-Wallis p = 0.11; all pairwise comparisons not significant). **(III)** Archetype 1 gene program in matched single-cell RNA-seq. Top, composition of Archetype 1 macrophages by fine cell annotation across conditions (Mac2_CCL14hi, Mac2, Mac1, MoDC, Mac_CX3CR1+, Monocyte, Other). Bottom, matrix dot-plot of program genes across Healthy-NL, Pso-NL, Pso-Les and Pso-PostRx; square color, mean expression scaled 0 to 1 per gene (white to dark red); circle size, percentage of cells expressing (key); the left strip marks Archetype 1 identity genes. **h**, Archetype 4 defines an ECM-remodeling gene program associated with poor survival in ccRCC. **(I)** Representative ccRCC section showing Archetype 4 (green), the macrophages within Archetype 4 (blue) and the fibroblasts within Archetype 4 (dark green). **(II)** Expression of the ECM-remodeling markers *POSTN* and *MMP14* on the section (log-normalized expression, dark to yellow). **(III)** Kaplan-Meier overall-survival analysis in the TCGA KIRC bulk cohort, stratified by a median split on the z-scored Archetype 4 top-40-gene consensus GP (33 genes mapped to the cohort); High (n = 303, red) versus Low (n = 303, blue); shaded bands, 95% confidence intervals; log-rank p = 3.24 x 10⁻².

Grouping these niches by shared cell-type-state composition (Methods) partitioned the embedding into 16 archetypes (Fig. 6d) that explained 76.2% of inter-niche compositional variance (PERMANOVA R² = 0.762, p < 1 × 10⁻⁴, 9,999 permutations), with 5–37 cell-type-states significantly enriched per archetype (Mann–Whitney U, BH-corrected q < 0.05), confirming that archetypes capture structured, biologically meaningful variation rather than clustering artifacts. Six of the 16 archetypes spanned multiple tissues and conditions (cross-tissue archetypes; Fig. 6e), while the remainder were tissue-confined. Pseudobulk profiling and examination of tissue, condition and cell-type composition identified the defining GP of each (Fig. 6f and Supplementary Fig. 20c): archetype 1 (*Resident scavenger*; *CD206, NR4A1, INMT*) occupied vasculature-associated niches of endothelial cells and fibroblasts, consistent with homeostatic tissue-resident scavenging^103,112,113^; archetype 2 (*Lymphoid aggregate*; *CD3E, CD8A, CD79A, MS4A1, CCL19*) suggested tertiary lymphoid structure identity^114,115^; archetype 4 (*ECM-remodeling*; *COL4A1/2, COL18A1, MMP14, THBS1*) occupied fibroblast- and pericyte-rich stroma; archetype 9 (*Hypoxic*; *HIF1A, CXCR4, POU2AF1, CYBA*) occupied hypoxic B-cell niches; archetype 13 (*Senescent CAF / SASP; CCN1, CCN2, ADAMTS1*) occupied senescent fibroblast and luminal-epithelial-rich niches; and archetype 17 (*Neutrophil-recruiting*; *CXCR1, CXCR2, C5AR1, FCGR2A, BCL2A1*) occupied epithelial niches with neutrophil infiltration. We note that the genes defining each archetype capture both the intrinsic state a macrophage adopts and the composition of the niche it occupies, since the two are coupled in situ. Rather than describing macrophage identity in isolation, each archetype resolves cell state and microenvironmental context as a single, jointly defined spatial axis, moving beyond the conventional treatment of the two as separable (see Discussion).

We next asked whether such structurally conserved archetypes are fixed, or can be remodeled by disease within a single tissue. We focused on archetype 1 (*Resident scavenger*), the most ubiquitous archetype, spanning healthy, psoriatic and eczematous contexts in skin sections^116^. Archetype 1 localized to the dermis (Fig. 6g-I), with no difference in per-section abundance between conditions (Kruskal–Wallis p = 0.11; Fig. 6g-II), indicating a structurally constitutive spatial domain. To ask whether this niche was nonetheless molecularly reprogrammed, we scored its GP in an independent scRNA-seq dataset spanning healthy, non-lesional, lesional and post-treatment psoriatic skin^116^ (Fig. 6g-III). A *CCL14*+ macrophage subset, whose GP is reported to be enriched in psoriatic lesions^117^, showed the highest expression of the archetype-1 gene program among myeloid cells, and its share of the archetype-1 macrophage compartment rose with disease activity (Jonckheere trend p = 0.016), remaining elevated after treatment. Within these cells, GP activity itself shifted with disease state. Genes governing antigen presentation (*HLA-B, HLA-DQA1*) and lipid handling (*NR1H3*) rose with disease and remained elevated post-treatment, whereas *IL18BP*, encoding the IL-18 antagonist^118–120^, and *CCR7*, which mediates immune egress^121,122^, were higher after treatment than during active disease. Together, these results show that an archetype’s structural presence can remain stable across disease states even as the cell population occupying it, and the genes it expresses, both change with disease activity.

We asked whether macrophages within a different cross-tissue archetype might instead occupy a pathological axis. Archetype 4 (*ECM-remodeling*) was enriched for transcripts governing matrix turnover (*MMP14, MRC2, LRP1*), deposition (*COL4A1/2, COL5A1/2*) and crosslinking (*LOXL1, LOXL2*). Archetype 4 macrophages co-localized most consistently with fibroblasts across the niche compendium (Supplementary Fig. 20c) and the archetype was most abundant in spatial data from kidney (ccRCC) and ovarian (adenocarcinoma) malignancy (Fig. 6f), settings where ECM-remodeling and fibrosis^123,124^ are drivers of immune exclusion^125–127^. Scoring an independent ovarian cancer atlas^128^ with the archetype 4 GP highlighted stromal and myeloid compartments (Supplementary Fig. 21a) and identified its constituent fibroblasts as matrix-remodeling CAFs (mCAFs; Cohen’s d > 1.0, BH-q < 0.05; Supplementary Fig. 21b), an ECM-producing state recently proposed as conserved across cancers^95^. This conservation was reproducible across ccRCC, LUAD, pediatric ovarian adenocarcinoma and melanoma within our datasets (mean Cohen’s d = 1.31; Supplementary Fig. 21c). Archetype 4 macrophages independently expressed this same program across the same tumor types (mean Cohen’s d = 1.51) albeit more weakly in melanoma (Supplementary Fig. 21d), and most closely resembled resident *C1QC+* and *LYVE1+* tumor-associated macrophage states (Supplementary Fig. 21e). We examined archetype 4 in two ccRCC sections^24^, where it mapped to macrophage- and fibroblast-rich fibrotic regions at the tumor–kidney interface, expressing *POSTN* and *MMP14* (Fig. 6h-I,II), and spanned perivascular, hypoxic and matrix-remodeling niches (Supplementary Fig. 22c,e). Because ECM-remodeling is carried out jointly by macrophages and fibroblasts, we asked whether the archetype 4 program is recoverable within each population separately. Ordering fibroblasts and macrophages independently along a diffusion pseudotime on their TERRA cell embeddings, we found archetype 4 genes expressed along both trajectories (Supplementary Fig. 22a,d). This GP coincided with the matrix-remodeling spatial domain defined directly from the niches (Supplementary Fig. 22b,c,e), localized to the fibrotic tumor–kidney interface. Extending beyond these two sections, archetype 4 recurred across an independent cohort of seven ccRCC tumor-boundary sections from different patients, where it was roughly two-fold enriched relative to tumor-core sections (median 34% vs 16% of cells; one-sided Mann–Whitney p = 0.019, OCT sections), and its fibroblasts and macrophages expressed the matrix-remodeling program in every section examined (Supplementary Fig. 23). Together, these results define archetype 4 as a conserved, cross-cancer matrix-remodeling niche in which macrophages and fibroblasts express a shared ECM program, spatially organized at the fibrotic tumor interface.

Finally, to examine the clinicopathological significance of archetype 4, we scored the GP in bulk RNA-seq from the Cancer Genome Atlas ccRCC cohort^129^. High expression of the archetype 4 GP was associated with significantly worse overall survival (top-40 z-scored signature, 33 genes mapped; Cox HR = 1.48, 95% CI 1.12–1.95, p = 0.005; median split, log-rank p = 3.2 × 10⁻²; Fig. 6h-III).

Together, we illustrate how TERRA can be leveraged to systematically group niches into recurrent, prototypic spatial archetypes in human tissues. By generating a catalog of macrophage-enriched archetypes, and by contextualizing two of these in inflammatory skin diseases and kidney cancer, we illustrate how a spatial foundation model can reveal how disease reprograms a niche’s molecular identity independently of its spatial architecture, and can be leveraged to identify GPs of prognostic significance, directly from spatial data, across organs and contexts, without any tissue-specific retraining.

## Discussion

TERRA is a self-supervised spatial foundation model, pretrained on more than 112 million human cells profiled by imaging-based spatial transcriptomics. TERRA is built around a context-aware tokenization that represents each cell together with its local tissue environment while preserving gene identity, and is trained with a joint-embedding predictive objective that infers masked molecular and spatial context directly from neighboring cells. Across the applications above, we demonstrate how TERRA leverages a vast, diverse *in situ* corpus to facilitate zero-shot identification of niches and cell types and their underpinning coordinated programs of genes in unseen sections, organs, and disease contexts without requiring task-specific fine-tuning. Extending this to biologically grounded and experimentally validated *in silico* perturbations, as well as cross-tissue cataloging of niches into spatial archetypes, we illustrate how a single, unified model achieves what single-cell foundation models trained on dissociated tissues cannot support by design, and that existing spatial foundation models only address in part. In particular, whereas recent perturbation-prediction models learn transcriptional responses from large perturbation screens of dissociated cells^130–134^, such data remain scarce for spatial assays; TERRA offers a complementary first step, using its large observational corpus to model *in silico* how a perturbation propagates through a cell and its surrounding niche.

Existing foundation models trained on dissociated scRNA-seq^8–10^ learn transferable gene-level structure but cannot observe the native tissue context in which gene networks, cell–cell communication and multicellular responses are organized. Conversely, recent spatial models^18,20,21^ capture aspects of tissue context but typically operate at a single scale, depend on dataset-specific training, or do not retain gene-level resolution suitable for interpretation and perturbation. Several design choices distinguish TERRA, and our ablations clarify why. First, predicting in latent space rather than reconstructing observed counts is well matched to spatial transcriptomics data, which are sparse, noisy and acquired with heterogeneous targeted panels. The predictive objective encourages the model to capture stable microenvironmental structure rather than memorize technology-specific noise. Second, TERRA’s combined tokenization encodes both a within-cell expression rank and the gene’s expression value, which outperformed encoding either the rank or value alone. The expression rank provides an implicit, scale-free normalization robust to library-size and sensitivity differences, making explicit normalization unnecessary and sometimes counterproductive. Meanwhile, the expression value preserves quantitative expression that ranking discards. Third, a lightweight batch metatoken is sufficient to integrate data across assays without any explicit batch correction loss or post-hoc alignment; removing it collapses cross-assay mixing, whereas retaining it yields shared embeddings while preserving biological structure. Together these properties allow a single model to span five imaging-based assays. By learning representations from such diverse data that are simultaneously gene-resolved and spatially contextualized, and by demonstrating strong zero-shot transfer and the ability to perform *in silico* gene perturbations, TERRA addresses a combination of requirements that prior models met only in part.

TERRA enabled recovery of both established and novel spatial biology in several zero-shot settings. Using a version of the model that had not been trained on pancreatic tissue, TERRA recovered expected epithelial trajectories during pancreatic development, and a large-lumen-ductal niche transcriptionally and spatially distinct from canonical pancreatic ductal cells. Although lineage relationships remain to be established, the subsequent acquisition of pancreatic ductal identity is consistent with the possibility that the large-lumen duct population reflects extrahepatic biliary-derived epithelium that integrates into the developing pancreatic ductal network^35,36^. Similarly, in unseen human kidney sections, TERRA facilitated zero-shot recapitulation of the highly ordered cellular architecture of the nephron and glomerulus identified by scRNA-seq studies, while capturing epithelial and immune niches which have previously been identified in kidney spatial atlases^54–62^. While prioritizing SGPs to niches with concordant biological functions in the kidney, TERRA also resolved an SGP corresponding to progressive angiogenic activity in the emerging proto-islet niche. We explored this further, and leveraged TERRA to identify a specialized islet-associated capillary state enriched within the direct precursors to pancreatic islets^46,47^.

These IA capillaries likely represent the progenitors that form specialized pancreatic islet ECs, and we identify tissue-specific candidate molecules that underpin their identity and specification, with implications for understanding how ECs shape pancreatic islet maturation^38,44^.

Extending the model to perform *in silico* perturbation of gene expression, TERRA also predicted the molecular consequences of long-term ICB therapy on the kidney, building on prior studies capturing the acute consequences of ICB nephrotoxicity, which have been clinically^84,86^ and molecularly^135^ interrogated. In nephrology, AI models have largely been used for patient or tissue-level predictions, such as anticipating risk of acute kidney injury^136^, or identifying clinically relevant features during histopathological assessment^137^. More recently, generative frameworks have been implemented for modeling proteins to interrogate the structural biology underpinning renal physiology and disease^138^ or for zero-shot annotation, cross-species analyses and *in silico* perturbations of scRNA-seq data^139^. TERRA extends these efforts to spatial transcriptomics data, using *in silico* knockout of ICB target-encoding genes in untreated kidney tissue to nominate a 23-gene signature, validated in kidney cortex exposed to long-term ICB usage where it mapped to lymphocytic infiltrates, detectable by scRNA-seq of PBMCs and upregulated in patients with clinically established ICB-associated kidney injury. Although it is not clear if ICB therapy reprograms circulating immune cells which then infiltrate the kidney, or if reprogrammed cells from the kidney are released into the blood, our findings nevertheless illustrate how a broadly trained spatial foundation model can be leveraged to anticipate clinically consequential cellular and molecular changes induced by specific therapies in tissue sections unseen to the model.

TERRA’s 112M cell pretraining corpus is, to our knowledge, among the largest, and the largest composed exclusively of single-cell-resolution imaging-based data, used to pretrain a spatial foundation model to date. This vast corpus provided the substrate from which we drew a targeted, nine-organ, 28-million-cell subset to systematically group macrophage niches into recurrent, prototypical spatial archetypes^3,140,141^, six of which recurred across multiple organs and disease contexts. Prior work on macrophage heterogeneity has largely cataloged intrinsic cell states through pan-cancer TAM atlases^142–145^, treating spatial context separately, if at all. By defining archetypes that couple each macrophage’s state to its niche, we resolve the two as a single object rather than as separable layers, which to our knowledge has not been done for macrophages at this scale. One of these was a macrophage-fibroblast-enriched, ECM-remodeling archetype, recurring across kidney, lung, ovarian and skin cancers, with an SGP that was prognostic in an independent cohort of patients with kidney cancer, where it localized to fibrotic tissue at the tumor-kidney interface. The location and composition of this archetype were consistent with the RCC pseudocapsule, an anatomically and clinically established structure whose invasion status is itself prognostic^146,147^. Peritumoral, fibroblast-rich stroma is not unique to RCC, and is a recognized route to T cell exclusion and ICB resistance in other cancers^148^. Thus, TERRA provides a generalizable framework to systematically compare the context of cell types and niches across diverse human spatial datasets, supporting biological discovery at scale.

The present study is bound by certain limitations. TERRA is pretrained on imaging-based assays with targeted gene panels, so gene coverage is constrained, although we show that the model can be adapted efficiently to sequencing-based platforms such as 10x Visium with lightweight fine-tuning. We anticipate it will extend to future or emerging assays as they enter the corpus. Neighborhoods are defined by a fixed k-nearest-neighbor graph; niche and gene program definitions still require expert interpretation; and evaluation relies on imperfect reference annotations that may underestimate the true quality of the learned representations. Moreover, the zero-shot analyses of pancreas and kidney were performed in small donor cohorts, limiting our ability to account for biologically meaningful variability such as biological sex or inter-patient heterogeneity. We anticipate that these limitations will be progressively overcome as *in situ* corpora continue to grow and diversify across tissues, conditions and platforms, and as natural extensions are realized, including joint pretraining with paired histology and protein measurements, modeling of three-dimensional and temporally resolved tissue, and tighter coupling between representation learning and experimental design.

More broadly, TERRA reframes spatial transcriptomics analysis from a collection of bespoke, dataset-specific pipelines toward a single reusable model that contextualizes each new experiment within the accumulated knowledge of a large in situ corpus. Across other domains of biology, foundation models have become reusable substrates whose learned representations are adopted far beyond their original task, from vision transformers for images^149^, to protein language models such as ESM^150^, to sequence-to-function models of the genome such as AlphaGenome^151^ and Borzoi^152^. TERRA offers an analogous foundation for spatial transcriptomics: a universal embedding of spatially resolved tissue that is a resource not only for biologists interpreting new sections, but also a substrate for building further AI models, including systems that integrate spatial transcriptomics with complementary modalities such as histological imaging, sequencing-based assays and protein measurements. As spatial technologies scale from targeted panels toward genome-wide interrogation across ever-larger tissue areas, we envision TERRA as a step toward foundation models that not only describe human tissue but predict how it responds to perturbation, providing a generalizable framework for discovery across tissue biology, therapeutic development and clinical application.

## Acknowledgments

We acknowledge core funding from the Wellcome Trust (WT220540/Z/20/A) and a program grant from the Medical Research Council (APP73647). M.H. is funded by Wellcome (WT107931/Z/15/Z) and the CIFAR MacMillan Multiscale Human program. M.L. and M.H. acknowledge support and funding from Open Targets. T.M. is supported by Krishnan-Ang Philanthropy. S.B. is supported by the Helmholtz Association under the joint research school “Munich School for Data Science—MUDS”. D.J.J. acknowledges support from a Wellcome Trust Accelerator Award (314710/Z/24/Z), a Kidney Research UK Project Grant (SG_MNRP_006_20250929) and from the NIHR Biomedical Research Centre at Great Ormond Street Hospital for Children NHS Foundation Trust and University College London. S.W. is funded by the Royal Society (CDF/R1/241008), and supported by the NIHR and Newcastle Biomedical Research Centre. L.S. is supported by a Wellcome Clinical Research Training Fellowship and Trinity Internal Graduate Studentship. The Francis Crick Institute receives its core funding from Cancer Research UK (CC2256), the UK Medical Research Council (CC2256), and the Wellcome Trust (CC2256). Additionally, F.J.R is supported by a personal fellowship from the UK Medical Research Council (MR/X019500/1) and J.J.S is supported by an MRC Clinical Research Training Fellowship (UKRI3629). The authors thank Nisha Bhardwaj and Richard Stone of Experimental Histopathology at the Francis Crick Institute for their technical support and guidance in this work. M.B. is supported by the Helmholtz Society, Helmholtz Portfolio Theme’Metabolic Dysfunction and Common Disease and German Center for Diabetes Research (DZD). We thank Aidan Maartens (Wellcome Sanger Institute) for providing feedback on the manuscript. S.B. is grateful to Paula, Pebble and Pixel Villa Fulton for their inspirational support. This research was performed with the support of the Network for Pancreatic Organ donors with Diabetes (nPOD; RRID:SCR_014641), a collaborative type 1 diabetes research project supported by Breakthrough T1D and The Leona M. & Harry B. Helmsley Charitable Trust (Grant#3-SRA-2023-1417-S-B). The content and views expressed are the responsibility of the authors and do not necessarily reflect the official view of nPOD. Organ Procurement Organizations (OPO) partnering with nPOD to provide research resources are listed at https://npod.org/for-partners/npod-partners.

## Author Contributions

S.B., A.V., and M.L. conceived the project with later contributions from M.V.S. who helped mature the method. S.B., M.V.S. and A.V. designed the method with contributions from M.L.; S.B. and M.V.S. implemented the method with contributions from A.M.; S.B., A.M., and B.C. curated and harmonized the datasets for pretraining. S.B., A.M., M.V.S., and Z.H. ran the benchmarks. S.O. and M.B. performed the pancreas analyses with feedback from D.J.J.; D.J.J. and A.M.F. performed the kidney analyses with feedback from A.A.; C.L. performed the archetype cross-tissue analyses with feedback from D.J.J., M.L., O.A.B. and M.H.; S.B., M.V.S., and A.V. supported all analyses. L.S., K.Ro., K.Ra., A.B., M.R.S., J.M., C.R.-S., B.C., S.M.S., V.L., J.M.B., C.K., J.O.J., G.D.S., B.R., C.T., M.Pa., T.L., H.S., A.R.F., A.L.T., D.A., S.W., J.J.S, F.J.R., M.Pr., Z.H., and D.J.J. contributed to sample collection, experimental work, and data generation and processing. O.A.B., M.N., R.V.T. and M.R.C. provided data and resources. S.B., D.J.J., S.O., A.M.F., C.L., O.A.B., T.M., M.H., M.V.S., M.B. and M.L. wrote the manuscript. F.J.T., H.L. and L.P. provided support for S.B., M.B., and M.V.S.; T.M., M.H., M.B. and M.L. provided supervision. M.L. designed the overall experiments and scientific direction. All authors read and approved the final manuscript.

## Methods

### Pretraining dataset: HST-Corpus-112M assembly and harmonization

We assembled HST-Corpus-112M, a corpus of more than 112 million single-cell-resolved spatial transcriptomic profiles (112,578,039 cells) from 636 tissue sections (batches) generated with five imaging-based assays (Xenium, MERFISH, CosMx, STARmap and ISS CARTANA), spanning 20 human tissues and including healthy tissue and 26 disease conditions (Fig. 1c and Supplementary Fig. 1). Of these, 75,552,216 cells (67%) were generated in-house by the authors and 37,025,823 cells (33%) were obtained from public resources, primarily 10x Genomics; per-dataset provenance is provided in the Data Availability section. Datasets were harmonized and preprocessed with a common pipeline available at github.com/Lotfollahi-lab/terra-reproducibility, producing standardized AnnData objects with spatial coordinates in micrometers and a shared gene namespace. Basic quality control removed cells with fewer than 10 expressed genes, and only genes with valid Ensembl identifiers (release 111) were retained. No statistical method was used to predetermine sample size, and no data were excluded unless explicitly stated.

To separate model development from zero-shot evaluation, we trained TERRA-112M on the full corpus and TERRA-96M on a 96-million-cell subset (421 batches) after holding out 215 batches for benchmarking and analysis of unseen datasets (Fig. 1c). We additionally defined HST-5K-Corpus, a Xenium-only subset of sections profiled with the Xenium Prime 5,000-gene panel, comprising 40,712,678 cells across 21 datasets (Supplementary Fig. 1e,f), used for the cross-tissue analyses.

### Analysis datasets

All datasets used to evaluate and apply TERRA were held out from the pretraining of the model used to analyze them. Unless otherwise stated they were analyzed zero-shot, without fine-tuning; fine-tuning was used only for niche label transfer (Fig. 2b) and for transfer learning to 10x Visium (Fig. 2c). All were profiled with imaging-based spatial transcriptomics; where specified, matched single-cell or single-nucleus RNA-seq data were used for annotation and validation, and all human samples were obtained with appropriate ethics approvals and informed consent. All benchmarking and analysis datasets are described below; per-dataset provenance is listed in the Data Availability section.

### Benchmarking datasets

For zero-shot evaluation spanning increasing levels of generalization we used Xenium eczema skin (3 patients, 3 samples; held-out samples; in-house data)^116^, NanoString CosMx non-small cell lung cancer (3 patients, 6 samples; held-out donors) obtained from https://brukerspatialbiology.com/products/cosmx-spatial-molecular-imager/ffpe-dataset/nsclc-ffpe-dataset/, Xenium melanoma (1 patient, 1 sample; held-out dataset) obtained from https://www.10xgenomics.com/datasets/xenium-prime-ffpe-human-skin and ISS CARTANA healthy reproductive tract (3 patients, 6 samples; held-out assay; in-house data)^153^ for niche identification (Fig. 2a); Xenium skin (19 patients, 79 samples, ∼1.1M cells; in-house data)^116^ for niche label transfer (Fig. 2b); a 10x Visium embryo z-stack (52 serial sections; in-house data) for transfer learning (Fig. 2c); Xenium developing pancreas (1 patient, 2 samples; in-house data) and STARmap healthy placenta (1 patient, 4 samples) obtained from https://zenodo.org/records/10981713 for cell-type identification (Supplementary Fig. 9d); and Xenium eczema (15 patients, 48 samples, ∼780k cells; in-house data) and Xenium psoriasis (4 patients, 31 samples, ∼360k cells; in-house data) for disease prediction (Supplementary Fig. 9e). For design ablations and cross-assay benchmarking we used paired mouse brain MERFISH and STARmap sections, obtained from https://cellxgene.cziscience.com/collections/0cca8620-8dee-45d0-aef5-23f032a5cf09 and https://zenodo.org/records/8327576, respectively, and annotated against the Allen Mouse Brain Reference Atlas by the original authors (Supplementary Note 1).

### Developing human pancreas

We profiled 12 sections of human fetal pancreas tissue from 7 developmental stages (8, 10, 11, 12, 13, 14 and 17 PCW) using the 10x Genomics Xenium platform. The human embryonic and fetal material was provided by the Joint MRC / Wellcome Trust (Grant # MR/006237/1) Human Developmental Biology Resource (http://www.hdbr.org). 8 donors were profiled in total, with a single donor analyzed at each developmental stage except for 17 PCW, for which 2 independent donors were included (Fig. 3a). The Xenium 5K gene panel (10x Genomics, 1000724) and an additional custom gene panel of 100 genes (10x Genomics, 1000766, Supplementary Table 4) were used. Cells with fewer than 10 transcript counts, a cell area <20 μm^2^ or >200 μm^2^, or nucleus-to-cell area ratio <0.1 or >0.9 were removed. Differentially expressed genes were identified using a Wilcoxon rank-sum test (Adjusted *P* < 0.05, log fold-change > 0.5) (Fig. 3,4 and Supplementary Figs. 12-15).

### Adult human pancreas

Human pancreatic tissues were obtained through the Network for Pancreatic Organ Donors with Diabetes (nPOD), an organ donor program that recovers and distributes pancreas and associated tissues from consented organ donors for research. We profiled 4 sections from 2 healthy adult human pancreas specimens using the 10x Genomics Xenium platform and used them for fetal-to-adult cross-stage comparison. The Xenium 5K gene panel (10x Genomics, 1000724) with an additional custom gene panel of 100 genes were used (10x Genomics, 1000766, Supplementary Table 4). Cells with fewer than 10 transcript counts were removed (Fig. 4g–o).

### Whole-embryo reference

For cross-tissue analysis, fetal pancreas data were integrated with a whole-embryo Xenium dataset^51^ (Fig. 4 and Supplementary Fig. 16).

### Human kidney

We profiled 12 sections of non-tumor kidney from clear cell renal cell carcinoma nephrectomy specimens using the Xenium 5K gene panel: nine untreated sections from five patients and three post-ICB sections from three patients.

#### HST-5K-Corpus

HST-5K-Corpus is the subset of HST-Corpus-112M with a Xenium 5K gene panel. Excluding the whole embryo and pancreas, this corpus was used for cross-tissue analysis. Its composition is described in Supplementary Fig. 1.

#### Preprocessing and cell-type annotation

Analysis datasets were harmonized and quality-controlled as for the pretraining corpus (see “HST-Corpus-112M assembly and harmonization”). Gene-expression matrices were then normalized and log-transformed, followed by PCA, k-nearest-neighbor graph construction and Leiden clustering. Annotation was dataset-specific. For the human kidney, cell types were assigned by label transfer from a public human kidney atlas^57^ using scANVI, resolving 31 cell types across 1,350,215 cells from all 12 sections (nine untreated and three post-ICB). For the fetal pancreas, cell types were annotated by Leiden clustering of the scVI latent space^154^. Epithelial, mesenchymal, and immune populations were then subsetted and re-clustered to resolve finer cellular subpopulations. Following the recovery of an additional 675 cells that had been omitted due to a metadata error, scANVI^63^ was used to transfer cell-type labels from the original annotated atlas to the final atlas of 1,586,341 cells. Analysis of ductal and endothelial subpopulations used TERRA cell embeddings. For the adult pancreas, cell types were annotated using TERRA cell embeddings. Ground-truth niche annotations used for benchmarking were sourced from the original publications or annotated by experts in the case of in-house data. For the archetype analysis, we used a subset of HST-5K-Corpus (Supplementary Fig. 1), excluding all developing tissues: embryonic cells were removed by tissue label, and the pancreas was excluded because the available sections derived from developing tissue whose fetal-like architecture would confound a cross-organ analysis of adult macrophages. After these exclusions the working corpus comprised 32,992,418 non-embryonic cells; niche and archetype construction were performed on this set, with the two developing-pancreas archetypes removed post hoc, leaving ≈28M cells across the nine adult organs in the final analysis. All niche-level analyses (sketching, niche identification, archetype grouping) were performed on the neighborhood embedding; per-cell-type state discovery (below) used the cell embedding. Macrophages were identified using relevant marker genes on the cell embedding, yielding 1,522,556 macrophages across the nine organs, and macrophage-enriched niches were defined and grouped into archetypes as described in the following sections.

### Cellular neighborhood construction and tokenization

For each index cell we defined a cellular neighborhood as the cell together with its k nearest spatial neighbors (k ≤ 10), from a k-nearest-neighbor graph on 2D centroid coordinates (Fig. 1a and Supplementary Fig. 2a). The neighborhood was serialized into an ordered gene-token sequence in three steps (Fig. 1d and Supplementary Fig. 2b): (1) neighbors were ranked by Euclidean distance to the index cell; (2) within each cell, genes were ranked by log-normalized expression; (3) gene tokens from all cells were concatenated in this order. The production tokenizer (cell graph) preserves each neighbor cell as a distinct segment; an alternative tokenizer that collapses the neighborhood into a single summed segment was evaluated as an ablation (Supplementary Fig. 2f–j). Sequences used 256 index-cell tokens and 2,560 neighborhood tokens (2,816 total), with shorter cells zero-padded. During training, the number of sampled neighbors was drawn uniformly per batch (0–10) and tokens outside the sampled neighbors were replaced by padding tokens; during inference, the number of sampled neighbors was fixed to k = 10.

Each gene token comprises four components (Fig. 1d): a gene-symbol token, a value token encoding (shifted-log-normalized) expression abundance, a gene-rank token (gene position within its cell), and a cell-rank token denoting the cell’s position within the neighborhood relative to the index cell. Formally, the input embedding of token i is

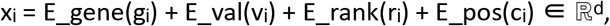

where gᵢ is the gene symbol, vᵢ the (shifted-log-normalized) value, rᵢ the within-cell gene rank and cᵢ the cell-rank/segment index; E_gene is a learnable embedding over the gene vocabulary (|V| = 23,407), E_val encodes the value, E_rank is a fixed sinusoidal embedding of the gene rank, E_pos encodes cell position (a sinusoidal segment embedding or relative spatial coordinates), and d = 384. An optional batch metatoken can be prepended to the sequence to account for sample- and assay-specific technical variation; it is used in TERRA-112M (Fig. 1d and Supplementary Fig. 6). Model-specific settings are listed in Supplementary Table 5.

## Model architecture and configurations

### Self-supervised pretraining and masking

TERRA was pretrained with a joint-embedding predictive architecture (JEPA) objective (Fig. 1e). Let X denote the full token sequence of a cellular neighborhood. Its non-padding gene tokens were partitioned by block masking into a context set C and a disjoint target set T: within each cell (segment), a fraction p of the non-zero gene tokens was assigned to T and the remainder served as context, distributing masking evenly across cells. The released models used a single context block and a single target block, with a per-block masking ratio of p = 0.6 for TERRA-112M and 0.8 for TERRA-96M (Supplementary Table 5).

A context encoder *f* encoded the context-masked view *X _C_*; mask tokens carrying the target positionswere then placed at the target positions and passed, together with the context representation, to a predictor *g*_φ_, which output predicted target representations

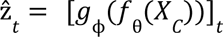 *for t* ∈ *T*. In parallel, a target encoder 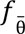 encoded the full (unmasked) sequence to produce the targets 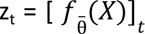. The context encoder and predictor were trained to minimize the L₁ prediction error over the target tokens (metatokens excluded):

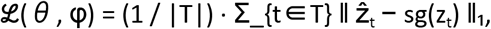

where sg denotes the stop-gradient operator. The target encoder was not updated by gradient descent; its parameters were an exponential moving average of the context-encoder parameters,

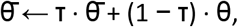

with momentum τ increased from 0.9995 to 1 over training. Here *f* θ and *g* **φ** are the encoder and predictor described in “Model architecture and configurations” (12-layer transformers of width 384 and 192, respectively).

### Multi-scale embedding extraction

At inference, the tokenized neighborhood was passed through the target encoder *f* θ without fine-tuning, with the batch metatoken zeroed (padded) so embeddings are covariate-agnostic. Per-gene-token outputs were aggregated by average pooling into four multi-scale embeddings (Fig. 1a and Supplementary Fig. 2e): cell-gene embeddings (index-cell gene tokens), neighborhood-gene embeddings (a given gene averaged across the neighborhood), cell embeddings (mean over index-cell gene tokens) and neighborhood embeddings (mean over all gene tokens in the neighborhood). Embeddings were visualized in two dimensions with UMAP (scanpy.tl.umap).

### Analysis settings

We use “zero-shot” to mean that the pretrained TERRA model is applied to unseen tissue sections to extract gene-, cell- and neighborhood-level embeddings with all model weights frozen, and that these embeddings are then used directly (for example, by Leiden clustering, label matching, or *in silico* perturbation). Two additional settings are used only where indicated. In supervised transfer, a classifier is trained on frozen TERRA embeddings while TERRA’s weights remain fixed; this includes the linear-probe niche label transfer (Fig. 2b) and the attention-based multiple-instance-learning disease classifier (Supplementary Fig. 9e). In lightweight adaptation, a subset of TERRA’s weights is updated, either by selective fine-tuning of TERRA layers (Fig. 2b) or by low-rank adaptation (LoRA) for transfer to 10x Visium (Fig. 2c). In all cases, the underlying TERRA embeddings are obtained zero-shot; only the supervised-transfer and adaptation settings introduce trained parameters.

### Evaluation metrics

Clustering against reference annotations was measured by NMI and ARI. For the mouse brain cross-technology benchmark, weighted NMI/ARI aggregated per-assay scores weighted by the number of cells in each dataset. Cross-assay integration was measured by the integration local inverse Simpson’s index (iLISI; higher = better mixing) and maximum mean discrepancy (MMD; lower = better mixing) using a multi-bandwidth Gaussian (RBF) kernel. For the niche label transfer benchmark, accuracy and macro-F1 score were computed at the cell level. For the disease classification benchmark, cell-weighted accuracy and AUC-ROC score aggregated per-bag predictions weighted by the number of cells in each bag.

### Ablation experiments

#### Experiment setup

All models used for the mouse-brain design ablations and the cross-assay benchmark (Supplementary Figs. 4–8; Supplementary Notes 1–3) were trained from scratch on the mouse-brain MERFISH and STARmap data. These are separate from the released human models (TERRA-112M and TERRA-96M) and were used only to establish and benchmark TERRA’s architecture under controlled, single-tissue conditions; mouse genes were keyed to their native mouse Ensembl IDs (ENSMUSG) rather than mapped to human orthologs.

#### Model architecture

We ablated tokenizer type, per-cell sequence length, expression encoding, spatial encoding, masking ratio, normalization strategy and the batch metatoken on mouse brain MERFISH and STARmap data (Supplementary Figs. 4-6; Supplementary Notes 1 & 2).

#### Scaling analyses

To characterize how zero-shot performance scales, we trained models across three axes, varying one at a time and holding the others at the default configuration, and evaluated niche identification (Supplementary Fig. 10) and cell-type identification (Supplementary Fig. 11) on the held-out datasets. Model size spanned GT Tiny (encoder dimension 192, 3 attention heads; ∼10M parameters), GT Small (384, 6 heads; ∼30M) and GT Base (768, 12 heads; ∼103M), each a 12-layer encoder–predictor with width and head count scaled with model size. Pretraining-corpus fraction spanned 1%, 10%, 25%, 50% and 100%, obtained by randomly sampling the corresponding fraction of tissue sections and retraining on the sampled sections, with held-out evaluation sections excluded; Masking ratio spanned per-block values of 0.5, 0.6, 0.7, 0.8 and 0.9. For the dataset-fraction and masking-ratio sweeps, model size was held at GT Base and corpus fraction at 100%. For each configuration, NMI and ARI were computed against ground-truth annotations across a sweep of Leiden clustering resolutions; bars show the mean and error bars the standard deviation across resolutions.

### Benchmarking tasks

Niche and cell-type identification. Neighborhood embeddings (niches) or cell embeddings (cell types) were clustered with the Leiden algorithm. The base resolution was chosen so that the number of clusters matched the number of ground-truth classes with >100 cells; NMI and ARI were then evaluated across a local resolution sweep around this value (±0.04 in steps of 0.01).

#### Niche transfer

On annotated tissues, niche labels were predicted in held-out, unseen samples. For Xenium skin (19 patients, 79 samples, ∼1.1M cells), evaluation was conducted through nested cross-validation (four outer folds for testing and one inner fold for hyperparameter selection) stratified by patient. Two training strategies were designed: for linear probe, the entire target encoder was kept frozen, and for selective unfreezing (fine-tuning), the query-key-value projection, output projection, and both feed-forward layers of the last transformer block of the target encoder were unfrozen while all remaining target encoder parameters were kept frozen. Then, a linear classification head was trained with cross-entropy loss on the TERRA neighborhood embeddings. These strategies were benchmarked against zero-shot embeddings from TERRA, CellPLM, Nicheformer, scGPT-spatial, and Novae. For zero-shot embeddings, k-nearest-neighbor classifiers assigned each test cell the majority niche label among its k nearest (by cosine similarity) neighbors in the embedding space. Performance was measured via accuracy and macro-F1 scores on the test partitions (Fig. 2b; bars show mean scores and dots show individual folds).

### Disease classification

On annotated tissues, disease labels were predicted for held-out unseen samples. For Xenium eczema (15 patients, 48 samples, ∼780k cells) and Xenium psoriasis (4 patients, 31 samples, ∼360k cells), evaluation was conducted through nested cross-validation (four outer folds for testing and one inner fold each for hyperparameter selection). Two settings were considered: unseen patient generalization, in which all samples from a given patient were held out together, and unseen sample generalization, in which individual samples were held out. TERRA cell embeddings were aggregated into per-sample bags and passed to a multiple-instance-learning (MIL) classifier from MultiMIL^155^, comprising a shared MLP cell aggregator (fully connected layer with layer normalization, LeakyReLU activation, and dropout), an attention-based pooling module that aggregated per-cell embeddings into a bag-level representation, and a linear classification head. This model was trained with cross-entropy loss on the bag-level representations to predict the disease label. Performance was benchmarked using cell-weighted accuracy and AUC-ROC scores against equivalent MIL classifiers applied to zero-shot embeddings from CellPLM, Nicheformer, scGPT-spatial, and Novae (Supplementary Fig. 9e; bars show mean scores and dots show individual folds).

### Transfer learning to 10x Visium

Because 10x Visium measures expression at the level of spots, each capturing several cells rather than a single cell, we mapped the platform onto TERRA’s tokenization by treating each spot as an index “cell.” A spot’s expression profile was tokenized exactly as a single cell (gene-symbol, value and gene-rank tokens), and its cellular neighborhood was built directly on the spot grid: from the 2D spot centroid coordinates we constructed the k-nearest-neighbor graph and ordered the neighboring spots by distance to the index spot, exactly as for imaging-based data (k ≤ 10; see “Cellular neighborhood construction and tokenization”). Each spot therefore yields a spot embedding and a spot-neighborhood embedding, the Visium analogues of TERRA’s cell and neighborhood embeddings. TERRA was then adapted to a held-out 10x Visium embryo z-stack (52 serial sections) by low-rank adaptation (LoRA), training on 42 sections with pretraining weights frozen and evaluating on 10 held-out sections (Fig. 2c). LoRA adapters (rank r = 16, scaling factor α = 256) were inserted into the query–key–value projection, the output projection, and both feed-forward layers of every transformer block of both the context encoder and the predictor; all pretrained weights were kept frozen and only the adapter parameters were updated, with the target encoder maintained as an exponential moving average (EMA) of the context encoder. For evaluation, spot embeddings were clustered with Leiden and scored by NMI and ARI against the ground-truth annotations, sweeping the Leiden resolution separately for each method to report its best NMI/ARI, and benchmarked against zero-shot TERRA and scGPT-spatial.

### Benchmarking baselines

TERRA was compared against zero-shot foundation models (CellPLM, Nicheformer, scGPT-spatial, Novae) applied without fine-tuning, and task-specific methods trained per dataset (neighborhood gene-expression PCA, NicheCompass, CellCharter, BANKSY with Harmony, GraphST with PASTE). All methods were run with default settings. Foundation models were run with the following versions and checkpoints as per their respective tutorials: Novae v0.1.0 (MICS-Lab/novae-human-0), scGPT-spatial v0.2.2 (scGPT_spatial_v1), Nicheformer v0.0.1 (nicheformer.ckpt), and CellPLM v0.1.0 (20230926_85M). All baseline methods were harmonized to a common cluster number (30) for the mouse brain benchmark. Methods requiring forced integration are annotated: post-hoc embedding alignment (BANKSY + Harmony) and prior spatial-coordinate alignment (GraphST + PASTE).

### Downstream analyses

#### Gene-program identification

Cell-intrinsic gene programs (GPs) and spatial gene programs (SGPs) were defined by clustering cell-gene and neighborhood-gene embeddings, respectively, with Leiden. For identifying fetal pancreas SGPs, the dataset was first filtered to only include cells within epithelial niches (LLD, tip-trunk, acinar-duct, proto-islet, acinar, mesenchyme-acinar). Neighborhood-gene embeddings were computed and a KNN-graph was constructed using the 25 nearest neighbors, followed by Leiden clustering (resolution = 0.3). Each resulting cluster was then sub-clustered (resolution = 0.2), yielding 53 SGPs, comprising 24-169 genes (median 91 genes per SGP). Gene enrichment analysis was performed using GSEApy^156^, and SGP activity scores were calculated using pyUCell^157^ (Fig. 3l). For archetype analysis, neighborhood embedding was used to partition cross-tissue niches, then all the cell gene programs characteristic of archetypes were performed using standard differential gene expression (DEG).

Consensus GPs were identified by intersecting the marker genes of archetype 4 cell GPs at tissue-level for both fibroblasts and macrophages, yielding common cell GPs that are cross-tissue.

#### Ligand-receptor analysis

Intercellular signaling interactions were inferred using CellPhoneDB v5 and its statistical framework, with 1,000 permutations, a minimum expression threshold of 5%, and a significance threshold of P < 0.05^158^. Analysis was performed across all niches and developmental stages. Endocrine alpha and beta cells, together with capillaries were further stratified by developmental stage prior to ligand-receptor analysis (Fig. 4f and Supplementary Table 3).

#### Cross-tissue and cross-stage analysis

Fetal pancreas data were integrated with a whole-embryo Xenium dataset and with adult pancreas in a shared TERRA neighborhood-embedding space (Fig. 4g–o). Cross-tissue niches were identified by Leiden clustering of joint TERRA neighborhood embeddings; relatedness between populations was quantified by the Sinkhorn (entropic optimal-transport) distance between their embedding distributions, computed in GeomLoss (blur = 0.3, scaling = 0.9, p = 2, debias = True) and visualized by PCA.

#### In silico perturbation

Because TERRA operates at gene-token resolution within tokenized cellular neighborhoods, perturbations are performed directly on the token sequence, and the neighborhood is then re-encoded with the pretrained (target) encoder. A perturbation is specified as a set of per-cell edits, each defining a target cell (a specific cell or all cells), a target gene (by Ensembl identifier, mapped to its gene token, or all genes) and a target scale (the index cell or its surrounding neighborhood). We used knockout perturbations, which set the gene’s value token to zero and remove (pad) the corresponding gene token. Edits are applied to the index-cell token block, the neighborhood token block, or both, enabling single-gene perturbations as well as coordinated multicellular perturbations; for example, perturbing a ligand across neighboring cells together with its cognate receptor in the index cell. The edited sequence is re-encoded to obtain perturbed gene-, cell- and neighborhood-level embeddings, and predicted effects are read out by comparing perturbed and unperturbed embeddings, visualized by UMAP and quantified by the shift in cell or neighborhood embeddings and by ranking the most-affected genes.

To quantify perturbation effects, the unperturbed and perturbed token sequences were passed through the same frozen target encoder, and contextualized token embeddings were extracted from the final encoder layer. Perturbation effects were quantified as the distance between unperturbed and perturbed embeddings using the entropic-regularized optimal-transport (Sinkhorn) distance with cost exponent p = 2 (W2). We computed this distance at two complementary resolutions.

At single-cell resolution, each cell yields a cloud of contextualized token embeddings, one per gene token. For every cell we computed the W2 distance between its unperturbed and perturbed token-embedding clouds, with the two datasets aligned cell-by-cell. Three variants were derived: a cell-intrinsic score over the index cell’s own gene tokens encoded under cell-only attention; a spatially contextualized cell score over the index cell’s gene tokens encoded under full neighborhood attention; and a neighborhood score over all gene tokens in the neighborhood. These per-cell scores were projected back onto tissue sections to spatially localize where a perturbation exerts its effect (Fig. 5g).

At population resolution, each cell is represented by a single pooled embedding, derived at the same three levels as above: a cell-intrinsic embedding (the average of the index cell’s gene tokens under cell-only attention), a spatially contextualized cell embedding (the average of the index cell’s gene tokens under full neighborhood attention), and a neighborhood embedding (the mean over all gene tokens in the neighborhood). Cells were grouped by a categorical annotation (either cell type or niche), chosen to match the embedding level (for example, cell-intrinsic embeddings grouped by cell type and neighborhood embeddings grouped by niche) and for each group and each embedding level the sets of pooled cell embeddings under the unperturbed and perturbed conditions were treated as two empirical distributions. The W2 distance between these distributions quantifies the aggregate shift of that population’s representation in response to the perturbation (Fig. 5f), reported relative to housekeeping-gene and random-gene perturbation controls. For each niche, per-sample W2 scores were averaged across samples, excluding sample–niche combinations represented by fewer than 20 cells, to obtain a single aggregate score per niche.

To evaluate whether simulated immune checkpoint blockade (ICB) perturbations recapitulated real treatment effects at the gene level, we compared gene embeddings from several conditions against a ground-truth reference derived from real ICB-treated tissue sections. For each gene, embeddings were collected across all available sections within a condition, L2-normalized, and treated as a point cloud in embedding space. Four conditions were compared: untreated control sections, housekeeping-gene perturbation controls, random-gene perturbation controls (ten independently sampled gene sets, each evaluated separately and then averaged), and model-simulated ICB perturbation (”target”) embeddings. For each gene, we computed the entropic-regularized Wasserstein-2 (Sinkhorn) distance between that condition’s gene-embedding point cloud and the corresponding ground-truth point cloud. The top 100 genes were defined as those showing the largest Sinkhorn distance between the control and ground-truth embeddings, representing the genes most affected by real ICB treatment. We report the mean Sinkhorn distance to ground truth over these top 100 genes for each condition, with the simulated ICB perturbation expected to lie substantially closer to the real-treatment ground truth than the control, housekeeping, and random-gene baselines.

To identify a putative ICB gene signature, we first ranked genes by the magnitude of their Sinkhorn (W2) shift between control and model-predicted ICB gene embeddings, and selected the top 100 most-shifted genes (a ranking distinct from the treatment-affected genes defined above). This candidate set was then intersected with differentially expressed genes (DEGs) between control and real ICB-treated samples, defined as genes with adjusted p-value < 0.05 and |log fold-change| > 1.5. This intersection yielded 23 genes, which we defined as the ICB signature genes.

### Sketch-based niche identification

To perform clustering on the HST-5K-Corpus and identify niches and archetypes, we used a leverage-score-like (based on importance score) sketching strategy adapted from geometric sketching^159^ to build a representative subsample, cluster it, and project the resulting labels back to the full corpus. Like geometric sketching, the aim was a compact subsample that preserves rare cellular neighborhoods otherwise lost to uniform sampling; we implemented this using explicit statistical-leverage scores rather than the plaid-covering algorithm of the original method.

#### Importance scoring and sampling

Within each tissue, we assigned every cell an importance score approximating its squared Euclidean distance from the tissue mean, computed by random projection of the mean-centered neighborhood embedding onto K_PROJ = 100 Gaussian random directions (scaled by 1/√K_PROJ) and taking the squared row norm. We then drew up to N_PER_TISSUE = 500,000 cells per tissue by importance-weighted sampling without replacement (probabilities proportional to this score), preserving rare, atypical neighborhoods that uniform sampling would miss. This yielded a sketch of representative cells across tissues (Fig. 6b).

#### Graph construction and clustering

On the sketch neighborhood embeddings we built a k-nearest-neighbor graph (k = 15, exact/brute-force Euclidean search using cuML *NearestNeighbours*). Edges were weighted with a Gaussian kernel using a per-cell adaptive bandwidth (σ = the median neighbor distance of each cell), the graph was symmetrized by taking the elementwise maximum with its transpose, and self-loops were removed. We clustered the resulting undirected weighted graph with the Leiden algorithm (cuGraph) across a sweep of resolutions (0.3–30), fixing the random seed (SEED = 0).

#### Projection to the full corpus

Niche labels from every clustered resolution were transferred to all non-sketch cells by k-nearest-neighbor majority vote (k = 15) in the neighborhood-embedding space, each query cell receiving the modal label of its 15 nearest sketch cells. Per-cell nearest-neighbor indices were cached so that additional resolutions could be re-projected without recomputing the search.

### Resolution selection and macrophage-enriched niche definition

To choose the clustering resolution in a principled, macrophage-focused way, we quantified how much new compositional structure each successive resolution revealed among macrophage niches. Across consecutive resolutions in the sweep we tracked the flow of macrophages from each parent niche to its child niches at the next resolution, and for every parent with at least MIN_MACS_PER_PARENT = 100 macrophages that split into at least two children (each receiving ≥ MIN_MACS_PER_CHILD = 30 macrophages), we measured the mean pairwise cosine distance between the sibling children’s neighborhood-embedding centroids. Averaged over qualifying parents, this “sibling divergence” curve reports whether a further increase in resolution separates macrophage niches into compositionally distinct siblings (high divergence) or merely fragments existing niches without adding structure (low, flat divergence). We complemented it with the within-niche variance reduction achieved at each split (the fractional drop from parent to mean-child trace of the covariance).

Sibling divergence rose with resolution and then plateaued; we selected the working resolution (resolution = 20), at the onset of this plateau, i.e. the point beyond which additional resolution no longer produced compositionally distinct macrophage niches. A niche at this resolution was defined as macrophage-enriched if it contained at least 30 macrophages and at least 2,500 cells in total. Each qualifying niche was classified by tissue composition as tissue-private (≥ 70% of its cells from a single organ) or cross-tissue otherwise.

### Cellular-state discovery and niche composition

To describe each niche by the functional states of its constituent cells rather than by cell type alone, we discovered fine-grained transcriptional states within each major cell type using the TERRA cell embeddings, and then summarized each niche by the within-cell-type distribution of these states. Cell types were first grouped into functional super-categories (e.g. T cells, B cells, plasma cells, macrophages, microglia, Kupffer cells, dendritic cells, neutrophils, mast cells, fibroblasts, pericytes, smooth-muscle cells and endothelial cells; tumor and epithelial populations kept as their own categories). For each super-category we pooled its cells across all tissues and clustered them on the cell embedding using a k-nearest-neighbor graph (k = 15) and Leiden clustering, with resolution set by category: 0.3 by default, 0.25 for rare populations (plasma, mast, B cells, dendritic cells, neutrophils) and 0.4 for tumor/epithelial populations. Tissue-resident macrophage categories (microglia, Kupffer cells) and other cross-organ populations were force-included even where they fell below the variance threshold used for display, so that resident-macrophage states were represented in the cross-organ analysis, and the resulting per-category state label was stored for each cell.

We then represented each macrophage-enriched niche by these states. For every qualifying niche (≥ 30 macrophages and ≥ 2,500 cells; n = 345 before pancreas removal) and every cell-type category present with at least 10 cells in that niche, we computed, for each of that category’s states, the fraction of the category’s cells in the niche assigned to that state, that is, the state count divided by the total number of cells of that category in the niche (categories with fewer than 10 cells in a niche were set to zero). Because these fractions are normalized within each cell type, summing to one across the states of a given category, each feature reflects the distribution of states a cell type adopts in a niche, independently of how abundant that cell type is in the niche or the corpus. Concatenating these (cell type, cell state) features across all categories produced a niche × (cell type, cell state) composition matrix, which was the input to archetype clustering. For the niche × cell-type enrichment heatmap (Supplementary Fig. 20b), cell-type composition was shown as log2 fold-enrichment of each cell type’s within-niche fraction over its global marginal frequency across all qualifying-niche cells.

### Archetype definition

Archetypes were defined by clustering niches on their cell-type-state composition. To make archetypes comparable across organs, we restricted the composition matrix to cell-type categories that recur across tissues (T cell, B cell, plasma, mast, dendritic cell, neutrophil, macrophage, fibroblast, pericyte, smooth-muscle cell and endothelial cell), dropping niches left with no signal in this cross-organ feature space. Each niche vector was L2-normalized (so that Euclidean distance reflects cosine similarity of composition), and niches were clustered by Ward-linkage hierarchical clustering on the Euclidean distances between L2-normalized vectors. The number of archetypes was chosen by scanning K = 3–20 and selecting the K with the highest silhouette score (cosine) that produced no singleton clusters, giving K = 18; archetypes were then re-indexed by size (largest first). Per-archetype mean composition was taken as the mean feature vector over member niches. Each niche was additionally annotated by tissue composition as tissue-private (≥ 70% from one organ) or cross-tissue, and archetypes recurring across multiple organs were designated cross-tissue archetypes.

Two archetypes composed almost entirely of developing-pancreas niches (> 90% pancreas by tissue composition) were removed post hoc, leaving 16 cross-tissue archetypes for all downstream analyses and display.

### Statistical validation of archetypes

To test whether archetype assignment captured structured compositional variation rather than arbitrary partitioning, we performed permutational multivariate analysis of variance^160^ on the niche composition matrix. Pairwise Bray–Curtis dissimilarities between niches were partitioned by archetype label, and the pseudo-F statistic and R² were assessed against a null distribution of N_PERM = 9,999 label permutations (fixed seed). Archetype identity explained 76.2% of inter-niche compositional variance (R² = 0.762, p < 1 × 10⁻⁴). To identify which cell-type-states defined each archetype, we tested every (cell type, cell state) feature for enrichment in each archetype versus all others with a one-sided Mann–Whitney U test, correcting across features within each archetype by the Benjamini–Hochberg procedure (q < 0.05); archetypes were characterized by 5–37 significantly enriched cell-type-states.

### Characterization and validation of the archetype-1 resident-scavenger niche

#### Spatial abundance across conditions

We tested whether archetype-1 abundance changed with skin disease using the section as the replicate unit. For each skin Xenium section with at least 200 cells, we computed the fraction of its cells assigned to archetype 1, and compared these per-section fractions across healthy, psoriatic and eczematous skin by a Kruskal–Wallis omnibus test with pairwise Mann–Whitney U tests (Benjamini–Hochberg-corrected). Archetype-1 abundance did not differ across conditions (Kruskal–Wallis p = 0.11; all pairwise comparisons n.s.).

#### Single-cell validation and the archetype-1 signature

Because the Xenium panels delineate the niche but lack the depth to resolve its program, we used a wide collection of public single-cell RNA-seq atlas of psoriatic and healthy skin^161–180^. Macrophage/monocyte-lineage cells (71,000) were subset, and raw counts were normalized to 10⁴ counts per cell and log1p-transformed (a copy of the raw counts was retained for pseudobulk analysis). We scored each cell for an archetype-1 consensus signature (*MRC1*, *CD163*, *CLEC4M*, *OIT3*, *C5AR1*, *FCGR2A*, *CCL14*, *INMT*) using score.genes by scanpy (ctrl_size = 50), and defined *archetype-1-high* macrophages as the top tercile by this score (score > 66th percentile). We confirmed that the score concentrated in the CCL14hi/MRC1+ scavenger subtypes.

#### Disease-activity gradient

To identify genes tracking disease activity within the niche, we tested for a monotonic increase across ordered conditions (healthy non-lesional → psoriatic non-lesional → psoriatic lesional) at the level of patient pseudobulks. For each donor × condition group with at least 10 cells, we computed the mean log-expression of each gene; genes expressed in a sufficient number of pseudobulks were tested for a monotonic trend with the Jonckheere–Terpstra test (one-sided, increasing), with Benjamini–Hochberg correction across genes. To distinguish archetype-1-specific rewiring from changes shared by all macrophages, we ran the same trend scan in archetype-1-high and archetype-1-low macrophages separately and retained as *archetype-1-specific* those genes with a significant increasing trend (BH q < 0.05, positive delta) in archetype-1-high but not archetype-1-low cells. Gene-program membership (scavenger/lipid, inflammatory/IFN, other) was annotated from curated gene sets, and a scavenger-module score was tested for the same trend at the donor level.

#### Composition of the archetype-1 niche

To ask whether the *CCL14hi Macrophages* share of the niche changed with disease, we computed, per donor and condition, the fraction of that donor’s archetype-1-high macrophages annotated as *Mac2_CCL14hi* (donors with ≥ 10 archetype-1-high macrophages retained). We tested the ordered trend (healthy non-lesional → psoriatic non-lesional → psoriatic lesional) with the Jonckheere–Terpstra test and compared lesional versus post-treatment psoriasis with a two-sided Mann–Whitney U test. The *CCL14hi* share rose with disease activity (Jonckheere z = 2.15, p = 0.016) and remained elevated after treatment (lesional vs post-treatment n.s.).

### Characterization of the archetype-4 ECM-remodeling niche

#### mCAF identification in an ovarian atlas

To resolve the fibroblast identity of archetype 4, we scored cells of an independent ovarian cancer single-cell atlas^128^ with the archetype-4 cell-type-state gene programs (top 100 marker genes per program, using score.genes() by scanpy, ctrl_size = 50), and compared archetype-4 fibroblasts in our corpus against the atlas’s annotated CAF subsets (mCAF, iCAF, starCAF, vCAF). Archetype-4 fibroblasts were selectively enriched for the mCAF (matrix CAF) signature (Cohen’s d > 1.0, BH q < 0.05).

#### Conserved cross-cancer program (pseudobulk)

We tested whether the archetype-4 program was reproduced across cancers by comparing archetype-4 to archetype-1 cells in four matched malignancies [ccRCC (kidney), LUAD (lung), pediatric ovarian adenocarcinoma and melanoma (skin)], separately in fibroblasts and macrophages. For each cell type we took the top 70 conserved archetype-4 marker genes and, within each cancer, computed per-cell mean expression over these genes and compared archetype-4 versus archetype-1 cells by a one-sided Mann–Whitney U test, with Cohen’s d and AUROC as effect sizes; per-cancer p-values were combined across well-powered cancers (≥ 100 archetype-4 cells) by Stouffer’s method (weighted by √n). The fibroblast program was enriched across all four cancers (mean Cohen’s d = 1.31, mean AUROC = 0.80, Stouffer p < 10⁻³⁰⁰); the macrophage program was enriched in ccRCC, LUAD and ovarian adenocarcinoma (mean Cohen’s d = 1.51, mean AUROC = 0.84), with a weaker but concordant trend in melanoma (Cohen’s d = 0.40, p = 5.8 × 10⁻³, n = 27).

#### Similarity to tumor-associated macrophage states

We compared archetype-4 macrophages to reference tumor-associated macrophage states from a pan-tissue myeloid atlas^143^ using the Sinkhorn divergence (entropy-regularized optimal transport; SamplesLoss from *geomloss*, blur ε = 0.05, scaling 0.9). Cells were embedded in a common 50-dimensional PCA space computed on 3,000 highly variable genes pooled across archetype-4 and atlas cells, with Harmony^181^ batch correction; divergences were estimated over 200 bootstraps (2,819 archetype-4 and 1,000 atlas cells per iteration; archetype-4 cells sampled per tissue in quotas of up to 1,000 across kidney, lung, ovary and skin). Archetype-4 macrophages were closest (lowest divergence) to resident *C1QC+* and *LYVE1+* macrophage states.

#### Diffusion pseudotime

To ask whether the archetype-4 program could be recovered within each cell type separately, we ordered ccRCC fibroblasts (n = 14,138) and macrophages (n = 7,461) independently along a diffusion pseudotime computed on their TERRA cell embeddings (using *scanpy* diffusion map and *sc.tl.dpt()*). Archetype-4 consensus genes were expressed along both trajectories.

#### Joint spatial domains

To define spatial domains from both niche partners simultaneously, we first clustered archetype-4 fibroblasts and macrophages into transcriptional states within the two ccRCC sections. For each cell type we re-embedded the cells from raw counts using rapids-singlecell^182^ (normalization to 10⁴ counts, log1p, 2,500 highly variable genes, scaling, 30-component PCA, Harmony batch integration over slide, and Leiden clustering at resolution 0.6 on a 15-nearest-neighbor graph). We then defined spatial domains by neighborhood composition: for every archetype-4 cell, we built a fingerprint by counting the fibroblast and macrophage states of all archetype-4 cells within a 75 µm radius (spatial *cKDTree* query), standardized these composition vectors, and clustered them with k-means. The number of domains was swept over k = 3–8 and set to k = 5 for the final analysis.

#### Survival analysis (TCGA-KIRC)

We tested the prognostic value of the archetype-4 program in the TCGA kidney renal clear-cell carcinoma (KIRC) bulk RNA-seq cohort^183^. Each tumor was assigned an archetype-4 score as the mean, across the top 40 archetype-4 consensus genes, of each gene’s expression z-scored across patients (33 of the 40 genes were present in the cohort); z-scoring gives every gene equal weight in the score. Overall survival was modeled with a univariate Cox proportional-hazards regression (lifelines *CoxPHFitter*) on the continuous score, with no additional covariates, and the association was visualized by Kaplan–Meier curves stratified at the median score with a log-rank test. A higher archetype-4 score was associated with shorter overall survival (univariate Cox HR = 1.48 per unit of the z-scored score, 95% CI 1.12–1.95, p = 0.005; median-split log-rank p = 3.2 × 10⁻²). Because this analysis was unadjusted, the association should be interpreted as a prognostic correlation rather than evidence of stage-independent prognostic value.

#### Validation in an extended tumor-boundary cohort

To test whether archetype 4 recurs beyond the two index sections, we examined all ccRCC sections sampled at the tumor boundary (seven sections, each from a different patient; pre-treatment). For each section we computed the fraction of cells assigned to archetype 4 and compared tumor-boundary against tumor-core sections. Because the archetype 4 program was essentially undetectable in FFPE sections (median 0.17% of cells vs 16.2% in OCT sections), reflecting reduced sensitivity of that preparation for this stromal program, the comparison was restricted to OCT sections. Archetype 4 abundance was approximately two-fold higher at the tumor boundary than in the tumor core (median 34.3% vs 16.0% of cells; one-sided Mann–Whitney U test, p = 0.019). To assess the matrix-remodeling program directly at single-cell resolution, we scored each archetype 4 cell for the archetype 4 ECM program (*POSTN, COL5A2, COL4A1, COL5A1, SULF1, COMP, SFRP4, IGFBP2, INHBA, MMP11, MMP14, CDH11, MRC2, LRP1*; via scanpy score_genes, ctrl_size = 50). Archetype 4 fibroblasts and macrophages expressed this program across all sections with matched annotations (six of seven; one section lacked single-cell annotations and contributed to the abundance analysis only), with the magnitude and spatial distribution of expression varying between patients (Supplementary Fig. 23).

## Data Availability

A list of datasets used for pretraining is available at: https://docs.google.com/spreadsheets/d/1nzfAsYx1Ja6nh6yj9He5jLmJ5lmphkAt/edit?usp=sharing&ouid=106833492727291477516&rtpof=true&sd=true. A subset of HST-Corpus-112M (excluding proprietary data with dataset IDs < 1000) will be released upon publication. The data used for mouse brain benchmarking and ablations are available at https://cellxgene.cziscience.com/collections/0cca8620-8dee-45d0-aef5-23f032a5cf09 (MERFISH) and https://zenodo.org/records/8327576 (STARmap).

## Code Availability

TERRA is available as a Python package (https://pypi.org/project/terra-st/) and its source code is maintained at https://github.com/Lotfollahi-lab/terra. We additionally provide code used to harmonize and prepare data for pretraining, and reproduce our analyses and benchmarking experiments at https://github.com/Lotfollahi-lab/terra-reproducibility. Our TERRA-112M and TERRA-96M models are released on Hugging Face under https://huggingface.co/lotfollahi-lab/TERRA-112M and https://huggingface.co/lotfollahi-lab/TERRA-96M. Documentation on how to use TERRA including tutorials is provided at https://terra-st.readthedocs.io/en/latest/.

