## Supplementary Material for "Multi-scale modeling of human tissues from spatial transcriptomics with TERRA"

### Supplementary Figures

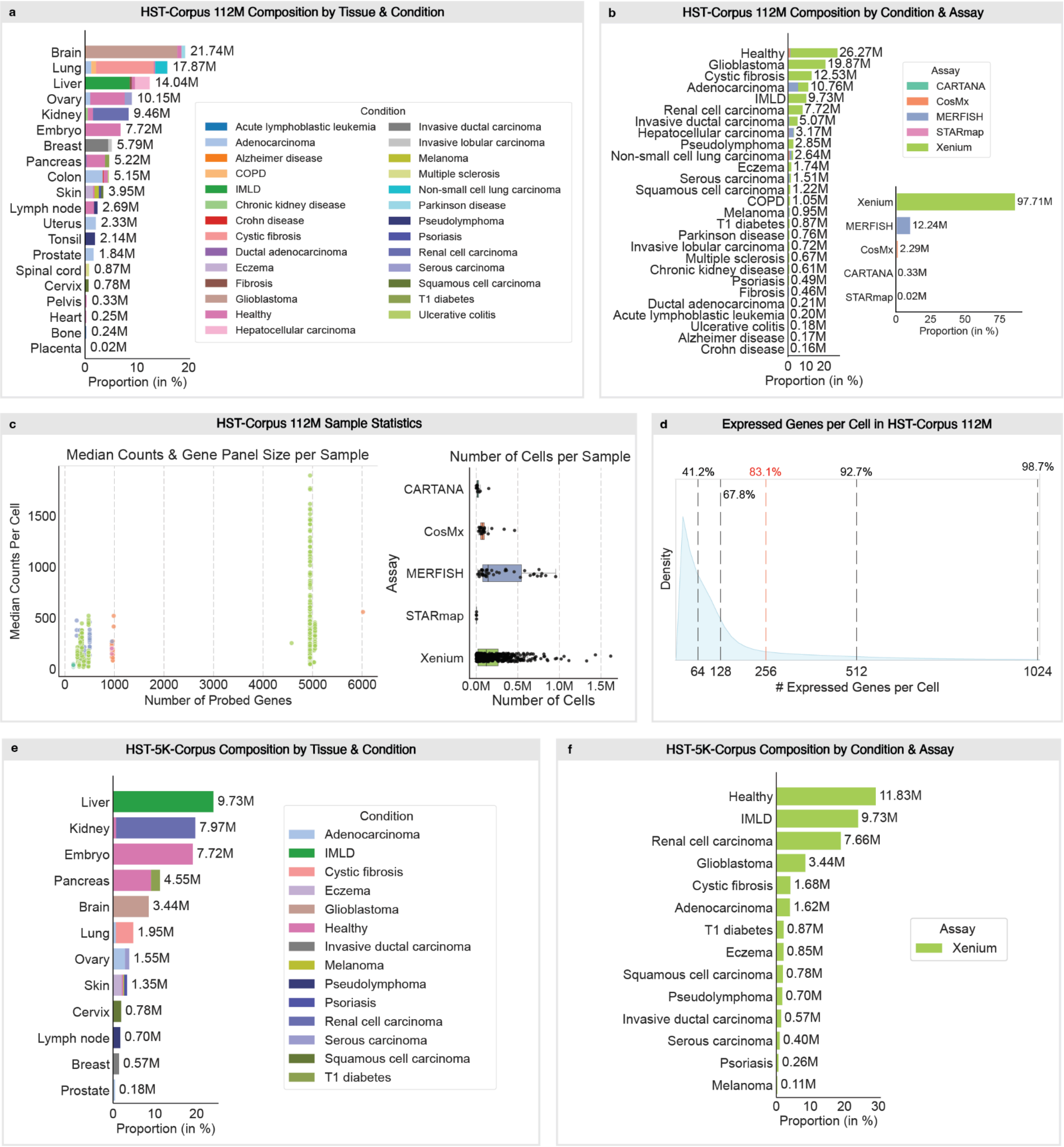


**Supplementary Fig. 1 | HST-Corpus overview**. **a**, Composition of HST-Corpus-112M by tissue (rows) and disease condition (color). The pretraining corpus spans 20 human tissues, with brain (21.74M cells), lung (17.87M) and liver (14.04M) contributing the largest fractions, and rare tissues such as bone (0.24M) and placenta (0.02M) also represented. Absolute cell counts per tissue are shown to the right of each bar; bar segments encode the proportional contribution of each disease condition (legend, right). **b**, Composition by disease condition (left) and by spatial transcriptomics assay (right). Cells originate from healthy tissue and 26 disease indications, most represented by healthy tissue (26.27M cells), glioblastoma (19.87M) and cystic fibrosis

(12.53M). Xenium contributes the majority of cells (97.71M), followed by MERFISH (12.24M), CosMx (2.29M), ISS CARTANA (0.33M) and STARmap (0.02M). **c**, Per-sample technical characteristics of HST-Corpus-112M. Left: median raw transcript counts per cell as a function of probed gene-panel size; each point is one tissue section, colored by assay (same colors as in b). Samples mostly partition into low-plex panels (~300–1,000 genes; ISS CARTANA, CosMx, MERFISH, STARmap, Xenium) and the high-plex Xenium Prime 5,000-gene panel. Right: distribution of cells per sample stratified by assay. **d**, Density distribution of the number of uniquely expressed genes per cell across HST-Corpus-112M. Dashed lines indicate the cumulative fraction of cells with at most 64, 128, 256, 512 and 1,024 expressed genes (41.2%, 67.8%, 83.1%, 92.7% and 98.7%, respectively). The 256-gene threshold (red), corresponding to TERRA's per-cell token capacity, captures 83.1% of all cells fully. **e**, Composition of HST-5K-Corpus, a Xenium-only subset of sections used for cross-tissue analysis and profiled with the Xenium Prime 5,000-gene panel, by tissue (rows) and disease condition (color). Liver (9.73M), kidney (7.97M) and embryo (7.72M) contribute the largest fractions. **f**, Composition of HST-5K-Corpus by disease condition; all cells derive from Xenium 5,000-gene panel sections. COPD, chronic obstructive pulmonary disease; IMLD, immune-mediated liver disease.

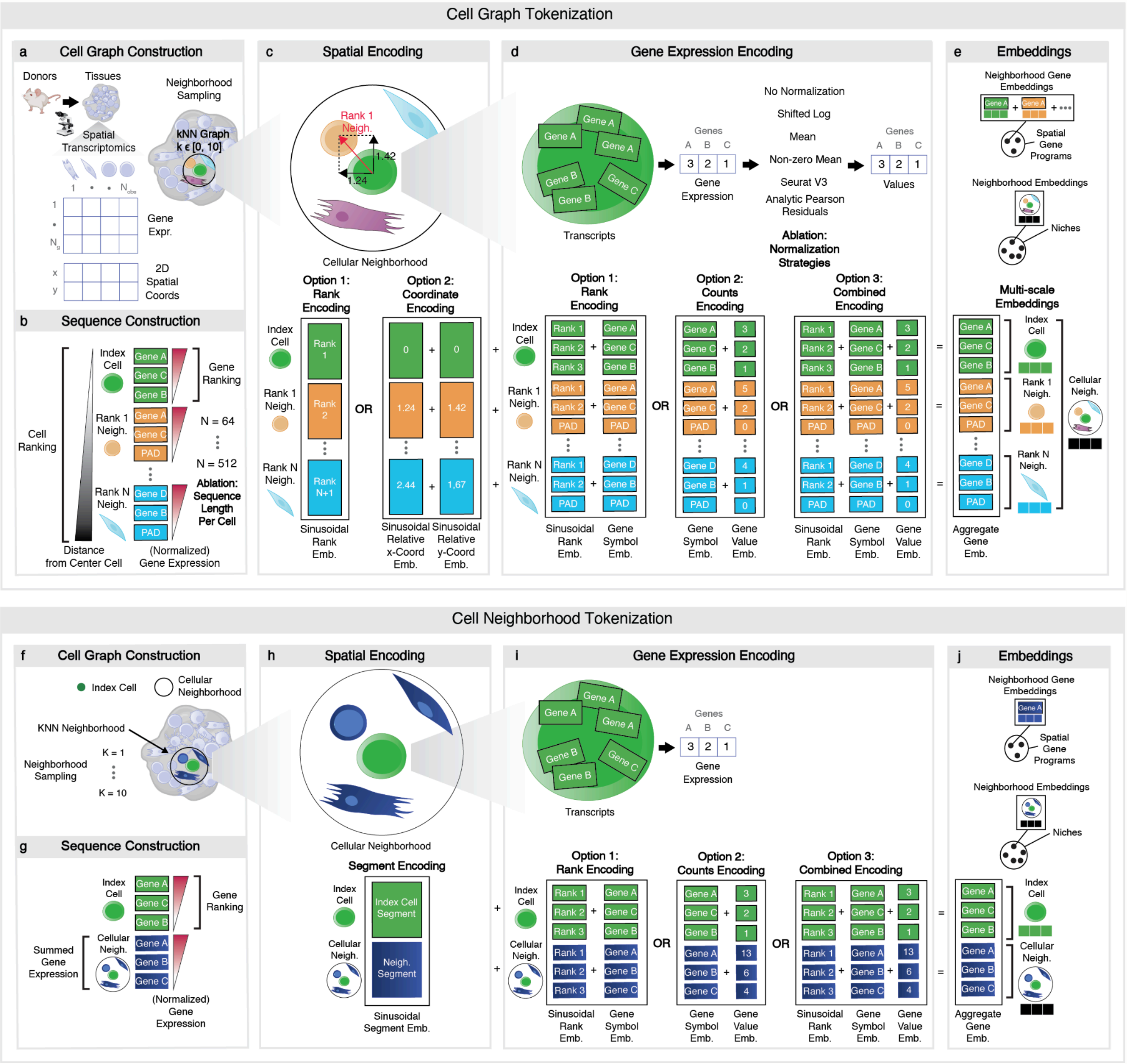


**Supplementary Fig. 2 | Tokenization strategies and ablations**. **a**, Cell-graph construction (production tokenizer): a kNN graph (k ∈ {1, …, 10}) over spatial coordinates defines, for each index cell, a cellular neighborhood. **b**, Sequence construction: neighbors are ordered by distance to the index cell, and within each cell genes are ordered by normalized expression; sequence length per cell N is varied as an ablation (N ∈ {64, …, 512}) with PAD where needed. **c**, Spatial-encoding options: sinusoidal rank embedding (Option 1) or sinusoidal embedding of relative (x, y) coordinates (Option 2). **d**, Gene-expression encoding. Counts are first normalized using one of six strategies (ablation: none, shifted log, mean, non-zero mean, Seurat V3, analytic Pearson residuals), then used in three different ways for tokenization: converted into ranked gene identities (Option 1), encoded and added to their gene identity (Option 2) or encoded, added to their gene identity and combined with a ranking (Option 3; default in TERRA). **e**, The transformer's token-level (gene) embeddings are aggregated into multi-scale representations along two complementary axes. Across genes, the gene embeddings of the index cell are mean-pooled into a cell embedding, and the gene embeddings of the entire neighborhood are mean-pooled into a neighborhood embedding, which is clustered to identify niches. Across cells, the embeddings of a given gene are mean-pooled over the neighborhood into a neighborhood gene embedding, and clustering these identifies spatial gene programs. **f–j**, Alternative cell-neighborhood tokenizer benchmarked as an ablation (Supplementary Note 1). **f**, The kNN neighborhood is collapsed into a single aggregated neighborhood. **g**, Gene expression is summed across the neighborhood and ranked. **h**, A binary segment embedding distinguishes index-cell from neighborhood tokens. **i**, Rank, counts and combined gene-expression encodings analogous to d, applied to summed counts. **j**, Multi-scale embeddings analogous to e. The neighborhood embedding is obtained by aggregating the gene tokens of the neighborhood segment, and neighborhood gene embeddings are the individual gene tokens of that segment.

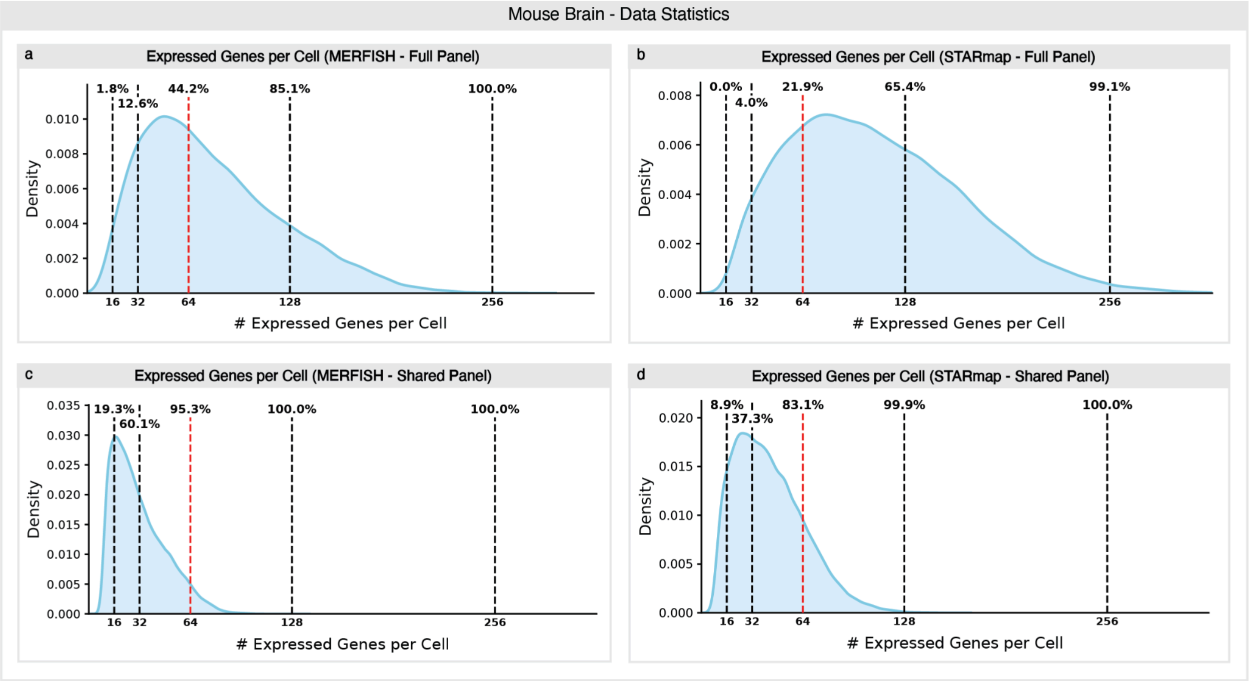


**Supplementary Fig. 3 | Data statistics for the mouse brain data**. Distribution of the number of expressed (non-zero) genes per cell for the MERFISH (1,085-gene) and STARmap (1,015-gene) mouse brain datasets used in the ablations. Dashed lines mark the cumulative percentage of cells expressing at most the indicated number of genes (16, 32, 64, 128, 256), equivalently, the fraction of cells fully represented without truncation at each candidate per-cell sequence length L. **a**, MERFISH, full gene panel: 44.2% of cells express ≤64 genes and 85.1% ≤128. **b**, STARmap, full gene panel: 21.9% ≤64 and 65.4% ≤128. **c**, MERFISH, restricted to the 431 genes shared between assays: 95.3% ≤64. **d**, STARmap, shared panel: 83.1% ≤64. Ablation models are pretrained on the full assay-specific panels (a,b), whereas benchmarking evaluation uses only the shared panel (c,d), where distributions are denser. The 64-gene threshold (red) was used as default for ablations and for benchmarking.

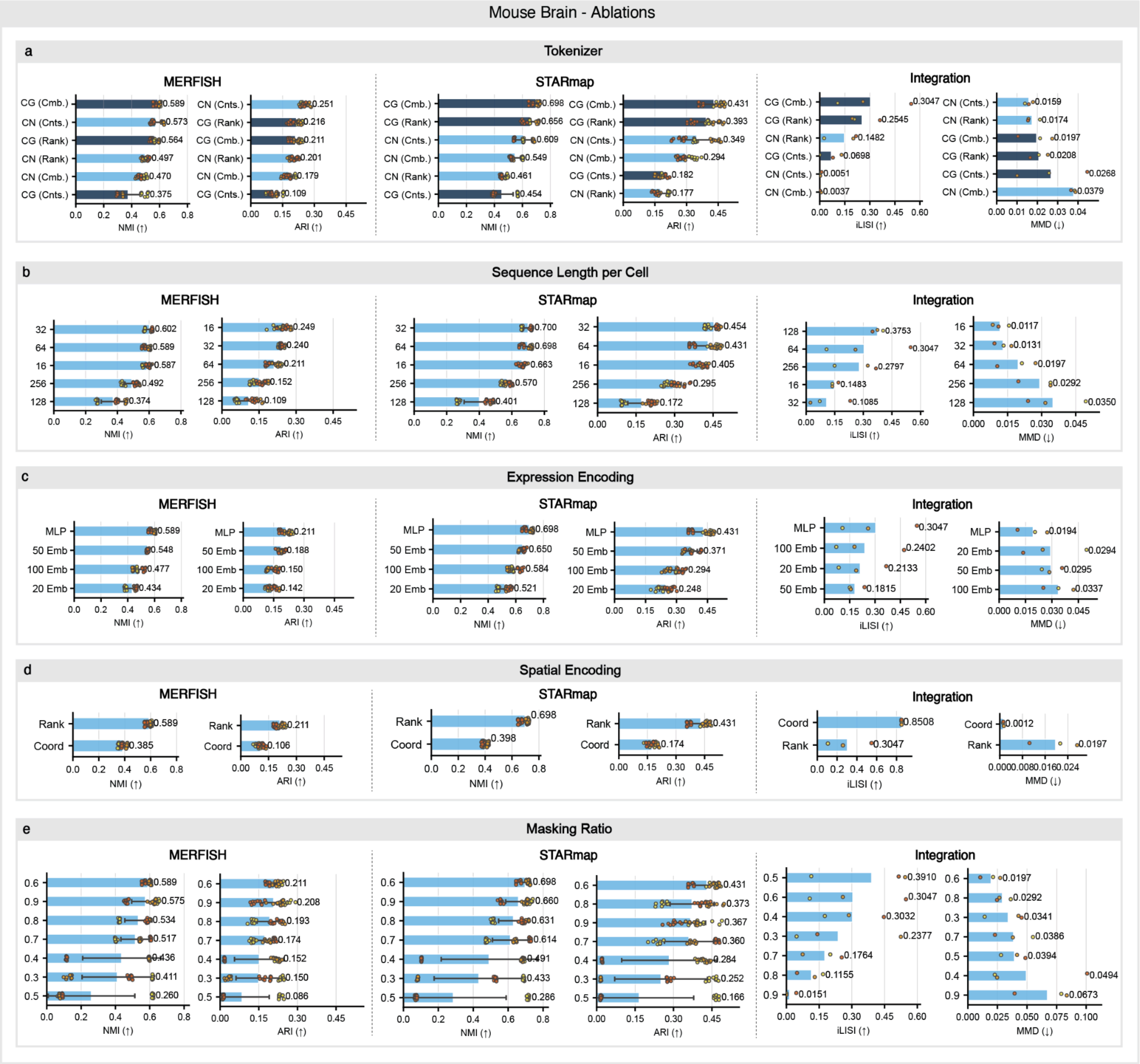


**Supplementary Fig. 4 | Comprehensive ablation sweeps on mouse brain data.** Effect of five design axes on niche identification (NMI and ARI, higher is better) for mouse brain MERFISH and STARmap datasets, and on cross-assay integration (iLISI, higher is better; MMD, lower is better). Bars within each axis are ordered by performance; bars show the mean and points show three training runs (9 varying Leiden clustering resolutions per run for NMI/ARI). In every axis all other settings are held at the default configuration (Cell Graph tokenizer, combined encoding, $L = 64$, MLP expression encoding, rank-based spatial encoding, masking ratio $\rho = 0.6$). **a**, Tokenizer: Cell Graph (CG) versus Cell Neighborhood (CN) tokenization, each with rank-only (Rank), counts-only (Cnts.) or combined (Cmb.) gene encoding. CG with combined encoding gives the best niche identification (MERFISH NMI = 0.589, STARmap NMI = 0.698) and the strongest integration (iLISI = 0.305). **b**, Per-cell sequence length $L \in \{16, 32, 64, 128, 256\}$: $L = 32$–$64$ is optimal for niche identification, whereas $L \geq 128$ degrades performance as most cells express fewer genes and sequences become dominated by low expression/noisy genes. **c**, Expression encoding: continuous MLP projection of counts versus discretized value embeddings (20, 50, 100 dimensions); MLP performs best at all dimensions. **d**, Spatial encoding: rank-based segment embedding versus coordinate-based encoding. Rank encoding is far better for niche identification (MERFISH NMI = 0.589 versus 0.385), whereas coordinate encoding attains artefactually high iLISI (0.851) by aligning cells in coordinate space rather than learning biologically meaningful structure. **e**, Masking ratio $\rho \in \{0.3–0.9\}$: $\rho = 0.6$ is optimal for niche identification; lower ratios ($\rho \leq 0.5$) show high variance across runs (training instability when the prediction task is too easy), while integration improves monotonically as masking decreases. NMI/ARI are computed against Allen Reference Atlas niche annotations; iLISI and MMD quantify MERFISH–STARmap integration. All ablation models were trained on the full gene panel, which is different across the two datasets; inference was performed on the shared gene panel.

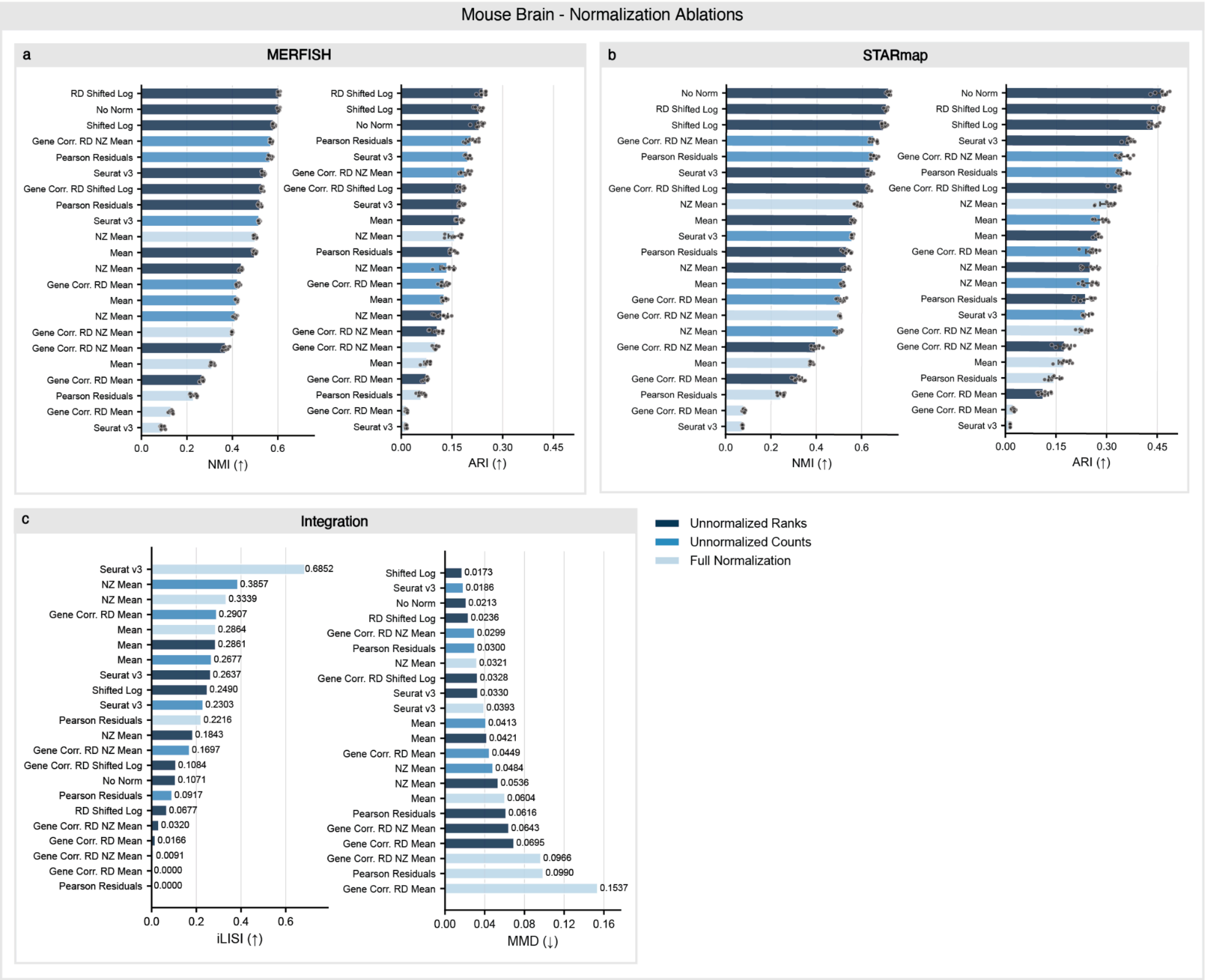


**Supplementary Fig. 5 | Normalization strategies ablation on mouse brain data.** Comparison of gene-expression normalization strategies for the combined (rank + counts) tokenizer, evaluated on mouse brain niche identification and cross-assay integration. Because the tokenizer has two components, normalization can be applied independently to the count component, the rank component, or both; bars are color-coded by category: unnormalized ranks (count component normalized only), unnormalized counts (rank component normalized only) and full normalization (both components normalized). Within each category, individual strategies include no normalization, shifted log, mean, non-zero (NZ) mean, read-depth- and gene-corrected read-depth variants, Seurat V3 and analytic Pearson residuals. Bars show the mean and points show varying Leiden clustering resolutions across one run; bars are ordered by performance within each panel. **a**,**b**, Niche identification for MERFISH (**a**) and STARmap (**b**), measured by NMI and ARI against Allen Reference Atlas annotations (higher is better). Raw values (no normalization / shifted log on raw ranks) rank among the top strategies on both assays, whereas explicit normalizations such as Seurat V3 and Pearson residuals degrade niche identification. **c**, Cross-assay integration, measured by iLISI (higher is better) and MMD (lower is better). Seurat V3 normalization achieves the best integration (iLISI = 0.685) despite its poor niche identification. Together these results indicate that rank-based tokenization already provides implicit normalization; ranking genes by expression is inherently robust to library-size and assay-specific scaling differences so additional explicit normalization is unnecessary and can be counterproductive for niche identification. All ablation models were trained on the full gene panel, which is different across the two datasets, and inference was performed on the shared gene panel.

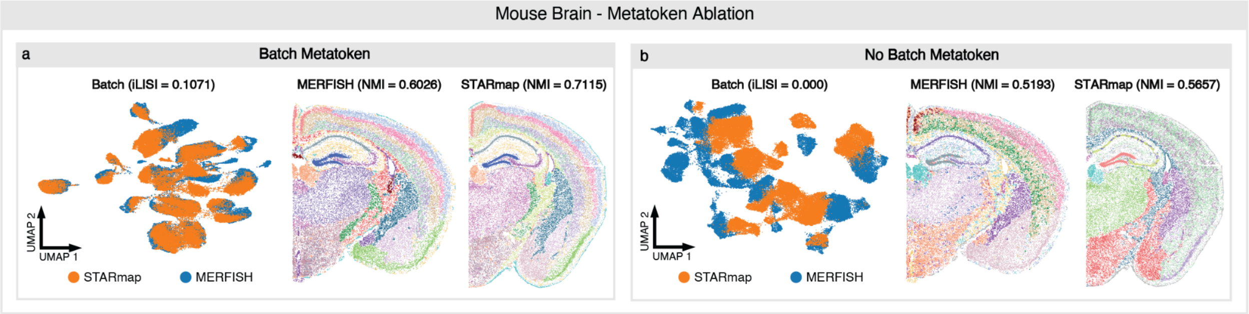


**Supplementary Fig. 6 | Metatoken ablation on mouse brain data. a**, With the batch metatoken, TERRA trained on mouse brain MERFISH and STARmap integrates the two assays in embedding space (left, UMAP of neighborhood embeddings colored by assay; iLISI = 0.107) while resolving anatomically coherent niches in each assay (middle and right, spatial maps colored by Leiden clusters; MERFISH NMI = 0.603, STARmap NMI = 0.712). **b**, Removing the batch metatoken collapses cross-assay integration (iLISI = 0.000) — MERFISH and STARmap cells separate by assay rather than by niche — and degrades niche identification (MERFISH NMI = 0.519, STARmap NMI = 0.566). iLISI, integration local inverse Simpson's index (higher indicates better assay mixing); NMI, normalized mutual information against Allen Reference Atlas niche annotations (higher is better). The metatoken provides a shared structural prior that enables cross-assay integration without explicit batch correction; at inference it is zeroed (padded) so the model produces assay-agnostic embeddings. Models were trained on the full gene panel, which is different across the two datasets, and inference was performed on the shared gene panel.

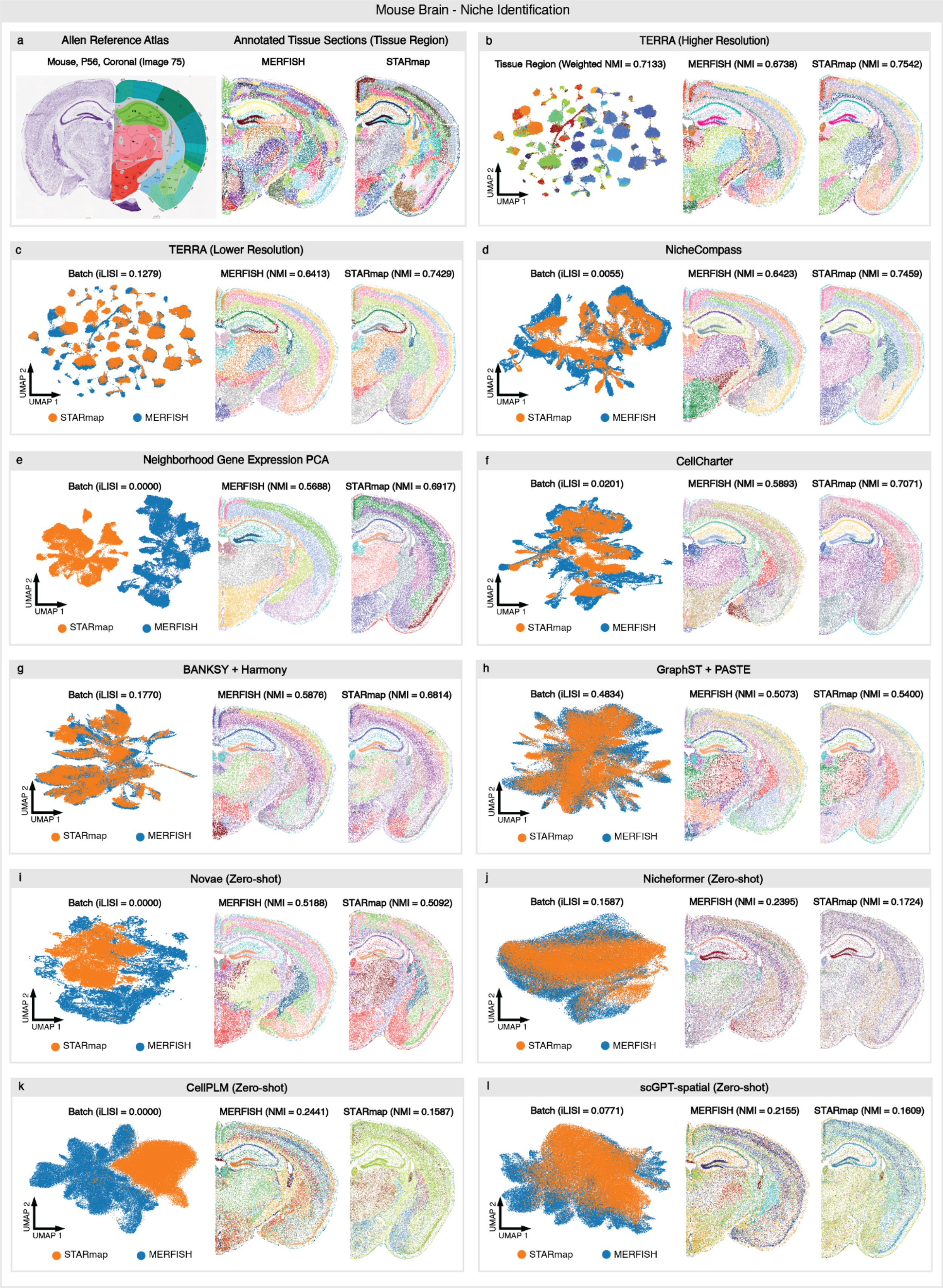

Mouse Brain - Niche Identification
a
Allen Reference Atlas
Annotated Tissue Sections (Tissue Region)
Mouse, P56, Coronal (Image 75)
MERFISH
STARmap
b
TERRA (Higher Resolution)
Tissue Region (Weighted NMI = 0.7133)
MERFISH (NMI = 0.6738)
STARmap (NMI = 0.7542)
UMAP 2
UMAP 1
c
TERRA (Lower Resolution)
Batch (iLISI = 0.1279)
MERFISH (NMI = 0.6413)
STARmap (NMI = 0.7429)
STARmap
MERFISH
d
NicheCompass
Batch (iLISI = 0.0055)
MERFISH (NMI = 0.6423)
STARmap (NMI = 0.7459)
e
Neighborhood Gene Expression PCA
Batch (iLISI = 0.0000)
MERFISH (NMI = 0.5688)
STARmap (NMI = 0.6917)
f
CellCharter
Batch (iLISI = 0.0201)
MERFISH (NMI = 0.5893)
STARmap (NMI = 0.7071)
g
BANKSY + Harmony
Batch (iLISI = 0.1770)
MERFISH (NMI = 0.5876)
STARmap (NMI = 0.6814)
h
GraphST + PASTE
Batch (iLISI = 0.4834)
MERFISH (NMI = 0.5073)
STARmap (NMI = 0.5400)
i
Novae (Zero-shot)
Batch (iLISI = 0.0000)
MERFISH (NMI = 0.5188)
STARmap (NMI = 0.5092)
j
Nicheformer (Zero-shot)
Batch (iLISI = 0.1587)
MERFISH (NMI = 0.2395)
STARmap (NMI = 0.1724)
k
CellPLM (Zero-shot)
Batch (iLISI = 0.0000)
MERFISH (NMI = 0.2441)
STARmap (NMI = 0.1587)
l
scGPT-spatial (Zero-shot)
Batch (iLISI = 0.0771)
MERFISH (NMI = 0.2155)
STARmap (NMI = 0.1609)

**Supplementary Fig. 7 | Cross-assay niche identification benchmarking on mouse brain data.** Comparison of TERRA against task-specific and foundation model methods (zero-shot) for niche identification across two imaging-based assays (MERFISH and STARmap) profiling matched coronal sections of the adult mouse brain. For every method, neighborhood-level representations were clustered into niches and assessed for (i) integration across assays, quantified by the integration local inverse Simpson's index (iLISI; higher indicates better MERFISH–STARmap mixing), and (ii) agreement with ground-truth anatomical annotations, quantified by normalized mutual information (NMI; higher is better) computed separately for each assay. To enable fair comparison, all methods were harmonized to 30 clusters; we additionally show TERRA (higher resolution) which has a more fine-grained clustering to show its strong correspondence with the ground truth anatomy. **a**, Reference annotations: the Allen Mouse Brain Reference Atlas (P56, coronal, image 75) and the corresponding ground-truth niche labels for the MERFISH and STARmap sections. **b**, TERRA at higher resolution (Leiden 1.4); the UMAP of neighborhood embeddings is colored by tissue region (weighted NMI = 0.713), and spatial maps show the identified niches for MERFISH (NMI = 0.674) and STARmap (NMI = 0.754). **c–l**, For each remaining method, the UMAP of neighborhood embeddings is colored by assay (STARmap, orange; MERFISH, blue) with the corresponding iLISI, and the two spatial maps show the identified niches for MERFISH and STARmap with their NMI: **c**, TERRA (lower resolution, 30 clusters); **d**, NicheCompass; **e**, neighborhood gene-expression PCA (expression summed across the neighborhood); **f**, CellCharter; **g**, BANKSY + Harmony; **h**, GraphST + PASTE; **i**, Novae (zero-shot); **j**, Nicheformer (zero-shot); **k**, CellPLM (zero-shot); **l**, scGPT-spatial (zero-shot). TERRA jointly achieved high cross-assay integration and the strongest niche identification, recovering anatomically coherent niches that align with the reference atlas, whereas the zero-shot foundation models (Nicheformer, CellPLM, scGPT-spatial) and several task-specific spatial baselines either failed to integrate the two assays or produced spatially incoherent niches.

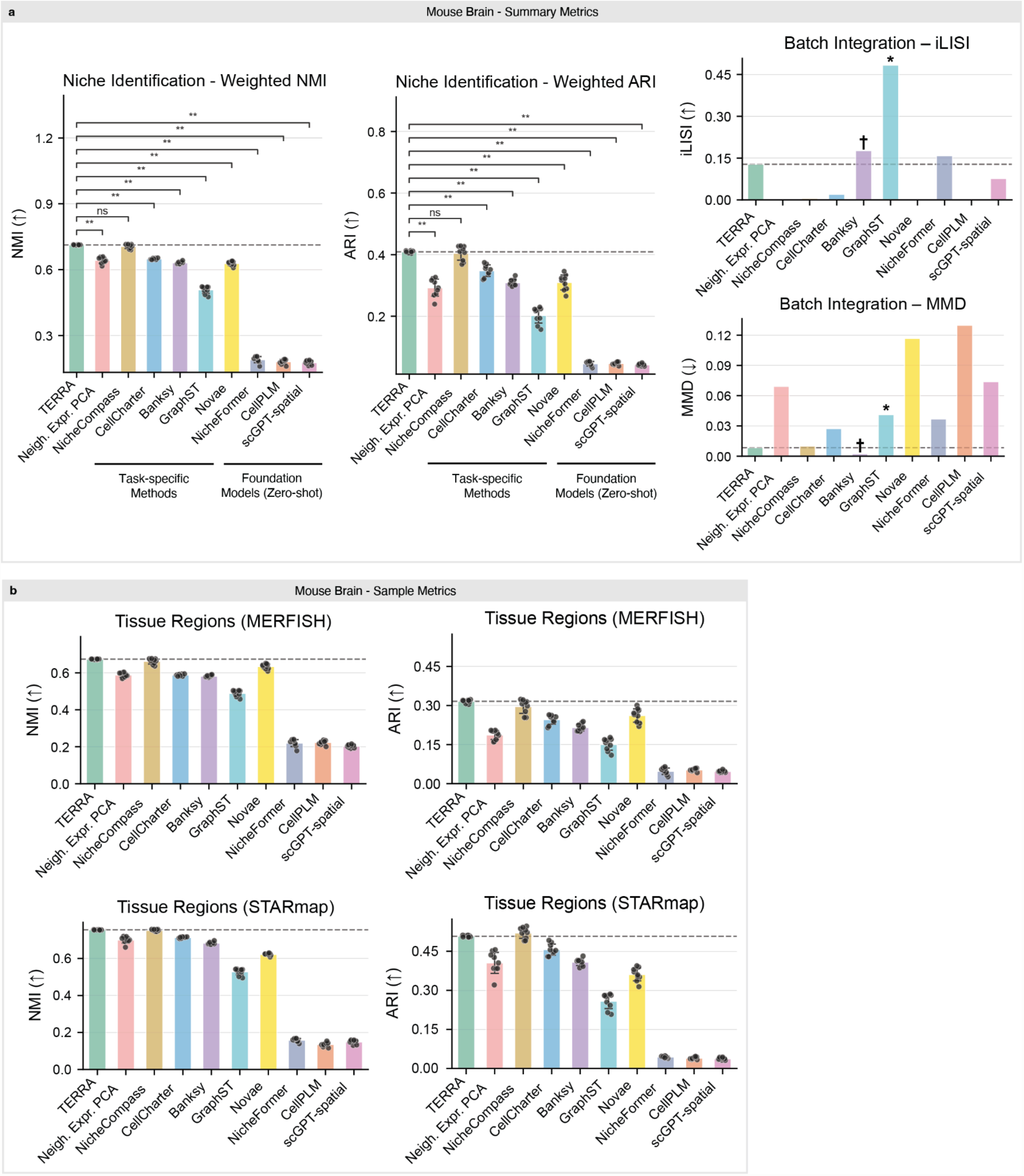


**Supplementary Fig. 8 | Quantitative benchmarking of cross-assay niche identification on mouse brain data.** Summary and per-sample metrics accompanying the qualitative comparison (Supplementary Fig. 7), evaluating TERRA against task-specific spatial methods (neighborhood gene-expression PCA, NicheCompass, CellCharter, BANKSY with Harmony, GraphST with PASTE) and zero-shot foundation models (Novae, Nicheformer, CellPLM, scGPT-spatial) on matched MERFISH and STARmap sections of the adult mouse brain. **a**, Summary metrics. Left, niche identification quantified by weighted NMI and weighted ARI against Allen Reference Atlas annotations (aggregated across both assays; higher is better). Right, cross-assay batch integration quantified by iLISI (higher indicates better MERFISH–STARmap mixing) and maximum mean discrepancy (MMD; lower indicates better mixing). Bars show the mean, dots show varying Leiden clustering resolutions and error bars the s.d.; the dashed line marks TERRA's value as a reference. Brackets denote statistical comparisons between TERRA and each method (two-sided test; **P < 0.01; ns, not significant); TERRA performed better than task-specific methods (while the niche identification performance difference to NicheCompass was not significant, it achieved considerably better integration) and significantly outperformed all zero-shot foundation-model baselines. In the integration panels, † indicates methods that require post-hoc embedding alignment (BANKSY + Harmony) and * indicates methods that require prior spatial-coordinate alignment (GraphST + PASTE); both forcibly integrate samples and therefore hold an inherent advantage on integration metrics while being unable to integrate heterogeneous samples natively. TERRA achieved among the best niche identification together with strong cross-assay integration (high iLISI, low MMD) without any forced alignment. Methods are grouped as our model (TERRA), task-specific methods and zero-shot foundation models. **b**, Per-sample metrics. NMI (left) and ARI (right) for tissue-region identification, computed per section for MERFISH (top) and STARmap (bottom). Bars show the mean across samples, dots show varying Leiden clustering resolutions and error bars the s.d.; the dashed line marks TERRA's value. TERRA matched or exceeded all methods on both assays, whereas the zero-shot foundation models Nicheformer, CellPLM and scGPT-spatial performed markedly worse.

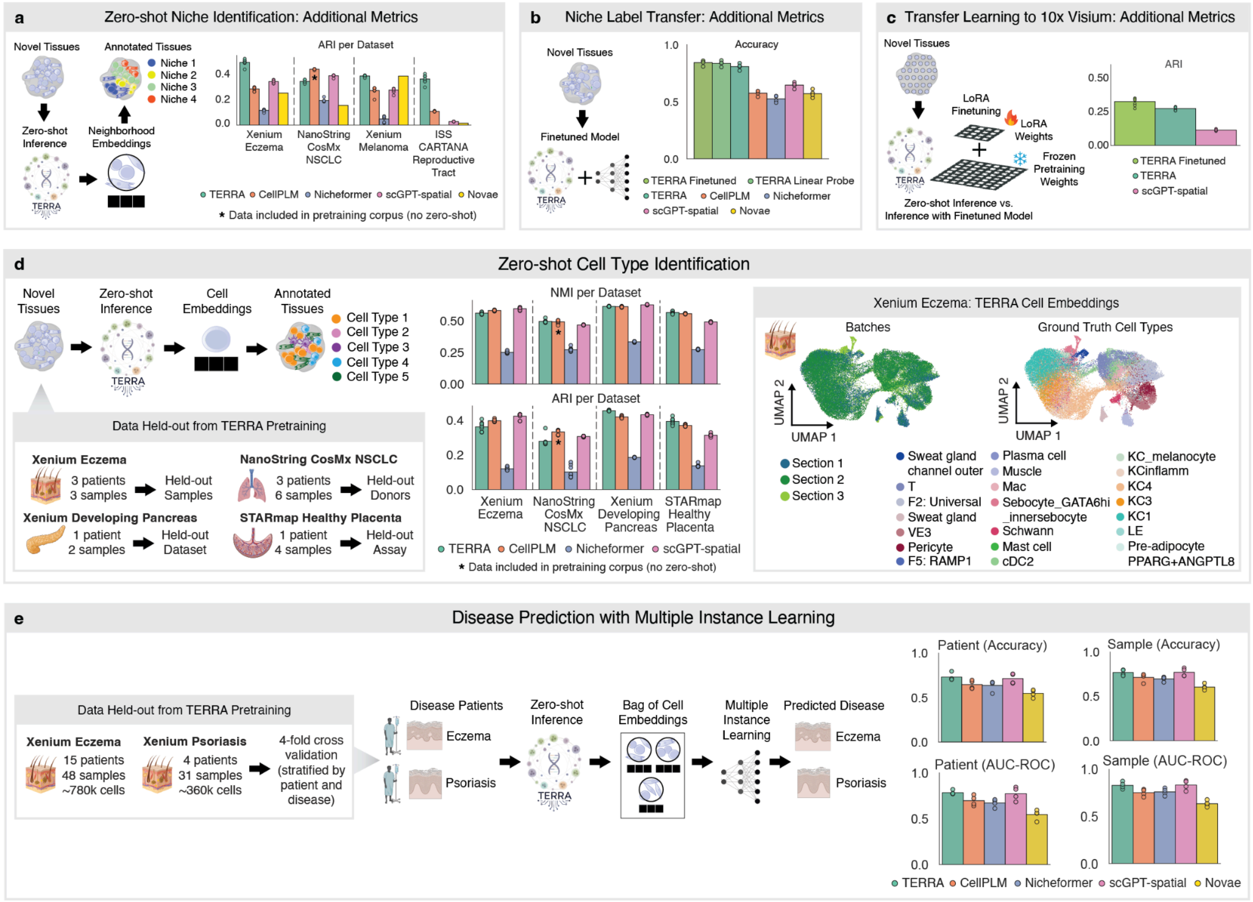


**Supplementary Fig. 9 | Extended benchmarking**. **a-c**, Additional metrics for the benchmarks in Fig. 2. **a**, Adjusted Rand index (ARI) for zero-shot niche identification on the four held-out datasets of Fig. 2a (Xenium eczema, held-out samples; NanoString CosMx NSCLC, held-out donors; Xenium melanoma, held-out dataset; ISS CARTANA healthy reproductive tract, held-out assay), for TERRA and four spatial foundation models (CellPLM, Nicheformer, scGPT-spatial and Novae); bars show the mean and dots individual Leiden clustering resolutions. By ARI, TERRA ranked first on the eczema and reproductive-tract datasets and was on par with the best baseline on melanoma (tied with Novae), whereas on NanoString CosMx NSCLC CellPLM and scGPT-spatial scored higher. The asterisk marks CellPLM on NanoString CosMx NSCLC, whose pretraining corpus included this dataset (not zero-shot). **b**, Cell-level accuracy for niche label transfer on Xenium skin (19 patients, 79 samples, ~1.1M cells; 4-fold cross-validation stratified by patient; Fig. 2b), for TERRA fine-tuned, TERRA linear probe, TERRA (zero-shot), CellPLM,

Nicheformer, scGPT-spatial and Novae; bars show the mean and dots individual folds. **c**, ARI for transfer learning to 10x Visium on the held-out embryo z-stack (LoRA on 42 sections, evaluated on 10 held-out sections; Fig. 2c), for TERRA fine-tuned, TERRA (zero-shot) and scGPT-spatial; bars show the mean and dots individual sections. **d**, Zero-shot cell-type identification on four datasets held out from TERRA pretraining: Xenium eczema skin (3 patients, 3 samples; held-out samples), NanoString CosMx non-small cell lung cancer (3 patients, 6 samples; held-out donors), Xenium developing pancreas (1 patient, 2 samples; held-out dataset) and STARmap healthy placenta (1 patient, 4 samples; held-out assay). Left, schematic of the evaluation workflow: pretrained TERRA was applied directly to each held-out tissue without fine-tuning to obtain cell embeddings, which were clustered with Leiden and matched to ground-truth cell-type annotations. Middle, normalized mutual information (NMI, top) and adjusted Rand index (ARI, bottom) between predicted clusters and ground-truth cell types, per dataset, comparing TERRA against CellPLM, Nicheformer and scGPT-spatial; bars show the mean and dots individual Leiden clustering resolutions. TERRA performed on par with CellPLM and scGPT-spatial and better than Nicheformer. Right, UMAP of TERRA cell embeddings on Xenium eczema, colored by tissue section (left) and by ground-truth cell type (right), illustrating batch integration and cell-type resolution under zero-shot inference. **e**, Disease prediction by multiple-instance learning (MIL) using TERRA cell embeddings on Xenium eczema (15 patients, 48 samples, ~780k cells) and Xenium psoriasis (4 patients, 31 samples, ~360k cells), evaluated by 4-fold cross-validation stratified by patient and disease under two settings: unseen-patient and unseen-sample generalization. Left, schematic: for each specimen, cell embeddings formed a bag passed to an attention-based MIL classifier that predicted eczema versus psoriasis. Right, cell-weighted classification accuracy (top) and AUC-ROC (bottom) under patient (left) and sample (right) splits, for TERRA against CellPLM, Nicheformer, scGPT-spatial and Novae; bars show the mean across folds and dots individual folds.

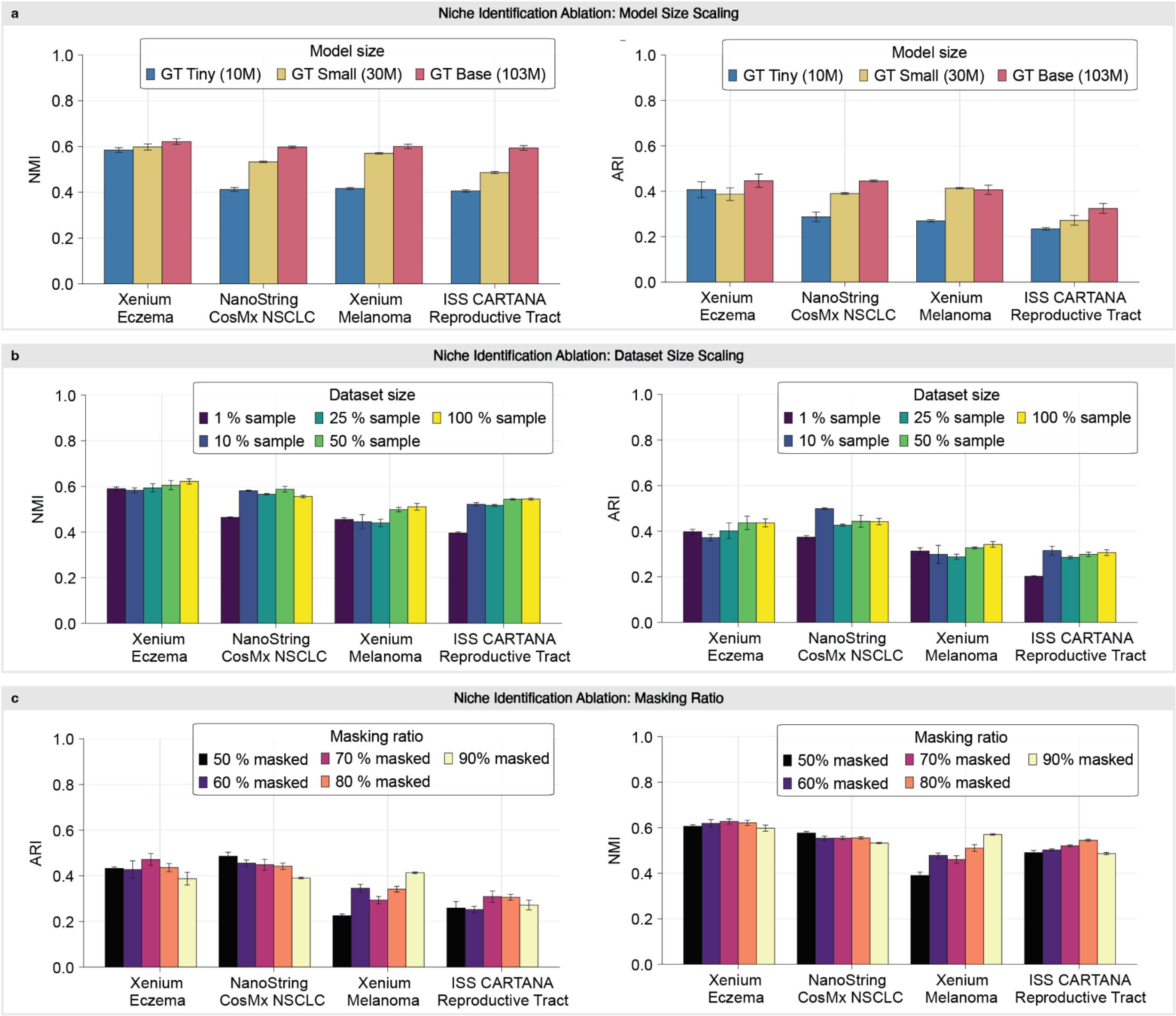


**Supplementary Fig. 10 | HST-Corpus-112M niche identification ablations.** Zero-shot niche identification performance for models pretrained on HST-Corpus-112M (NMI, left; ARI, right) across four held-out datasets: Xenium eczema, NanoString CosMx NSCLC, Xenium melanoma and ISS CARTANA reproductive tract. Bars show the mean across varying Leiden clustering resolutions and error bars show s.d. **a**, Model-size scaling: GT Tiny (10M parameters), GT Small (30M) and GT Base. **b**, Pretraining dataset-size scaling: 1%, 10%, 25%, 50% and 100% of HST-Corpus-112M, holding model size at GT Base (103M). **c**, Masking-ratio scaling: 50%, 60%, 70%, 80% and 90%, holding model size and dataset size at GT Base / 100%.

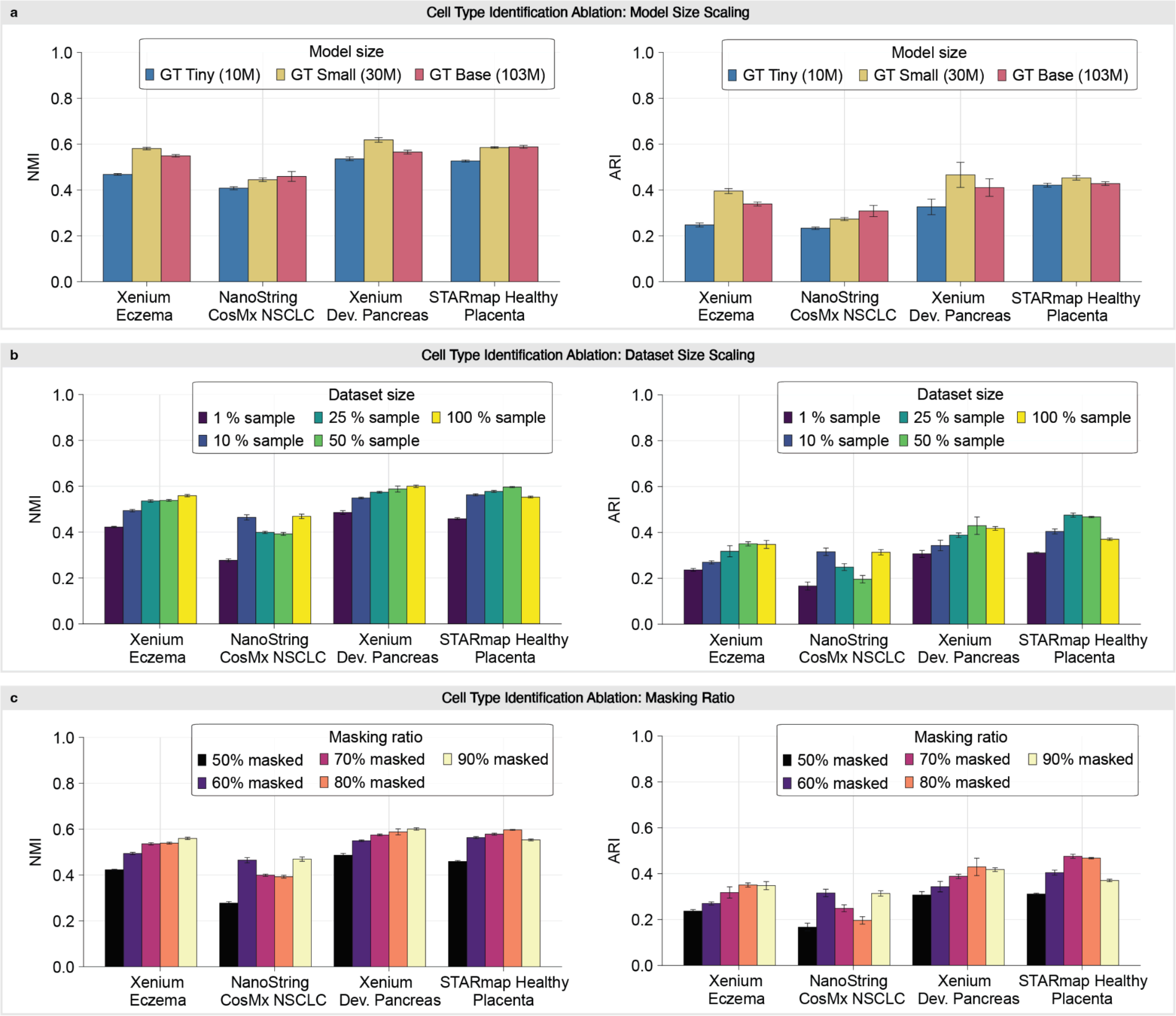


**Supplementary Fig. 11 | HST-Corpus-112M cell-type identification ablations.** Zero-shot cell-type identification performance for models pretrained on HST-Corpus-112M (NMI, left; ARI, right) across four held-out datasets: Xenium eczema, NanoString CosMx NSCLC, Xenium developing pancreas and STARmap healthy placenta. Bars show the mean across varying Leiden clustering resolutions and error bars show s.d. **a**, Model-size scaling: GT Tiny (10M parameters), GT Small (30M) and GT Base (103M). **b**, Pretraining dataset-size scaling: 1%, 10%, 25%, 50% and 100% of HST-Corpus-112M, holding model size at GT Base. **c**, Masking-ratio scaling: 50%, 60%, 70%, 80% and 90% (TERRA default: 60%), holding model size and dataset size at GT Base / 100%.

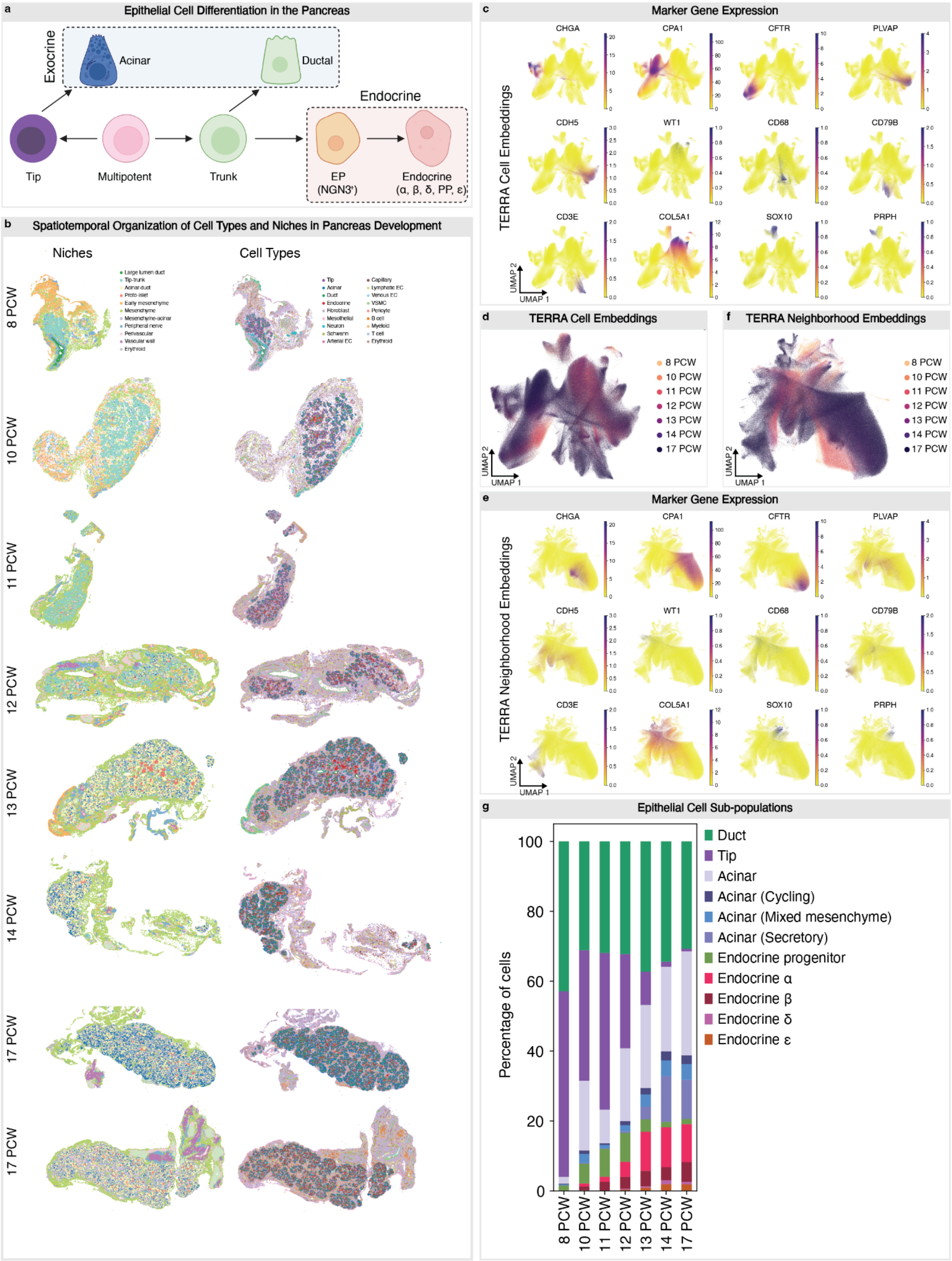


**Supplementary Fig. 12 | A single cell spatial transcriptomic atlas of pancreas development**. **a**, Schematic of pancreatic epithelial cell development. **b**, Niche and cell type organization across the indicated developmental stages. **c**, UMAP of TERRA cell embeddings showing transcript counts of the indicated marker genes. **d**, UMAP of TERRA cell embeddings showing developmental stage. **e**, UMAP of TERRA neighborhood embeddings showing transcript counts of the indicated marker genes. **f**,

UMAP of TERRA neighborhood embeddings showing developmental stage. **g**, Proportions of epithelial subpopulations across developmental stages.

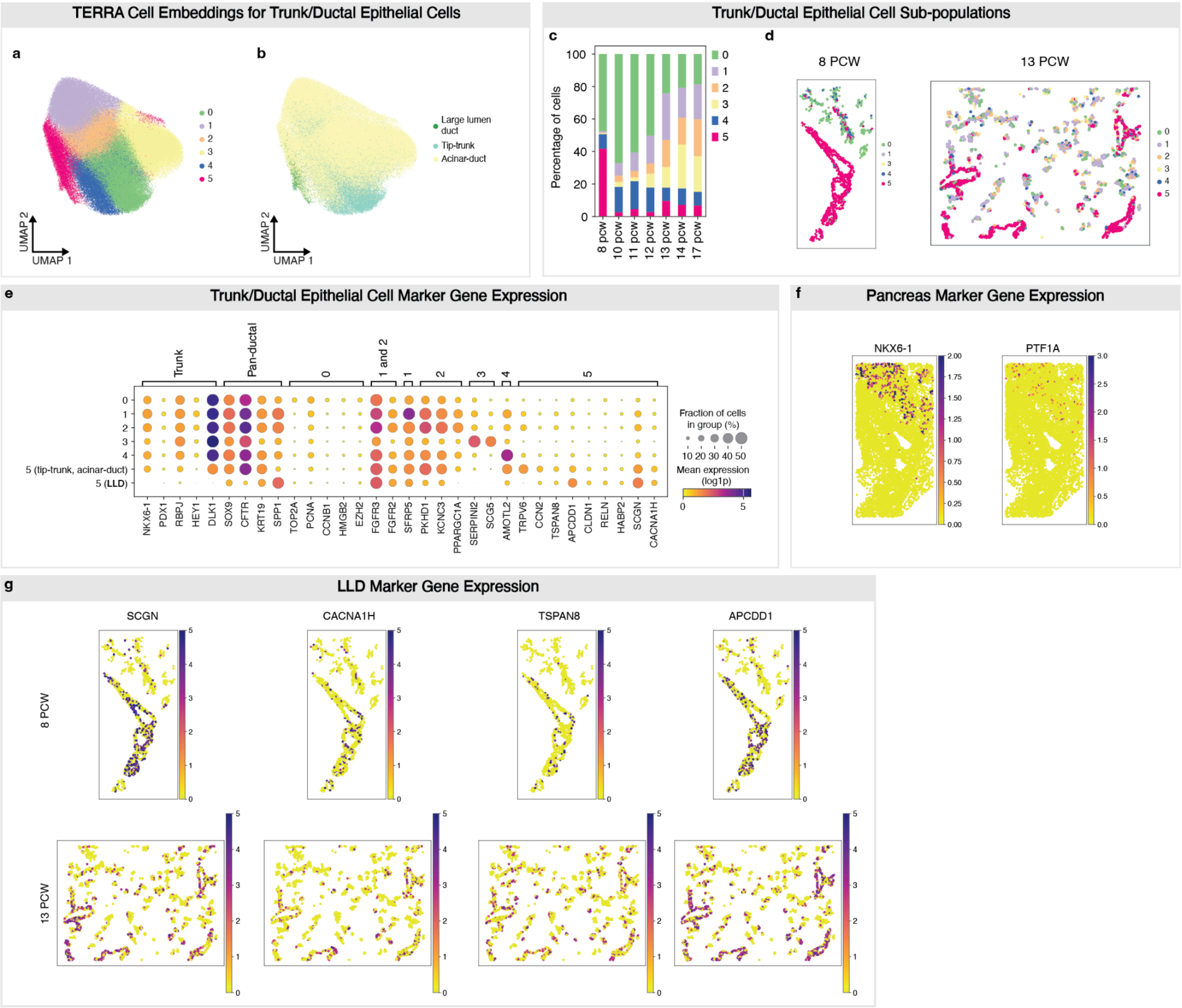


**Supplementary Fig. 13 | Ductal subpopulations during pancreas development**. **a**, UMAP of trunk and ductal epithelial TERRA cell embeddings colored by Leiden cluster. Only trunk and ductal cells residing within the LLD, tip-trunk, and acinar-duct niches were included. **b**, UMAP of trunk and ductal epithelial TERRA cell embeddings colored by niche membership. Only trunk and ductal cells residing within the LLD, tip-trunk, and acinar-duct niches were included. **c**, Proportions of trunk and ductal Leiden clusters across developmental stages. **d**, Spatial organization of trunk and ductal clusters at 8 (left) and 13 (right) PCW. **e**, Dotplot of marker gene expression in trunk and ductal Leiden clusters. Cluster 5 is shown separately for cells residing within the LLD niche or tip-trunk and acinar-duct niches. **f**, Spatial expression (log1p normalized counts) of pancreatic lineage transcription factors *NKX6-1* (trunk regulator) and *PTF1A* (tip regulator) at 8 PCW within the region highlighted in Fig. 3g. **g**, Spatial expression (log1p normalized counts) of the indicated marker genes at 8 and 13 PCW within the regions highlighted in Fig. 3g.

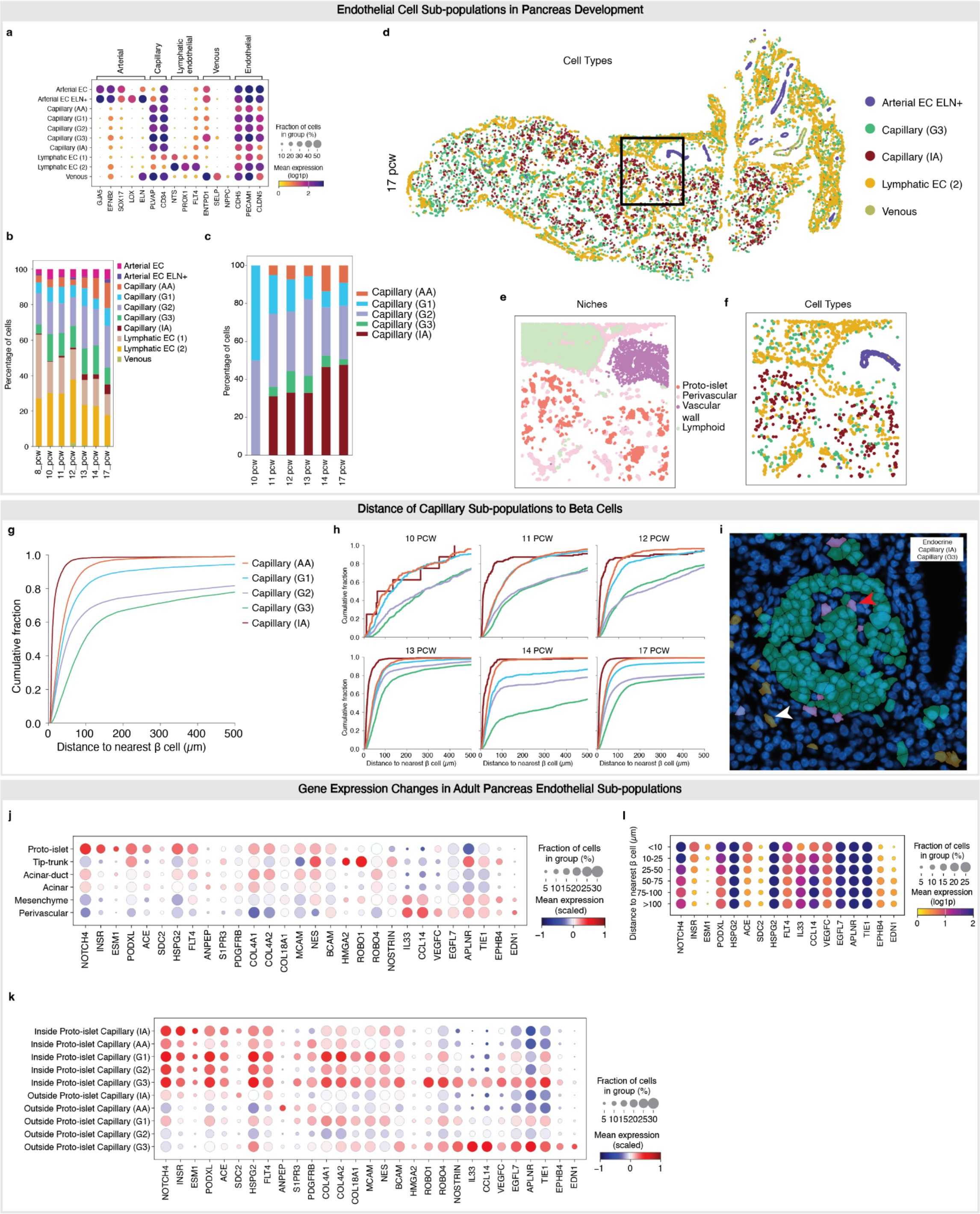


**Supplementary Fig. 14 | Endothelial cell subpopulations during pancreas development**. **a**, Expression of representative marker genes in fetal pancreatic endothelial cell subpopulations. **b**, Proportions of endothelial cell subpopulations across pancreas development. **c**, Proportions of capillary subpopulations within proto-islets across pancreas development. **d**, Spatial organization of the indicated endothelial subpopulations at 17 PCW (donor Sml9). **e**, Magnified view of the region highlighted in **d**, showing the spatial organization of cells in the indicated niches. **f**, Magnified view of the region highlighted in **d**, showing the spatial organization of the endothelial subpopulations shown in **d**. **g**, Cumulative frequency distribution of distances from capillary subpopulations to the nearest endocrine β cell. **h**, Cumulative frequency distribution of distances from capillary subpopulations to the nearest endocrine β cell in each of the indicated developmental stages. **i**, Xenium browser view of sample Hml7 (13 PCW) showing DAPI staining with cell masks overlaid for endocrine cells (α and β cells), IA capillaries, and G3 capillaries. The red and white arrows denote an IA capillary cell and a G3 capillary cell respectively. **j**, Dotplot of marker gene expression in capillary cells stratified by niche membership, irrespective of capillary subpopulation. **k**, Dotplot of marker gene expression in each capillary subpopulation stratified by whether cells reside within or outside of proto-islets. **l**, Dotplot of gene expression in G3 capillaries stratified by distance to the nearest endocrine β cell.

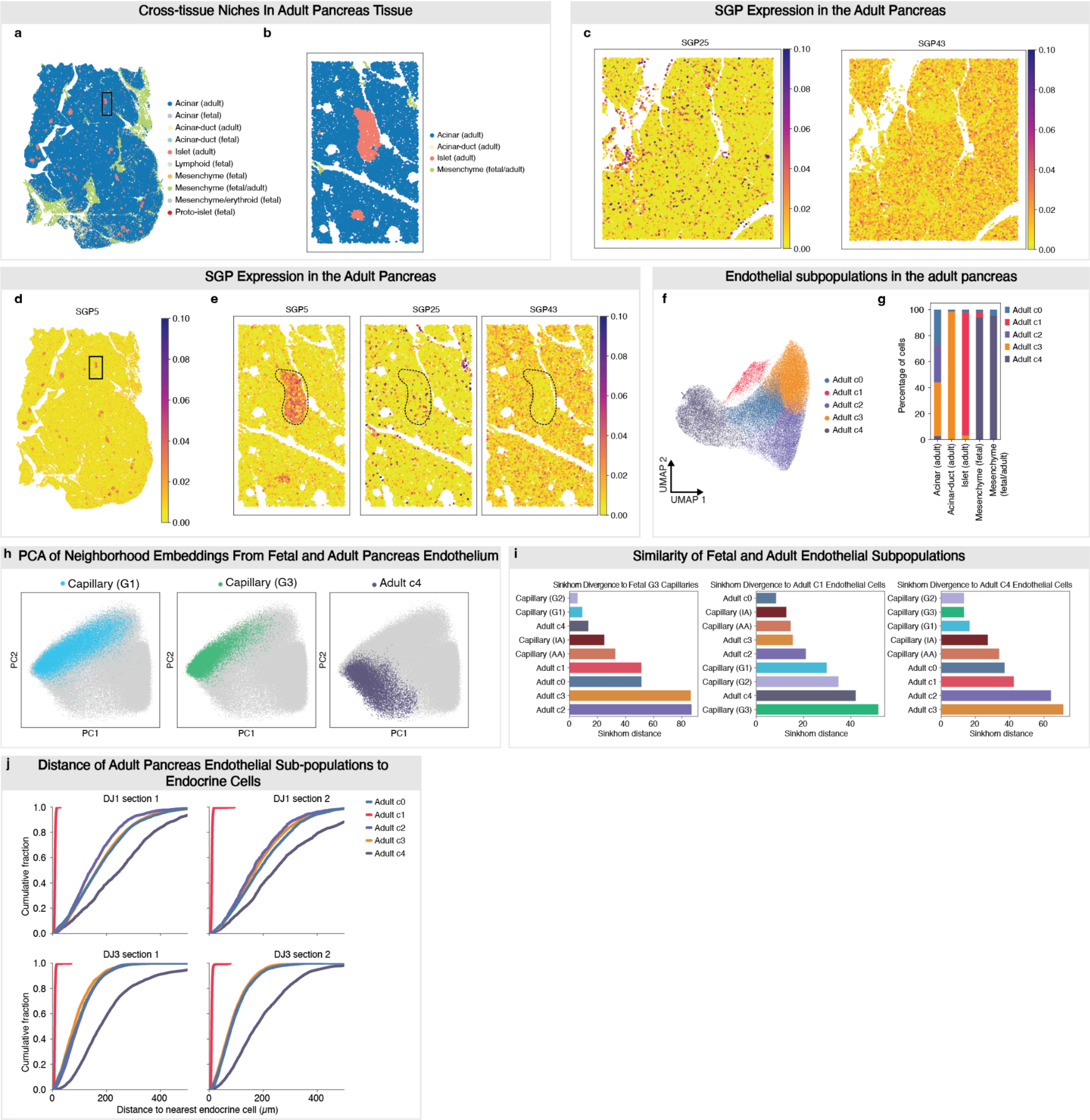


**Supplementary Fig. 15 | Analysis of fetal and adult pancreatic endothelial cells. a**, Niche organization in healthy adult pancreas tissue (donor DJ-3) showing niches from the joint integration of fetal and healthy adult pancreas tissue. **b**, Magnified view of the region highlighted in **a**, colored by the indicated niches. **c,** Magnified view of the region highlighted in main Fig. **4j,** showing expression of SGP25 and SGP43 (donor DJ-1). **d,** SGP expression in healthy adult pancreas tissue (donor DJ-3). **e**, Magnified view of the region highlighted in **d**, showing SGP expression. **f**, UMAP of adult healthy pancreas endothelial TERRA neighborhood embeddings. **g**, Proportions of healthy adult pancreas endothelial cell clusters across niches identified by the joint analysis. **h**,

PCA of fetal and healthy adult endothelial TERRA joint neighborhood embeddings. Fetal pancreas G1 and G3 capillaries are shown, along with adult cluster 4. **i**, Sinkhorn divergence between endothelial subpopulations and the indicated reference subpopulation computed from TERRA joint neighborhood embeddings. **j**, Cumulative frequency distribution of distances from adult endothelial subpopulations to the nearest endocrine cell.

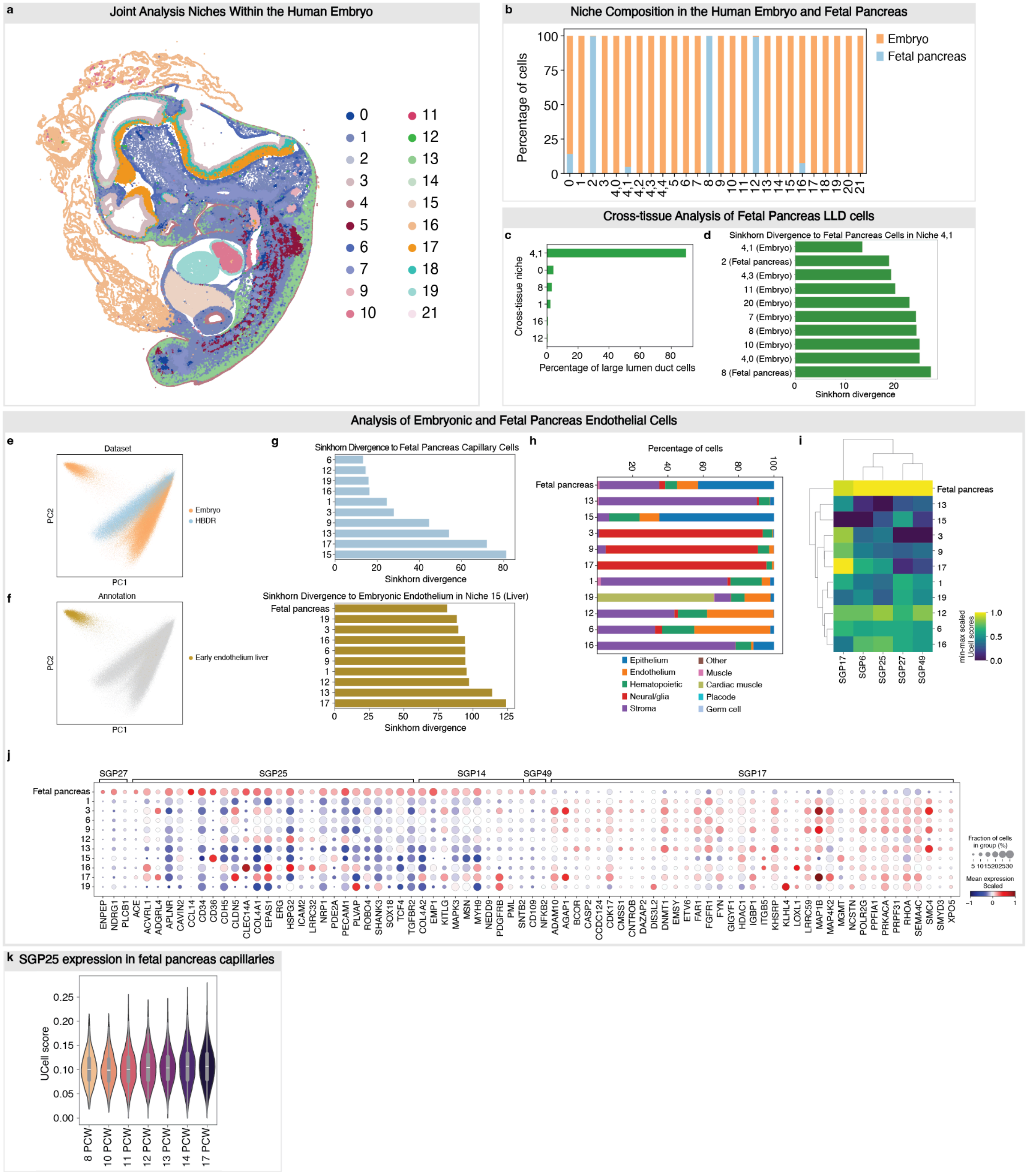


**Supplementary Fig. 16 | Joint analysis of embryo and fetal pancreas endothelial cells. a**, Niche organization from the joint analysis of embryonic and fetal pancreas tissue projected onto an embryonic section (S193). **b**, Proportion of cells from each dataset across niches identified by the joint analysis of embryonic and fetal pancreas tissue. Niche 4 was sub-clustered into 5 additional clusters (4.0-4.4). **c**, Niche membership of fetal pancreas large-lumen duct (LLD) cells in the joint embryonic and fetal pancreas analysis. Only the top 6 niches are shown. **d**, Sinkhorn divergence from fetal pancreas cells in niche 4.1 (containing LLD cells) computed from joint TERRA neighborhood embeddings. Cells were grouped by dataset and niche membership. Only groups containing >1,000 cells were considered, and only the ten most similar groups are shown. **e**, PCA of TERRA joint neighborhood embeddings of embryonic endothelial cells and fetal pancreas capillaries. **f**, Same as **e**, showing embryonic liver endothelial cells. **g**, Sinkhorn distance relative to fetal pancreas capillaries (top) and endothelial cells in embryonic liver niche 15 (bottom). **h**, Cell-type composition of the indicated niches from the joint analysis. Only niches with >1,000 endothelial cells were included. **i**, Heatmap showing min-max normalized SGP expression in endothelial cells in the indicated niches from the joint analysis. Fetal pancreas niches (2, 8, 12) are combined. **j**, Dotplot showing gene expression in endothelial cells across the indicated niches from the joint analysis. Fetal pancreas capillaries from niches 2, 8, and 12 are combined. Genes shown are differentially expressed in fetal pancreas capillary cells relative to embryonic endothelial cells. The genes shown from SGP17 are downregulated in fetal pancreas capillary cells relative to embryonic endothelial cells. Genes shown from SGP6, SGP25, SGP27, and SGP49 are upregulated in fetal pancreatic capillaries. **k**, SGP25 expression (UCell scores) in fetal pancreas capillary cells across developmental stages.

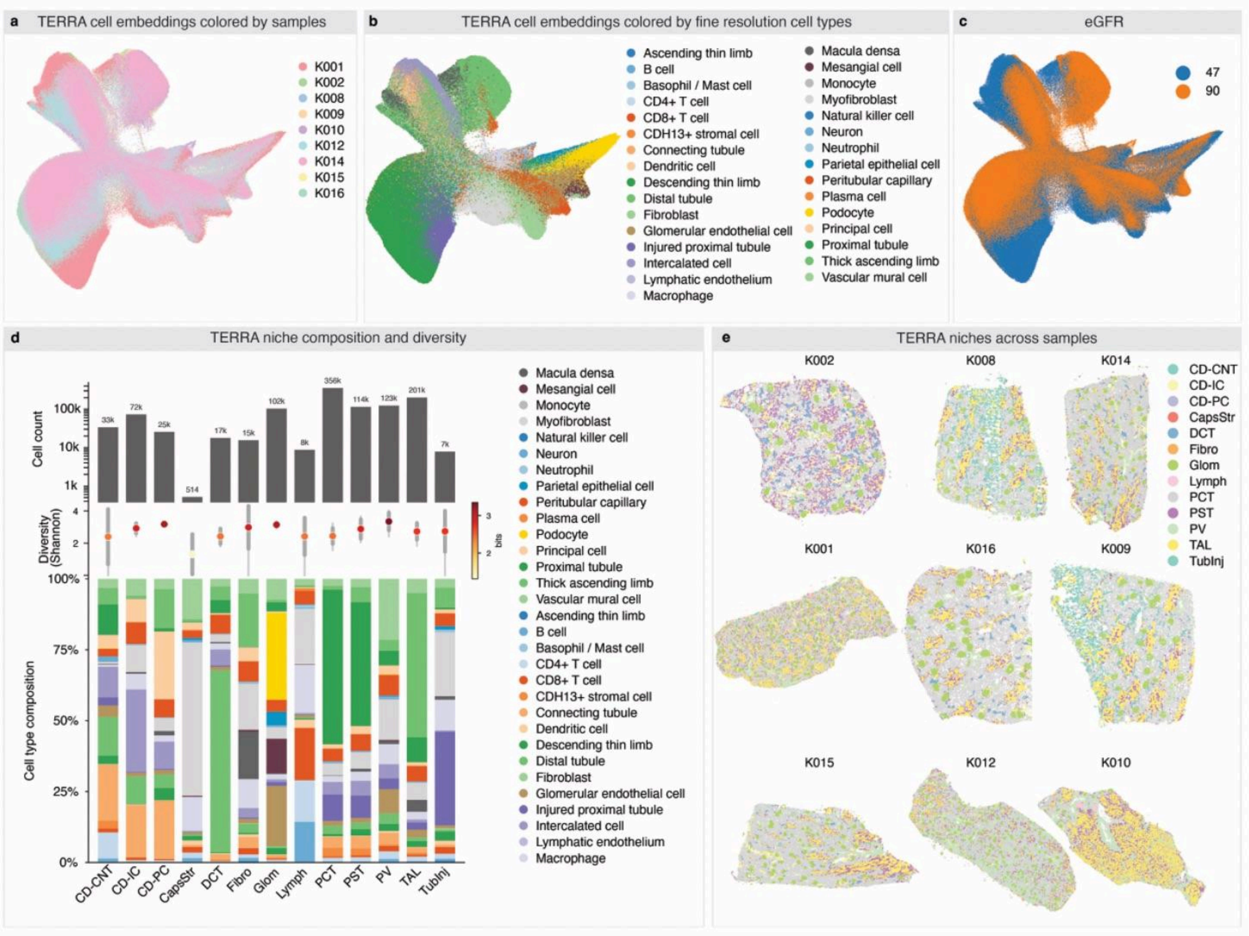


**Supplementary Fig. 17 | TERRA-based characterization of adjacent normal kidney tissue. a**, UMAP of TERRA cell embeddings colored by sample. **b**, Fine-grained cell-type annotations for the kidney projected onto the TERRA cell embedding space. **c**, Sample eGFR values mapped onto the TERRA cell embedding. **d**, Niche composition defined by cell-type identity and density, quantified via Shannon diversity scores together with total cell counts across all samples. **e**, Samples colored according to niches derived from TERRA neighborhood embeddings.

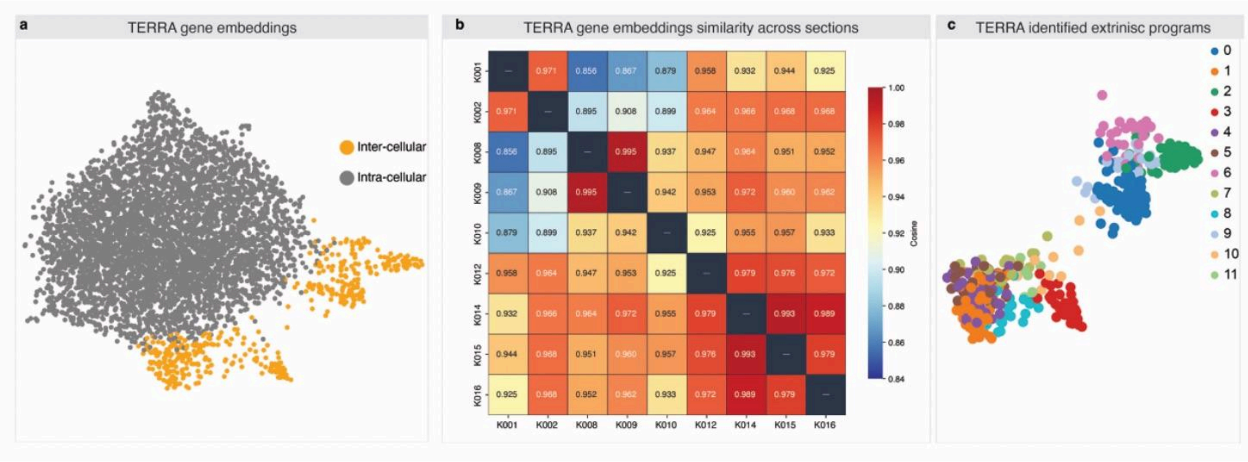


**Supplementary Fig. 18 | TERRA gene embeddings reveal coherent spatial gene programs. a**, UMAP representation of spatial gene programs within a kidney section, colored by broad functional category. **b**, Matrix plot showing cosine similarity of spatial gene embeddings across samples. **c**, Leiden clustering applied to TERRA gene embeddings of intra-cellular genes.

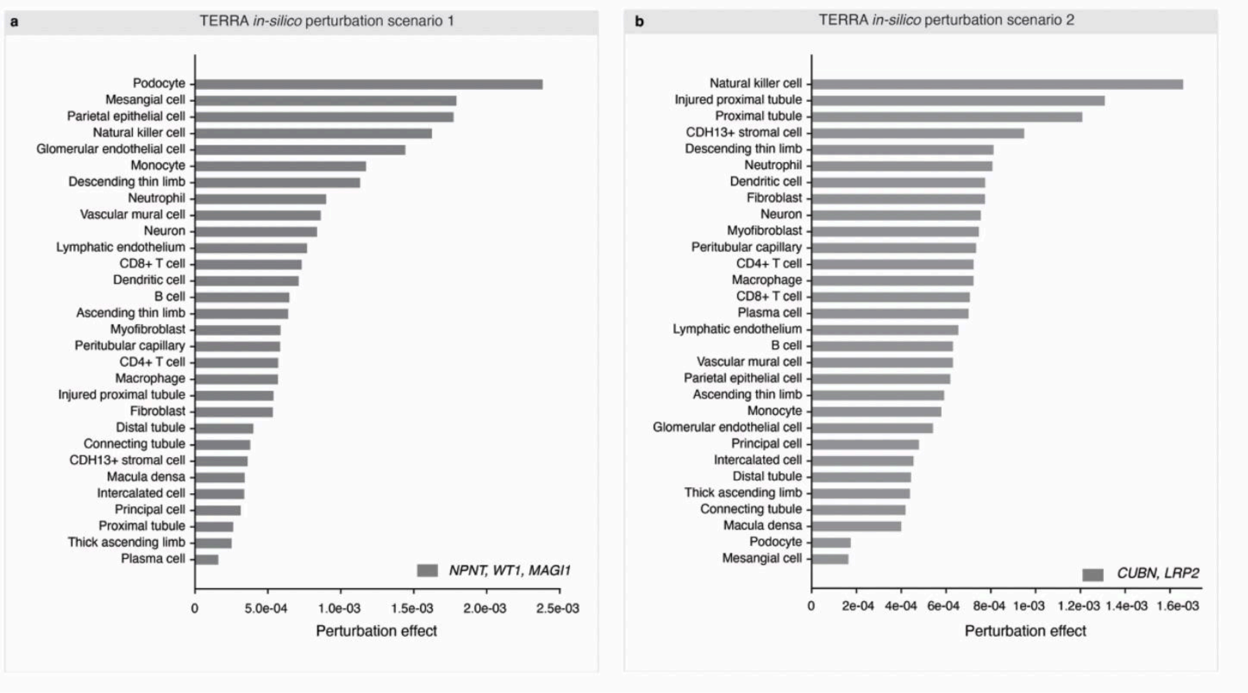


**Supplementary Fig. 19 | Perturbation effects on fine-grained cell types under scenarios 1 and 2. a**,**b**, Perturbation effects quantified as Wasserstein-2 (W2) distance between unperturbed and perturbed TERRA spatial cell embeddings (Methods).

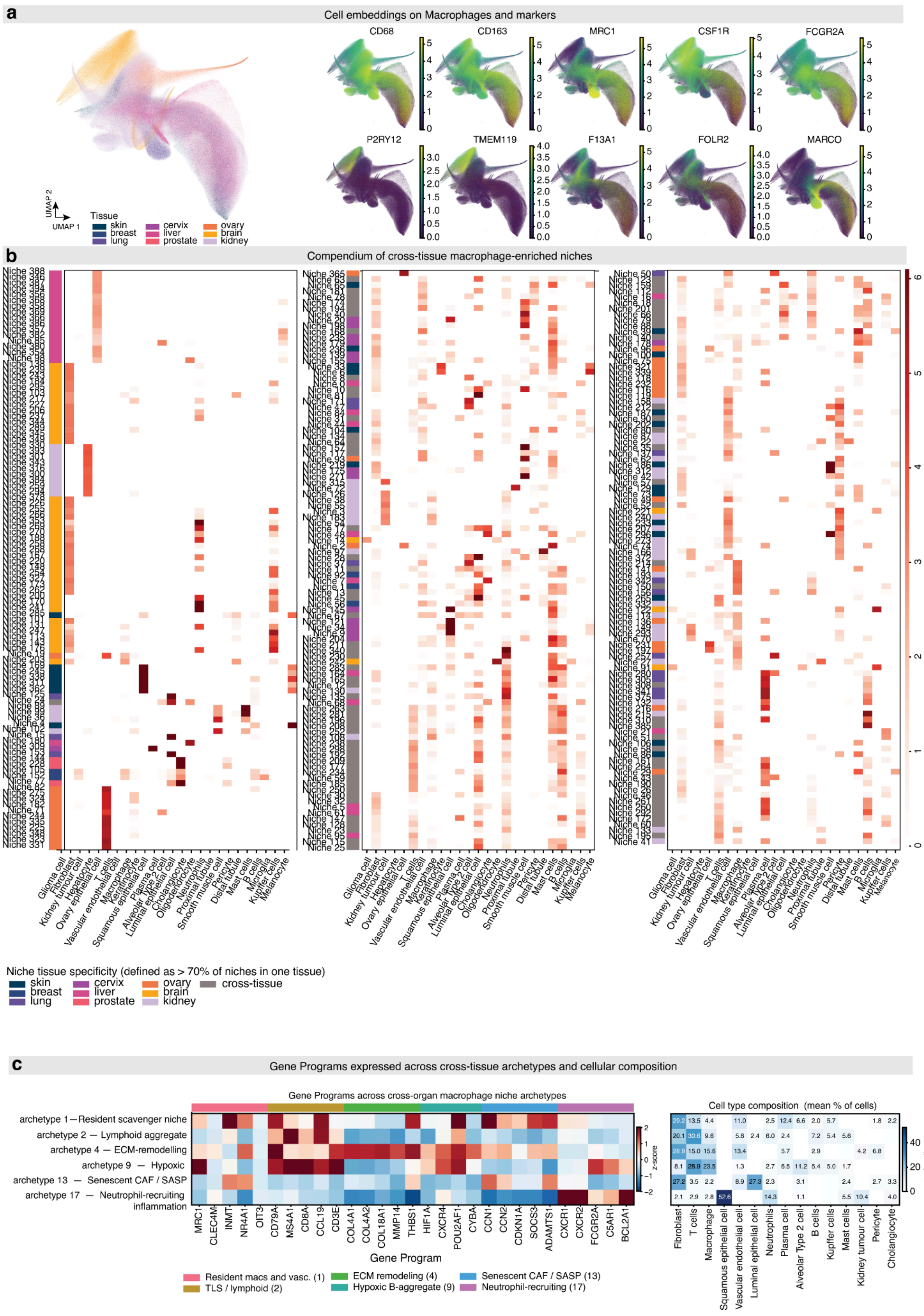
a
Cell embeddings on Macrophages and markers
CD68
CD163
MRC1
CSF1R
FCGR2A
P2RY12
TMEM119
F13A1
FOLR2
MARCO
UMAP 1
UMAP 2
Tissue
skin
breast
lung
cervix
liver
prostate
ovary
brain
kidney
b
Compendium of cross-tissue macrophage-enriched niches
log2 enrichment (obs / expected)
Glioma cell
Fibroblast
Kidney tumour cell
Hepatocyte
Ovary epithelial cell
T cells
Vascular endothelial cell
Macrophage
Keratinocyte
Squamous epithelial cell
Plasma cell
Alveolar Type 2 cell
Luminal epithelial cell
Cholangiocyte
Oligodendrocyte
Neutrophils
Proximal tubule
Smooth muscle cell
Pericyte
Distal tubule
Mast cells
B cells
Microglia
Kupffer cells
Melanocyte
Niche tissue specificity (defined as > 70% of niches in one tissue)
skin
breast
lung
cervix
liver
prostate
ovary
brain
kidney
cross-tissue
c
Gene Programs expressed across cross-tissue archetypes and cellular composition
Gene Programs across cross-organ macrophage niche archetypes
archetype 1 — Resident scavenger niche
archetype 2 — Lymphoid aggregate
archetype 4 — ECM-remodelling
archetype 9 — Hypoxic
archetype 13 — Senescent CAF / SASP
archetype 17 — Neutrophil-recruiting inflammation
MRC1
CLEC4M
INMT
NR4A1
OIT3
CD79A
MS4A1
CD8A
CCL19
CD3E
COL4A1
COL4A2
COL18A1
MMP14
THBS1
HIF1A
CXCR4
POU2AF1
CYBA
CCN1
CCN2
CDKN1A
SOCS3
ADAMTS1
CXCR1
CXCR2
FCGR2A
C5AR1
BCL2A1
Gene Program
z-score
Resident macs and vasc. (1)
TLS / lymphoid (2)
ECM remodeling (4)
Hypoxic B-aggregate (9)
Senescent CAF / SASP (13)
Neutrophil-recruiting (17)
Cell type composition (mean % of cells)
Fibroblast
T cells
Macrophage
Squamous epithelial cell
Vascular endothelial cell
Luminal epithelial cell
Neutrophils
Plasma cell
Alveolar Type 2 cell
B cells
Kupffer cells
Mast cells
Kidney tumour cell
Pericyte
Cholangiocyte
% of cells

**Supplementary Fig. 20 | Cell embedding, niche compendium and cell gene programs underlying the cross-tissue macrophage archetypes. a**, TERRA cell embedding of the 1,522,556 extracted macrophages, colored by tissue of origin (left), and the same embedding colored by normalized expression of pan-macrophage markers (*CD68*, *CD163*, *MRC1*, *CSF1R*, *FCGR2A*; top row) and tissue-resident macrophage markers (*P2RY12* and *TMEM119*, microglia; *F13A1* and *FOLR2*, metabolic-polarized; *MARCO*, alveolar; bottom row). Color scales are capped at the per-gene 99th percentile. **b**, Compendium of 298 macrophage-enriched niches, split across three panels for readability. The heatmap shows the fraction of cells of each cell type (columns) within each niche (rows). Niches are ordered by hierarchical clustering (Ward linkage on cell-type composition); the colored strip on the left of each panel marks tissue origin (≥70% of niche cells from a single tissue) or cross-tissue assignment, using the same tissue palette as in **a**. **c**, Cell gene programs (left) and mean cellular composition (right) of the six cross-organ archetypes (1, resident scavenger; 2, lymphoid aggregate; 4, ECM-remodeling; 9, hypoxic; 13, senescent CAF / SASP; 17, neutrophil-recruiting). Left, expression z-scores of representative archetype-defining genes, computed across all 16 archetypes and shown for the six cross-organ archetypes only; the colored strip above the heatmap groups genes by their associated program (key, below). Right, mean percentage of cells of each cell type per archetype.

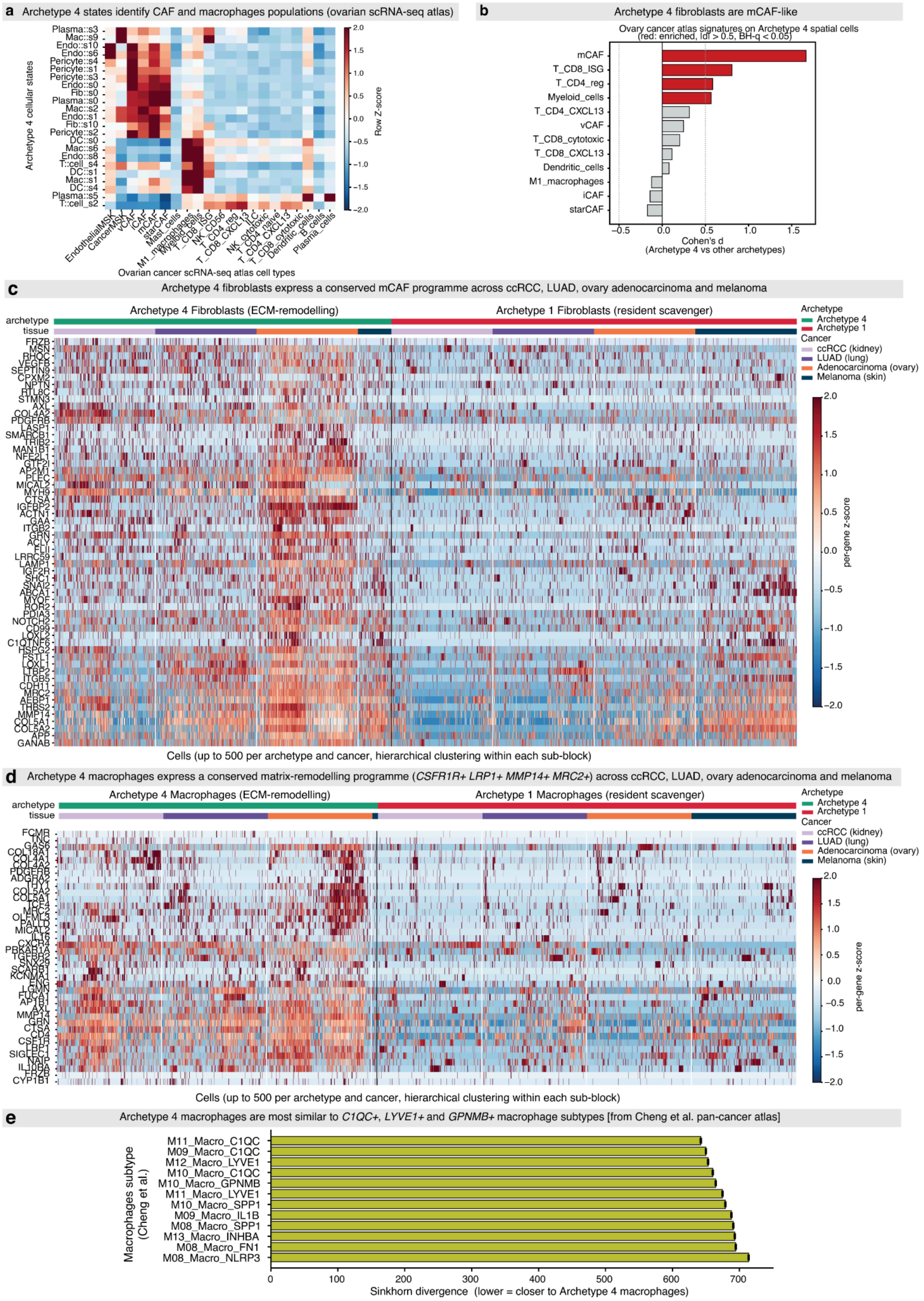
a
Archetype 4 states identify CAF and macrophages populations (ovarian scRNA-seq atlas)
Archetype 4 cellular states
Plasma::s3
Mac::s9
Endo::s10
Endo::s6
Pericyte::s4
Pericyte::s1
Pericyte::s3
Endo::s0
Fib::s0
Plasma::s0
Mac::s2
Endo::s1
Fib::s10
Pericyte::s2
DC::s0
Mac::s6
Endo::s8
T::cell_s4
DC::s1
Mac::s1
DC::s4
Plasma::s5
T::cell_s2
Row Z-score
Ovarian cancer scRNA-seq atlas cell types
b
Archetype 4 fibroblasts are mCAF-like
Ovary cancer atlas signatures on Archetype 4 spatial cells
(red: enriched, |d| > 0.5, BH-q < 0.05)
mCAF
T_CD8_ISG
T_CD4_reg
Myeloid_cells
T_CD4_CXCL13
vCAF
T_CD8_cytotoxic
T_CD8_CXCL13
Dendritic_cells
M1_macrophages
iCAF
starCAF
Cohen's d
(Archetype 4 vs other archetypes)
c
Archetype 4 fibroblasts express a conserved mCAF programme across ccRCC, LUAD, ovary adenocarcinoma and melanoma
Archetype 4 Fibroblasts (ECM-remodelling)
Archetype 1 Fibroblasts (resident scavenger)
archetype
tissue
Archetype
Archetype 4
Archetype 1
Cancer
ccRCC (kidney)
LUAD (lung)
Adenocarcinoma (ovary)
Melanoma (skin)
per-gene z-score
Cells (up to 500 per archetype and cancer, hierarchical clustering within each sub-block)
d
Archetype 4 macrophages express a conserved matrix-remodelling programme (CSFR1R+ LRP1+ MMP14+ MRC2+) across ccRCC, LUAD, ovary adenocarcinoma and melanoma
Archetype 4 Macrophages (ECM-remodelling)
Archetype 1 Macrophages (resident scavenger)
archetype
tissue
Archetype
Archetype 4
Archetype 1
Cancer
ccRCC (kidney)
LUAD (lung)
Adenocarcinoma (ovary)
Melanoma (skin)
per-gene z-score
Cells (up to 500 per archetype and cancer, hierarchical clustering within each sub-block)
e
Archetype 4 macrophages are most similar to C1QC+, LYVE1+ and GPNMB+ macrophage subtypes [from Cheng et al. pan-cancer atlas]
Macrophages subtype
(Cheng et al.)
M11_Macro_C1QC
M09_Macro_C1QC
M12_Macro_LYVE1
M10_Macro_C1QC
M10_Macro_GPNMB
M11_Macro_LYVE1
M10_Macro_SPP1
M09_Macro_IL1B
M08_Macro_SPP1
M13_Macro_INHBA
M08_Macro_FN1
M08_Macro_NLRP3
Sinkhorn divergence (lower = closer to Archetype 4 macrophages)

**Supplementary Fig. 21 | Archetype 4 niches harbor a conserved mCAF – matrix-remodeling macrophage axis across cancer indications.** **a**, Mean expression (row z-score across all Archetype 4 cellular states) of established stromal and myeloid markers from an ovarian cancer scRNA-seq atlas[1] (columns) across Archetype 4 spatial cellular states (rows). Endothelial, fibroblast and pericyte states co-cluster with CAF reference signatures (mCAF, iCAF, vCAF, starCAF); macrophage states align with myeloid and M1-macrophage references. **b**, Cohen's *d* effect sizes for the enrichment of each ovarian scRNA-seq atlas cell-type signature (Archetype 4 cells vs. all other archetypes) in spatial cells of the ovarian cancer dataset. Bars are colored red when log2 fold-change > 0.5 and Benjamini-Hochberg-adjusted $q < 0.05$; mCAF and myeloid signatures show the strongest enrichment. **c**, Per-cell z-scored expression of the 57 Archetype-4-conserved fibroblast marker genes (rows; identified by per-tissue differential expression of Archetype 4 vs. all other macrophage-enriched archetypes, retained if significant in ≥3 of 4 tissues at logFC > 0.20 and BH $q < 0.05$) across 500 randomly sampled fibroblasts per archetype × tissue × cancer condition (columns; skin melanoma sampled to availability, n = 165). Columns are clustered within each archetype × tissue × cancer sub-block by correlation distance (average linkage); gene order is fixed by global hierarchical clustering of the gene-by-cell matrix. Annotation strips above the heatmap denote archetype (top) and tissue × cancer (bottom). Sampled fibroblasts come from ccRCC (kidney), LUAD (lung), ovary adenocarcinoma and melanoma (skin) only. **d**, As in **c** but for the 39 Archetype-4-conserved macrophage marker genes (≥2 of 3 well-powered tissues; skin Mac n = 27, sampled to availability). **e**, Sinkhorn divergence[2] (mean ± SEM across 200 bootstrap iterations, 2,819 Archetype 4 macrophages vs. 1,000 atlas cells per iteration) between Archetype 4 macrophages and each macrophage sub-type of the pan-cancer tumor-infiltrating myeloid atlas[3], computed in a shared PCA-50 embedding (3,000 highly variable genes, Harmony-corrected for platform). Lower values indicate closer distributional similarity; the three closest macrophages subtypes (*M11_Macro_C1QC*, *M09_Macro_C1QC*, *M12_Macro_LYVE1*) correspond to the *C1QC+/LYVE1+* matrix-remodeling tumor macrophage states described in the original atlas.

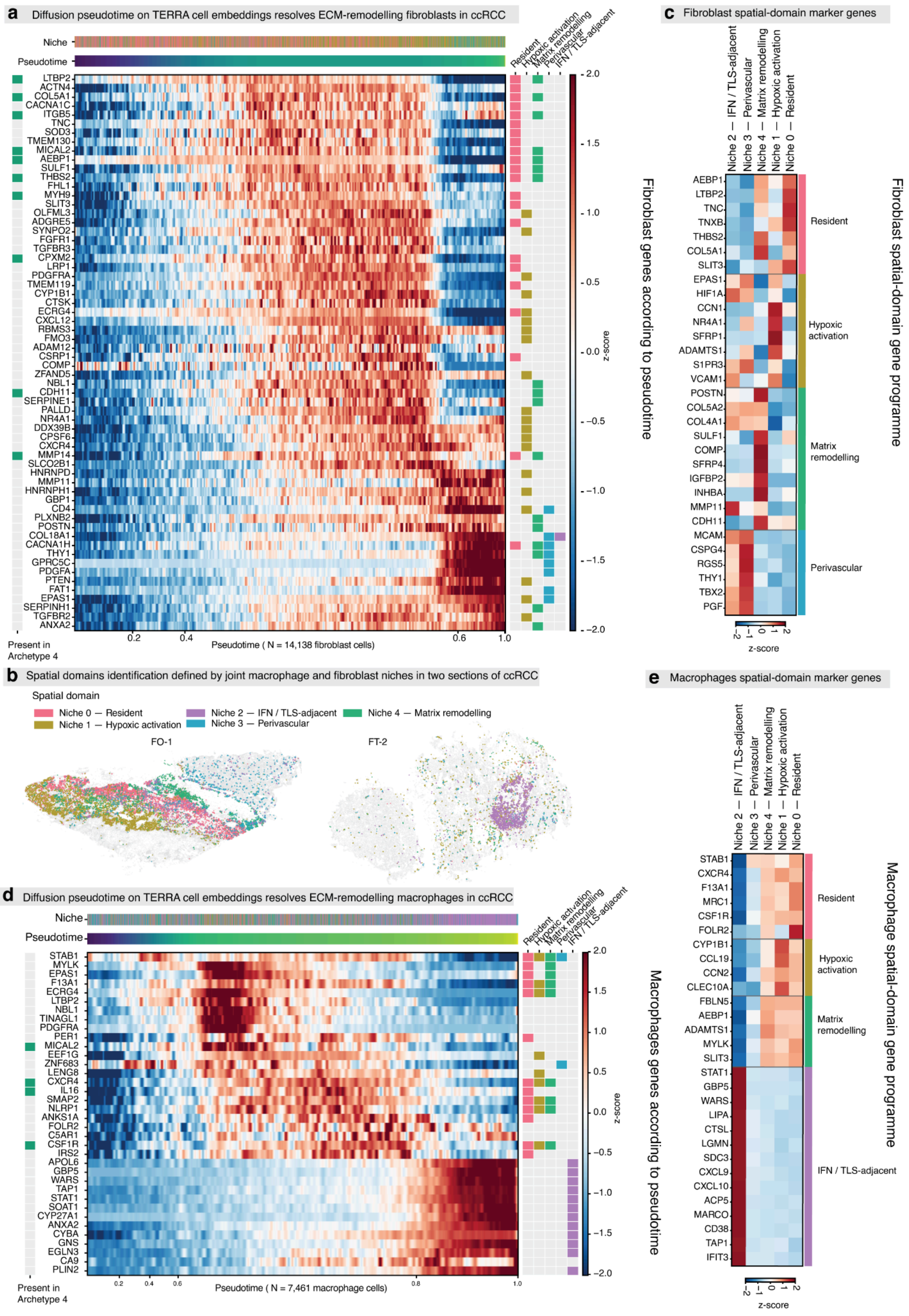
a Diffusion pseudotime on TERRA cell embeddings resolves ECM-remodelling fibroblasts in ccRCC
Niche
Pseudotime
Resident
Hypoxic activation
Matrix remodelling
Perivascular
IFN / TLS-adjacent
LTBP2
ACTN4
COL5A1
CACNA1C
ITGB5
TNC
SOD3
TMEM130
MICAL2
AEBP1
SULF1
THBS2
FHL1
MYH9
SLIT3
OLFML3
ADGRE5
SYNPO2
FGFR1
TGFBR3
CPXM2
LRP1
PDGFRA
TMEM119
CYP1B1
CTSK
ECRG4
CXCL12
RBMS3
FMO3
ADAM12
CSRP1
COMP
ZFAND5
NBL1
CDH11
SERPINE1
PALLD
NR4A1
DDX39B
CPSF6
CXCR4
MMP14
SLCO2B1
HNRNPD
MMP11
HNRNPH1
GBP1
CD4
PLXNB2
POSTN
COL18A1
CACNA1H
THY1
GPRC5C
PDGFA
PTEN
FAT1
EPAS1
SERPINH1
TGFBR2
ANXA2
Present in Archetype 4
0.2
0.4
0.6
1.0
Pseudotime ( N = 14,138 fibroblast cells)
2.0
1.5
1.0
0.5
0.0
−0.5
−1.0
−1.5
−2.0
z-score
Fibroblast genes according to pseudotime
b Spatial domains identification defined by joint macrophage and fibroblast niches in two sections of ccRCC
Spatial domain
Niche 0 — Resident
Niche 1 — Hypoxic activation
Niche 2 — IFN / TLS-adjacent
Niche 3 — Perivascular
Niche 4 — Matrix remodelling
FO-1
FT-2
c Fibroblast spatial-domain marker genes
Niche 2 — IFN / TLS-adjacent
Niche 3 — Perivascular
Niche 4 — Matrix remodelling
Niche 1 — Hypoxic activation
Niche 0 — Resident
AEBP1
LTBP2
TNC
TNXB
THBS2
COL5A1
SLIT3
EPAS1
HIF1A
CCN1
NR4A1
SFRP1
ADAMTS1
S1PR3
VCAM1
POSTN
COL5A2
COL4A1
SULF1
COMP
SFRP4
IGFBP2
INHBA
MMP11
CDH11
MCAM
CSPG4
RGS5
THY1
TBX2
PGF
Resident
Hypoxic activation
Matrix remodelling
Perivascular
−2 −1 0 1 2
z-score
Fibroblast spatial-domain gene programme
d Diffusion pseudotime on TERRA cell embeddings resolves ECM-remodelling macrophages in ccRCC
Niche
Pseudotime
Resident
Hypoxic activation
Matrix remodelling
Perivascular
IFN / TLS-adjacent
STAB1
MYLK
EPAS1
F13A1
ECRG4
LTBP2
NBL1
TINAGL1
PDGFRA
PER1
MICAL2
EEF1G
ZNF683
LENG8
CXCR4
IL16
SMAP2
NLRP1
ANKS1A
FOLR2
C5AR1
CSF1R
IRS2
APOL6
GBP5
WARS
TAP1
STAT1
SOAT1
CYP27A1
ANXA2
CYBA
GNS
EGLN3
CA9
PLIN2
Present in Archetype 4
0.2
0.4
0.6
0.8
1.0
Pseudotime ( N = 7,461 macrophage cells)
2.0
1.5
1.0
0.5
0.0
−0.5
−1.0
−1.5
−2.0
z-score
Macrophages genes according to pseudotime
e Macrophages spatial-domain marker genes
Niche 2 — IFN / TLS-adjacent
Niche 3 — Perivascular
Niche 4 — Matrix remodelling
Niche 1 — Hypoxic activation
Niche 0 — Resident
STAB1
CXCR4
F13A1
MRC1
CSF1R
FOLR2
CYP1B1
CCL19
CCN2
CLEC10A
FBLN5
AEBP1
ADAMTS1
MYLK
SLIT3
STAT1
GBP5
WARS
LIPA
CTSL
LGMN
SDC3
CXCL9
CXCL10
ACP5
MARCO
CD38
TAP1
IFIT3
Resident
Hypoxic activation
Matrix remodelling
IFN / TLS-adjacent
−2 −1 0 1 2
z-score
Macrophage spatial-domain gene programme

**Supplementary Fig. 22 | Diffusion pseudotime and joint spatial-domain analysis resolve the Archetype-4 matrix-remodeling program in fibroblasts and macrophages of two sections of ccRCC. a**, Diffusion pseudotime computed on the TERRA cell embeddings for ccRCC fibroblasts (n = 14,138), ordering cells from a resident state. The heatmap shows per-gene z-scored expression along pseudotime (columns, cells ordered by pseudotime; rows, genes). Top bars indicate the assigned spatial-domain niche and pseudotime value per cell. The right-hand strips mark, for each gene, the spatial domain in which it is a marker (Resident, Hypoxic activation, Matrix remodeling, Perivascular, IFN / TLS-adjacent) and whether it belongs to the Archetype-4 program (green squares, left). **b**, Spatial domains defined jointly from the macrophage and fibroblast niches, shown for two ccRCC sections (FO-1, FT-2). Cells are colored by spatial domain: Niche 0 (Resident), Niche 1 (Hypoxic activation), Niche 2 (IFN / TLS-adjacent), Niche 3 (Perivascular) and Niche 4 (Matrix remodeling); unassigned cells are gray. In FO-1 the matrix-remodeling domain (Niche 4) localizes to the fibrotic scar at the tumor–normal interface. **c**, Fibroblast spatial-domain gene program. Mean per-gene z-scored expression of domain-specific marker genes (rows) across the five spatial domains (columns), grouped by the domain in which each gene is most highly expressed (Resident, Hypoxic activation, Matrix remodeling, Perivascular). **d**, Diffusion pseudotime computed on the TERRA cell embeddings for ccRCC macrophages (n = 7,461), ordering cells from a resident state. The heatmap shows per-gene z-scored expression along pseudotime (columns, cells ordered by pseudotime; rows, genes). Top bars indicate the assigned spatial-domain niche and pseudotime value per cell. The right-hand strips mark, for each gene, the spatial domain in which it is a marker (Resident, Hypoxic activation, Matrix remodeling, IFN / TLS-adjacent) and whether it belongs to the Archetype-4 program (green squares, left). **e**, Macrophage spatial-domain gene program. Mean per-gene z-scored expression of domain-specific marker genes (rows) across the five spatial domains (columns), grouped by the domain in which each gene is most highly expressed (Resident, Hypoxic activation, Matrix remodeling, IFN / TLS-adjacent).

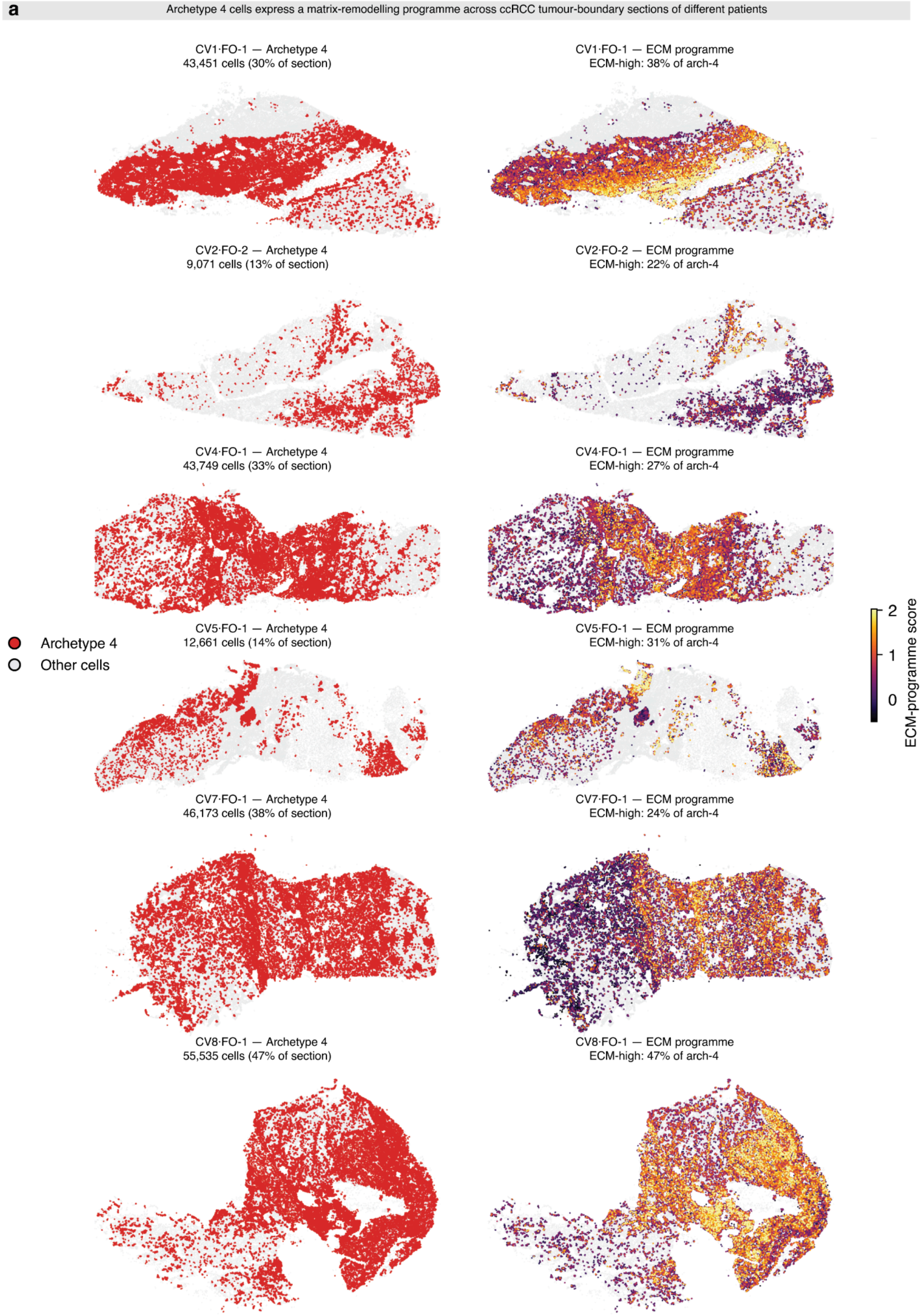
a
Archetype 4 cells express a matrix-remodelling programme across ccRCC tumour-boundary sections of different patients
CV1·FO-1 — Archetype 4
43,451 cells (30% of section)
CV1·FO-1 — ECM programme
ECM-high: 38% of arch-4
CV2·FO-2 — Archetype 4
9,071 cells (13% of section)
CV2·FO-2 — ECM programme
ECM-high: 22% of arch-4
CV4·FO-1 — Archetype 4
43,749 cells (33% of section)
CV4·FO-1 — ECM programme
ECM-high: 27% of arch-4
CV5·FO-1 — Archetype 4
12,661 cells (14% of section)
CV5·FO-1 — ECM programme
ECM-high: 31% of arch-4
CV7·FO-1 — Archetype 4
46,173 cells (38% of section)
CV7·FO-1 — ECM programme
ECM-high: 24% of arch-4
CV8·FO-1 — Archetype 4
55,535 cells (47% of section)
CV8·FO-1 — ECM programme
ECM-high: 47% of arch-4
Archetype 4
Other cells
ECM-programme score
2
1
0

**Supplementary Fig. 23 | Archetype 4 cells express a matrix-remodeling program across ccRCC tumor-boundary sections from different patients. a**, Paired spatial maps of six ccRCC tumor-boundary Xenium sections, each from a different patient (CV1, CV2, CV4, CV5, CV7, CV8; all pre-treatment, tumor-boundary sampling). In each row, the left panel shows all cells of the section (gray) with archetype 4 cells highlighted in red; the number and percentage of archetype 4 cells in the section are indicated. The right panel shows the same archetype 4 cells colored by their matrix-remodeling (ECM) program score, a per-cell signature score computed over the archetype 4 ECM program genes (*POSTN*, *COL5A2*, *COL4A1*, *COL5A1*, *SULF1*, *COMP*, *SFRP4*, *IGFBP2*, *INHBA*, *MMP11*, *MMP14*, *CDH11*, *MRC2*, *LRP1*); remaining cells are shown in gray. The color scale (dark to yellow) is shared across all sections. "ECM-high" denotes the percentage of a section's archetype 4 cells scoring above the cohort-wide 66th-percentile threshold of the ECM score. Archetype 4 cells were present and expressed the matrix-remodeling program in all six sections, with the magnitude and spatial distribution of ECM-program expression varying between patients.

### Supplementary Notes

**Supplementary Note 1 | Design ablations on mouse brain data**

To establish TERRA's design and to benchmark it against existing approaches under controlled conditions, we used a paired mouse brain benchmark in which MERFISH[4] and STARmap[5] profile matched coronal sections, with ground-truth tissue regions annotated against the Allen Mouse Brain Reference Atlas (P56, coronal). Models were pretrained on the full assay-specific gene panels (MERFISH: 1,085 genes; STARmap: 1,015 genes) and evaluated on the 431 genes shared between the two assays (Supplementary Fig. 3). Niche identification was quantified by normalized mutual information (NMI) and adjusted Rand index (ARI) against the reference annotations, and cross-assay integration by the integration local inverse Simpson's index (iLISI; higher indicates better mixing) and maximum mean discrepancy (MMD; lower is better).

Tokenization and architecture. We swept the principal design axes with all other settings held at the default configuration (Supplementary Fig. 4). A cell-graph tokenizer with combined (rank + value) gene encoding gave the best niche identification and the strongest integration; per-cell sequence lengths of 32–64 tokens were optimal (at L = 32 and L = 64, respectively, 60.1% and 95.3% of MERFISH cells and 37.3% and 83.1% of STARmap cells are fully represented without truncation on the shared gene panel (Supplementary Fig. 3)), whereas longer sequences degraded performance as sequences became dominated by low-expression genes; a continuous MLP value encoding outperformed discretized value embeddings at all dimensions; and rank-based spatial (segment) encoding substantially outperformed coordinate-based encoding: coordinate encoding attained artifactually high iLISI by aligning cells in coordinate space rather than learning biologically meaningful structure. A masking ratio of 0.6 was optimal for niche identification, with lower ratios producing high run-to-run variance, consistent with the prediction task becoming too easy.

Normalization and the batch metatoken. Because the combined tokenizer has separate count and rank components, we compared normalization applied to either component or both (Supplementary Fig. 5; Supplementary Note 2). Raw values and shifted-log normalization ranked among the top strategies for niche identification, whereas explicit normalizations such as Seurat V3 and analytic Pearson residuals degraded it; Seurat V3 achieved the best integration despite poor niche identification. These results indicate that rank-based tokenization already provides an implicit, scale-free normalization, so additional explicit normalization is unnecessary and can be counterproductive. Adding a batch metatoken enabled integration across the two assays without any explicit batch-correction objective: with the metatoken, MERFISH and STARmap were mixed in embedding space while anatomically coherent niches were preserved, whereas removing it collapsed cross-assay integration and degraded niche identification (Supplementary Fig. 6). At inference the metatoken is zeroed (padded), so the model produces assay-agnostic embeddings.

**Supplementary Note 2 | Normalization strategies**

TERRA's combined tokenizer represents each gene token by both its within-cell expression rank and its value. Normalization can therefore be applied independently to two streams — the rank stream (which determines how genes are ordered within a cell) and the value stream (which sets the value token) — and within each stream at up to three composable levels: a cell-level depth normalization, a gene-level per-gene scaling, and a count-level within-cell transform. The three categories in Supplementary Fig. 5 reflect where normalization is applied: unnormalized ranks (only the value/count stream normalized; gene order left on raw counts), unnormalized counts (only the rank stream normalized), and full normalization (both streams). Within each category, the labeled strategies are compositions of the building blocks below; for example, "Gene Corr. RD NZ Mean" applies gene-corrected read-depth normalization at the cell level followed by non-zero-mean scaling at the gene level, and "RD Shifted Log" applies read-depth normalization followed by a shifted-log transform.

No normalization. Raw integer counts are used directly. Because the tokenizer ranks genes by expression within each cell, raw counts already yield a well-defined, library-size-invariant ordering.

Cell-level (depth) normalization. Read depth (RD) divides each cell's counts by its total count and rescales to a fixed target depth (10,000), i.e. counts-per-10k. Gene-corrected read depth (Gene Corr. RD) is the same per-cell rescaling but with a target depth that increases with the size of the gene panel (target = $a + b \cdot N\_genes$, with $a = 153.48$ and $b = 0.0487$ fit by linear regression across the corpus), so that the expected depth is comparable across assays with different panel sizes.

Gene-level (per-gene) scaling. Each gene is divided by a corpus-wide per-gene factor loaded from precomputed normalization factors. Mean uses the per-gene mean expression across all cells; non-zero mean (NZ Mean) uses the per-gene mean over only the cells in which the gene is detected, reducing the influence of sparsity. When a cell-level depth normalization is also applied, the gene factor is the corresponding statistic computed on the depth-normalized data (e.g. gene-corrected-read-depth non-zero mean), so the two levels compose consistently.

Count-level (within-cell) transforms. Shifted log applies $\log(1 + x)$ (natural log; scanpy.pp.log1p). Analytic Pearson residuals compute negative-binomial residuals under an offset model with shared overdispersion ($\theta = 100$), $(x - \mu)/\sqrt{\mu + \mu^2/\theta}$ with $\mu$ the expected count from the cell- and gene-marginals, clipped to $\pm\sqrt{n\_obs}$[6]; this is a standalone transform and is not combined with cell- or gene-level normalization. Seurat V3 variance-stabilizing normalization centers each gene by its mean and scales by the expected standard deviation obtained from a LOESS fit (span 0.3, degree 2) of log-variance against log-mean, applied per batch (clipping omitted)[7].

Constraints and rationale. Shifted-log and other strictly monotone within-cell transforms preserve the within-cell gene ordering, so on the rank stream they leave the ranking unchanged and act only on the value token; they are therefore most consequential for the value/count stream. Analytic Pearson residuals and Seurat V3, which recenter and rescale across cells, can change the ranking and are the strategies that most strongly alter the representation. Across the sweep (Supplementary Fig. 5), leaving the rank stream on raw counts (with at most a shifted-log value token) gave the best niche identification, whereas explicit cross-cell normalizations (Seurat V3, Pearson residuals) degraded it: Seurat V3 achieved the strongest cross-assay integration but the weakest niche identification. These results indicate that rank-based tokenization already provides an implicit, library-size-robust normalization, so additional explicit normalization is unnecessary and can be counterproductive; TERRA therefore uses raw within-cell ranks with a shifted-log value token by default.

**Supplementary Note 3 | Cross-assay benchmarking on mouse brain data**

Using the mouse brain benchmark described in Supplementary Note 1, we compared TERRA against task-specific spatial methods that are fit directly on the data (neighborhood gene-expression PCA, NicheCompass, CellCharter, BANKSY with Harmony, and GraphST with PASTE) and against zero-shot foundation models applied without fine-tuning (Novae, Nicheformer, CellPLM, scGPT-spatial). To enable a fair comparison, we trained TERRA on the 431 shared genes for this comparison. For every method, neighborhood-level representations were clustered into 30 niches and evaluated both for agreement with Allen Reference Atlas annotations (NMI and ARI) and for cross-assay integration between MERFISH and STARmap (iLISI, higher is better; MMD, lower is better) (Supplementary Figs. 7, 8).

TERRA was the only method to achieve top-tier niche identification and strong cross-assay integration simultaneously. It attained the highest niche-identification scores overall, statistically indistinguishable from the strongest task-specific method, NicheCompass, and significantly better than every other task-specific method and every zero-shot foundation-model baseline. Critically, TERRA integrated MERFISH and STARmap (high iLISI, low MMD) without any explicit batch correction or alignment, whereas NicheCompass integrated the two assays far less effectively. TERRA therefore improved on NicheCompass overall, combining equivalent niche identification with substantially stronger integration. The only methods to exceed TERRA's integration did so through procedures that forcibly integrate samples — post-hoc embedding alignment (BANKSY with Harmony) or prior spatial-coordinate alignment (GraphST with PASTE) — conferring an inherent advantage on integration metrics while degrading niche identification and precluding native integration of heterogeneous samples. The zero-shot foundation models, by contrast, largely failed to integrate the two assays, and where they showed apparent mixing they did so by collapsing biological structure, as reflected in their poor niche-identification scores.

### Supplementary Tables

Supplementary Tables 1-4 are provided as separate machine-readable files (Microsoft Excel, one table per file). Supplementary Table 5 is presented below.

**Supplementary Table 1 | Fetal pancreas spatial gene programs (SGPs) and gene-enrichment analysis.** Provided as a separate Excel file with two sheets. The first sheet lists the 53 spatial gene programs (SGP1–SGP53) identified by Leiden clustering of TERRA neighborhood-gene embeddings across the fetal pancreas epithelial niches, one column per SGP giving its constituent genes (Methods). The second sheet contains the gene-enrichment analysis underlying the heatmap in Fig. 3l (GSEApy; Methods): rows are enriched terms and columns are the SGPs shown in Fig. 3l, with each value the $-\log_{10}$ adjusted P value of that term for that SGP.

**Supplementary Table 2 | Fetal pancreas capillary subpopulation marker genes.** Provided as a separate Excel file with one sheet per fetal capillary subpopulation (acinar-associated (AA), general G1, G2 and G3, and islet-associated (IA)). Each sheet lists the differentially expressed marker genes of that subpopulation together with the associated test statistics (Wilcoxon rank-sum test; Methods).

**Supplementary Table 3 | Fetal pancreas cell–cell communication results.** Provided as a separate Excel file (single sheet) containing ligand–receptor interactions inferred with CellPhoneDB v5 (1,000 permutations; minimum expression 5%; Methods) across cell types and developmental stages. Columns give the interaction identifier and interacting pair, the interacting partners and genes, interaction metadata (secreted, receptor, directionality, classification), the sending and receiving populations (each stratified by developmental stage), the interaction score and the permutation P value.

**Supplementary Table 4 | Xenium custom gene panel for fetal and adult pancreas.** Provided as a separate Excel file with two sheets (fetal and adult pancreas). Each lists the genes of the custom 100-gene add-on panel (10x Genomics, 1000766) used alongside the Xenium 5,000-gene panel (10x Genomics, 1000724), with gene symbol, Ensembl ID and probe count.

| | TERRA-112M | TERRA-96M |
|---|---|---|
| Pretraining cells / split | 112M (full corpus) | 96M (training split; 215 sections held out) |
| Encoder | 12 layers, dim 384, 6 heads, MLP ×4 | 12 layers, dim 384, 6 heads, MLP ×4 |
| Predictor | 12 layers, dim 192, 6 heads, MLP ×4 | 12 layers, dim 192, 6 heads, MLP ×4 |

| | | |
|---|---|---|
| Parameters (encoder / predictor) | ~30.7M / ~5.6M | ~30.7M / ~5.6M |
| Gene vocabulary | 23,407 tokens | 23,407 tokens |
| Sequence (index / neighborhood / total) | 256 / 2,560 / 2,816 | 256 / 2,560 / 2,816 |
| Neighbors (k) | 10 | 10 |
| Count normalization | shifted-log | shifted-log |
| Value encoding | 20-bin soft (value-bins) | continuous MLP |
| Positional encoding | segment (cell-rank) | coordinate (relative x/y) |
| Masking (block; per-block ratio) | 0.6 | 0.8 |
| Batch size (per GPU) | 224 | 288 |
| Optimizer | AdamW | AdamW |
| LR schedule | warmup→cosine, $1\times10^{-6} \rightarrow 6\times10^{-5} \rightarrow 1\times10^{-6}$ | constant $2\times10^{-5}$ |
| Weight-decay schedule | 0.4 → 4.0 (cosine) | 0.04 → 0.4 (cosine) |
| EMA momentum | 0.9995 → 1.0 | 0.9995 → 1.0 |
| Precision / attention | bfloat16 / flash | bfloat16 / flash |
| Hardware | 16× H100 (DDP) | 16× H100 (DDP) |

**Supplementary Table 5 | TERRA-112M and TERRA-96M hyperparameters.** Model, data and training hyperparameters for the two released models (Methods). Relationship between the design ablations and the released configurations. The ablations (Supplementary Figs. 4–6) characterize each design axis in isolation on the controlled mouse-brain benchmark; the released models' configurations (Supplementary Table 5) were established during model development for the large, multi-tissue human corpus. The primary model, TERRA-112M, is consistent with the ablation optima on most axes, including rank-based (segment) spatial encoding and a masking ratio of 0.6. The principal difference is the per-cell sequence length: the released models use 256 tokens per cell rather than the 32–64 optimal on the low-plex mouse panels (~1,000 genes, mostly <64 expressed per cell), because the human analyses use the Xenium 5K panel, whose higher per-cell gene counts require a longer sequence to avoid truncation (Supplementary Fig. 1d). Because the mouse-brain benchmark differs from the deployment setting in species, tissue diversity and gene-panel plexity, the mouse-brain optima are not expected to transfer exactly; the remaining differences (e.g. TERRA-96M's coordinate spatial encoding and higher masking ratio) reflect development-stage choices in that earlier, held-out model.
